# Suboptimality in Perceptual Decision Making

**DOI:** 10.1101/060194

**Authors:** Dobromir Rahnev, Rachel N. Denison

## Abstract

**Short Abstract:** Human perceptual decisions are often described as optimal, but this view remains controversial. To elucidate the issue, we review the vast literature on suboptimalities in perceptual tasks and compile the proposed hypotheses about the origins of suboptimal behavior. Further, we argue that general claims about optimality are virtually meaningless and result in a false sense of progress. Instead, real progress can be achieved by building observer models that account for both optimal and suboptimal behavior. To achieve such progress, the field should focus on assessing the hypotheses about suboptimal behavior compiled here and stop chasing optimality.

**Long Abstract:** Human perceptual decisions are often described as optimal. Critics of this view have argued that claims of optimality are overly flexible and lack explanatory power. Meanwhile, advocates for optimality have countered that such criticisms single out a few selected papers. To elucidate the issue of optimality in perceptual decision making, we review the extensive literature on suboptimal performance in perceptual tasks. We discuss eight different classes of suboptimal perceptual decisions, including improper placement, maintenance, and adjustment of perceptual criteria, inadequate tradeoff between speed and accuracy, inappropriate confidence ratings, misweightings in cue combination, and findings related to various perceptual illusions and biases. In addition, we discuss conceptual shortcomings of a focus on optimality, such as definitional difficulties and the limited value of optimality claims in and of themselves. We therefore advocate that the field drop its emphasis on whether observed behavior is optimal and instead concentrate on building and testing detailed observer models that explain behavior across a wide range of tasks. To facilitate this transition, we compile the proposed hypotheses regarding the origins of suboptimal perceptual decisions reviewed here. We argue that verifying, rejecting, and expanding these explanations for suboptimal behavior – rather than assessing optimality per se – should be among the major goals of the science of perceptual decision making.

## 1. INTRODUCTION

How do people make perceptual judgments based on the available sensory information? This fundamental question has been a focus of psychological research from the 19th century on (Fechner 1860; Helmholtz 1856). Many perceptual tasks naturally lend themselves to what has traditionally been called “ideal observer” analysis, whereby the optimal behavior is mathematically determined given a set of assumptions such as the presence of sensory noise, and human behavior is compared to this standard (Geisler 2011; Green and Swets 1966; Ulehla 1966). The extensive literature on this topic includes many examples of humans performing similarly to an ideal observer but also many examples of suboptimal behavior. Perceptual science has a strong tradition of developing models and theories that attempt to account for the full range of empirical data on how humans perceive (Macmillan and Creelman 2005).

Recent years have seen an impressive surge of Bayesian theories of human cognition and perception (Gershman, Horvitz, and Tenenbaum 2015; Griffiths, Lieder, and Goodman 2015; Tenenbaum et al. 2011). These theories often depict humans as optimal decision-makers, especially in the area of perception. A number of high-profile papers have shown examples of human perceptual behavior that is close to optimal (Ernst and Banks 2002; Körding and Wolpert 2004; Landy et al. 1995; Shen and Ma 2016), while other papers have attempted to explain apparently suboptimal behaviors as being in fact optimal (Weiss, Simoncelli, and Adelson 2002). Consequently, many statements by researchers in the field leave the impression that humans are essentially optimal in perceptual tasks:

> “… psychophysics is providing a growing body of evidence that human perceptual computations are ‘Bayes’ optimal’.” (Knill & Pouget, 2004, p. 712)

> “Across a wide range of tasks, people seem to act in a manner consistent with optimal Bayesian models” (Vul, Goodman, Griffiths, & Tenenbaum, 2014, p.1)
>
> “These studies with different approaches have shown that human perception is close to the Bayesian optimal” (Körding & Wolpert, 2006, p. 321)

Despite a number of recent criticisms of such assertions regarding human optimality (Bowers and Davis 2012a, 2012b; Eberhardt and Danks 2011; Jones and Love 2011; Marcus and Davis 2013, 2015), as well as statements from some of the most prominent Bayesian theorists that their goal is not to demonstrate optimality (Goodman et al. 2015; Griffiths et al. 2012), the quotes above indicate that the view that humans are (close to) optimal when making perceptual decisions has taken strong foothold.

The main purpose of this paper is to counteract assertions about human optimality by bringing together the extensive literature on suboptimal perceptual decision making. While the description of the many findings of suboptimality will occupy a large part of the paper, we do not advocate for a shift of labeling observers from “optimal” to “suboptimal.” Instead, we will ultimately argue that we should abandon any emphasis on optimality or suboptimality and return to building a science of perception that attempts to account for all types of behavior.

The paper is organized into six sections. After introducing the topic (Section 1), we explain the Bayesian approach to perceptual decision making and explicitly define a set of Standard Assumptions that typically determine what behavior is considered optimal (Section 2). In the central section of the paper, we review the vast literature of suboptimal perceptual decision making and show that suboptimalities have been reported in virtually every class of perceptual tasks (Section 3). We then discuss theoretical problems with the current narrow focus on optimality such as difficulties in defining what is truly optimal and the limited value of optimality claims in and of themselves (Section 4). Finally, we argue that the way forward is to build observer models that give equal emphasis to all components of perceptual decision making, not only the decision rule (Section 5). We conclude that the field should abandon its emphasis on optimality and instead focus on thoroughly testing the hypotheses that have already been generated (Section 6).

## 2. DEFINING OPTIMALITY

Optimality can be defined within a large number of frameworks. Here we adopt a Bayesian approach, because it is both widely used in the field and general: other approaches to optimality can often be expressed in Bayesian terms.

### 2.1 The Bayesian approach to perceptual decision making

The Bayesian approach to perceptual decision making starts with specifying the generative model of the task. The model defines the sets of world states, or stimuli, *S*, internal responses *X*, actions *A*, and relevant parameters Θ (such as the sensitivity of the observer). We will mostly focus on cases in which two possible stimuli *s*_1_ and *s*_2_ are presented, and the possible “actions” *a*_1_ and *a*_2_ consist of reporting that the corresponding stimulus was shown. The Bayesian approach then specifies the following quantities (see Figure 1 for a graphical depiction):

**Figure 1.**
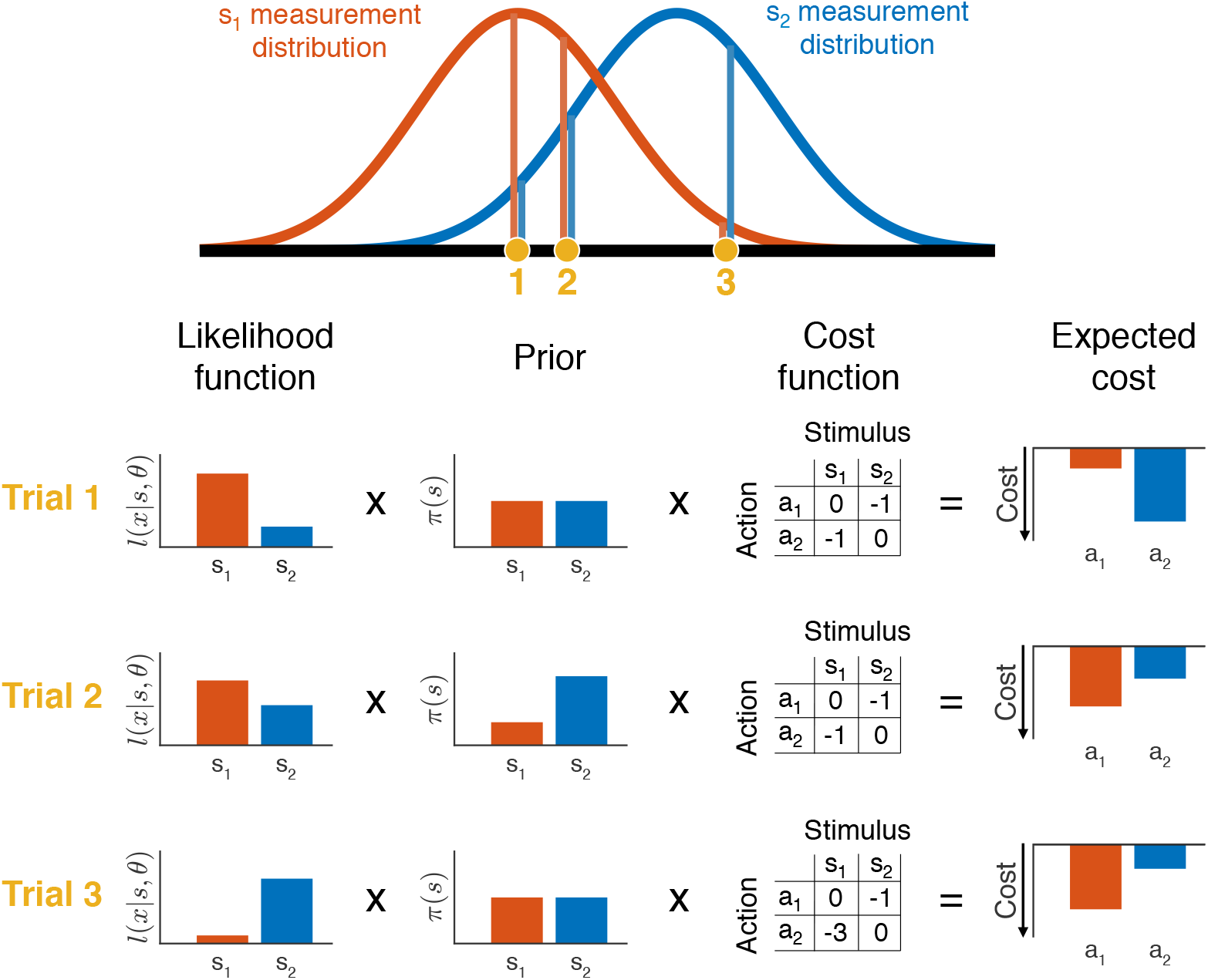
Graphical depiction of Bayesian inference. An observer is deciding between two possible stimuli – *s*_11_ (e.g., leftward motion) and *s*_21_ (e.g., rightward motion) – which produce Gaussian measurement distributions of internal responses. The observer’s internal response varies from trial to trial, depicted by the three yellow circles for three example trials. On a given trial, the likelihood function is the height of each of the two measurement densities at the value of the observed internal response (lines drawn from each yellow circle) – i.e., the likelihood of that internal response given each stimulus. For illustration, a different experimenter-provided prior and cost function is assumed on each trial. The action *a_i_* corresponds to choosing stimulus *s_i_*. We obtain the expected cost of each action by multiplying the likelihood, prior, and cost corresponding to each stimulus, and then summing the costs associated with the two possible stimuli. The optimal decision rule is to choose the action with the lower cost (equivalent to choosing the bar with less negative values). In trial 1, the prior and cost function are unbiased, so the optimal decision depends only on the likelihood function. In trial 2, the prior is biased toward *s*_21_, making a_2_ the optimal choice even though *s*_11_ is slightly more likely. In trial 3, the cost function favors a_1_, but the much higher likelihood of *s*_21_ makes a_2_ the optimal choice.

#### Likelihood function

An external stimulus can produce a range of internal responses. The measurement density *p*(*x*|*s*, θ) is the probability density of obtaining an internal response *x* given a particular stimulus *s*. The likelihood function, *l*(*s*|*x, θ*) is equal to the measurement density but is defined for a fixed internal response as opposed to a fixed stimulus.

#### Prior

The prior *π*(*s*) describes one’s assumptions about the probability of each stimulus *s*.

#### Cost function

The cost function *L*(*s*,*a*) (also called loss function) specifies the cost of taking a specific action for a specific stimulus.

#### Decision rule

The decision rule *δ*(*x*) indicates under what combination of the other quantities you should perform one action or another.

We refer to the likelihood function, prior, cost function, and decision rule as the LPCD components of perceptual decision making. According to Bayesian Decision Theory (Körding and Wolpert 2006; Maloney and Mamassian 2009), the optimal decision rule is the one that minimizes the expected loss, over all possible stimuli, for the chosen action *a*. After applying Bayes’ theorem, we obtain the optimal decision rule as a function of the likelihood, prior, and cost function:

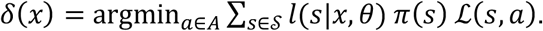

### 2.2 Standard Assumptions

Determining whether observers’ decisions are optimal requires the specification of the four LPCD components. How do researchers determine the quantitative form of each component? Below we present a typical set of Standard Assumptions related to each LPCD component.

#### Likelihood function assumptions

The Standard Assumptions here include Gaussian measurement distributions and stimulus encoding that is independent from other factors such as stimulus presentation history. Note that the experimenter derives the likelihood function from the assumed measurement distribution.

#### Prior and cost function assumptions

The Standard Assumption about observers’ internal representations of the prior and cost function is that they are identical to the experimentally-defined quantities. Unless specifically mentioned, experiments reviewed below present *s*_1_ and *s*_2_ equally often, which is equivalent to a uniform prior (e.g., 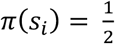 for two stimuli), and expect observers to maximize percent correct, which is equivalent to a cost function that punishes all incorrect responses, and rewards all correct responses, equally.

#### Decision rule assumptions

The Standard Assumption about the decision rule is that it is identical to the optimal decision rule.

Finally, additional general Standard Assumptions include expectations that observers can perform the proper computations on the LPCD components. Note that as specified, the Standard Assumptions consider the Gaussian variability at encoding as the sole corrupting element for perceptual decisions. Section 3 assembles the evidence against this claim.

The attentive reader may object that the Standard Assumptions cannot be universally correct. For example, assumptions related to the likelihood function are likely false for specific paradigms (e.g., measurement noise may not be Gaussian), while assumptions about observers adopting the experimentally-defined prior and cost function are likely false for complex experimental designs (Beck et al. 2012). Nevertheless, we take the Standard Assumptions as a useful starting point for our review since, explicitly or implicitly, they are assumed in most (although not all) studies. In Section 3 we label all deviations from behavior prescribed by the Standard Assumptions as examples of suboptimality. We discuss alternative ways of defining optimality in Section 4 and ultimately argue that general statements about the optimality or suboptimality of perceptual decisions are meaningless.

## 3. REVIEW OF SUBOPTIMALITY IN PERCEPTUAL DECISION MAKING

We review eight categories of tasks for which the optimal decision rule can be determined. For each task category, we first note any relevant information about the measurement distribution, prior, or cost function. We then plot the measurement distributions together with the optimal decision rule (which we depict as a criterion drawn on the internal responses X). We then review specific suboptimalities within each task category. For each explanation of apparently suboptimal behavior below, we indicate the LPCD component proposed to have been violated using the notation [***LPCD component***, such as [***decision rule***]. Note that violations of the assumed measurement distributions result in violations of the assumed likelihood functions. In some cases, the suboptimalities have been attributed to issues that apply to multiple components (indicated as [***general***]) or issues of methodology (indicated as [***methodological***]).

### 3.1 Criterion in 2-choice tasks

In the most common case, observers must distinguish between two possible stimuli, *s*_1_ and *s*_2_, presented with equal probability and associated with equal reward. In Figure 2 we plot the measurement distributions and optimal criteria for the cases of equal and unequal internal variability. The criterion used to make the decision corresponds to the decision rule.

**Figure 2.**
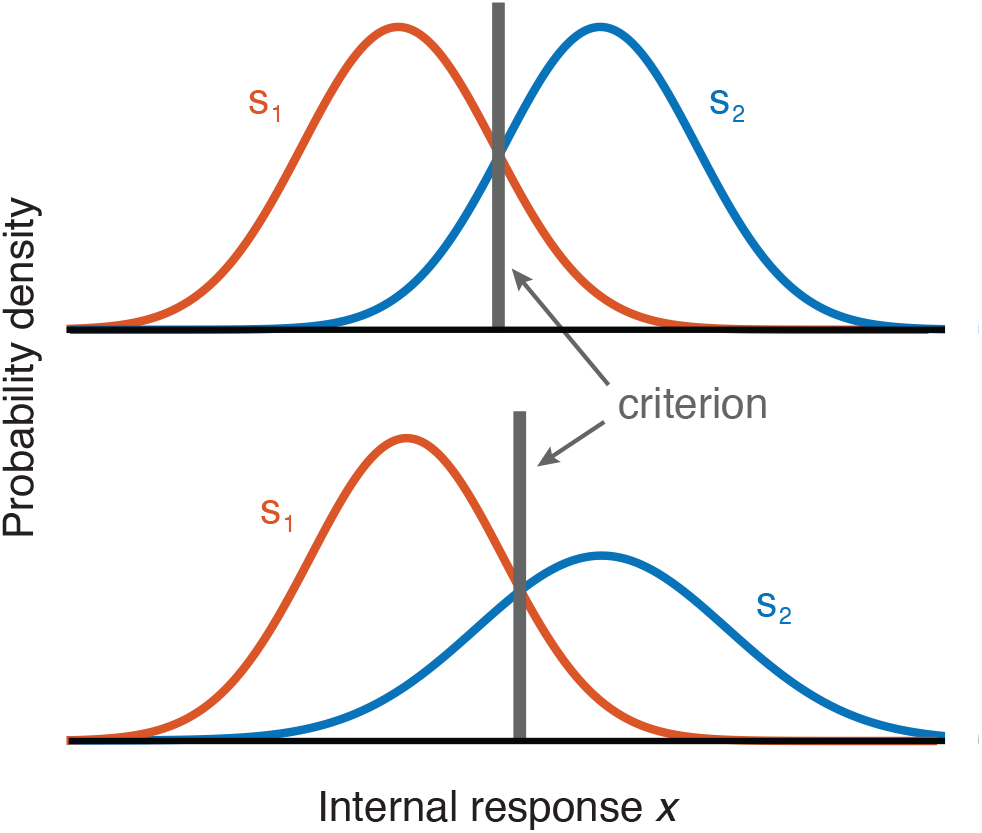
Depiction of the measurement distributions (colored curves) and optimal criteria (equivalent to the decision rules) in 2-choice tasks. The upper panel depicts the case when the two stimuli produce the same internal variability (*σ*_1_ = *σ*_2_). The gray vertical line represents the location of the optimal criterion. The lower panel shows the location of the optimal criterion when the variability of the two measurement distributions differs (*σ*_1_ < *σ*_2_, in which case the optimal criterion results in a higher proportion of *s*_11_ responses).

#### 3.1.1 Detection criteria

Many tasks involve the simple distinction between noise (*s*_1_) and signal+noise (*s*_2_). These are usually referred to as detection tasks. In most cases *s*_1_ is found to produce smaller internal variability than *s*_2_ (Green and Swets 1966; Macmillan and Creelman 2005; Swets, Tanner, and Birdsall 1961), from where it follows that an optimal observer would choose *s*_1_ more often than *s*_2_ even when the two stimuli are presented at equal rates (Figure 2). Indeed, many detection studies find that observers choose the noise distribution *s*_1_ more than half of the time (Gorea & Sagi, 2000; Green & Swets, 1966; Rahnev et al., 2011; Reckless et al., 2014; Solovey, Graney, & Lau, 2015; Swets et al., 1961). However, most studies do not allow for the estimation of the exact measurement distributions for individual observers and thus it is an open question how optimal observers in those studies actually are. A few studies have reported conditions in which observers choose the noise stimulus *s*_1_ *less* than half of the time (Morales et al., 2015; Rahnev et al., 2011; Solovey et al., 2015). Assuming that the noise distributions in those studies also had lower variability, such behavior is likely suboptimal.

#### 3.1.2 Discrimination criteria

Detection tasks require observers to distinguish between the noise vs. signal+noise stimuli but other tasks require observers to discriminate between two roughly equivalent stimuli. For example, observers might discriminate left vs. rightward motion or clockwise vs. counterclockwise grating orientation. For these types of stimuli, the measurement distributions for each stimulus category can be safely assumed to have approximately equal variability (Macmillan and Creelman 2005; See et al. 1997). Such studies find that the average criterion location across the whole group of observers is usually close to the optimal but individual observers can still exhibit substantial biases (e.g., Whiteley & Sahani, 2008). In other words, what appears as an optimal criterion on average (across observers) may be an average of suboptimal criteria (Mozer, Pashler, and Homaei 2008; Vul et al. 2014). This issue can appear within an individual observer, too, with suboptimal criteria on different trials averaging out to resemble an optimal criterion (see Section 3.2). To check for criterion optimality within individual observers, we re-analyzed the data from a recent study in which observers discriminated between a grating tilted 45 degrees clockwise or counterclockwise from vertical (Rahnev et al. 2016). Seventeen observers came for four sessions on different days completing 480 trials each time. Using a binomial test, we found that 57 of the 68 total sessions exhibited significant deviation from unbiased responding. Further, observers tended to have relatively stable biases as demonstrated by a positive criterion correlation across all pairs of sessions (all p’s < .003). Thus, even if the performance of the group appears to be close to optimal, individual observers often deviate substantially from optimality.

#### 3.1.3 Two-stimulus tasks

The biases observed in detection and discrimination experiments led to the development of the two-alternative forced-choice (2AFC) task, in which both stimulus categories are presented on each trial, in order to reduce individual bias (Macmillan and Creelman 2005). 2AFC tasks separate the two stimuli either temporally (also referred to as 2-interval-forced-choice or 2IFC tasks) or spatially. Note that in recent years researchers have begun to use the term “2AFC” for tasks in which only one stimulus is presented. To avoid confusion, we adopt the term “2-stimulus tasks” to refer to tasks where two stimuli are presented (the original meaning of 2AFC) and the term “1-stimulus tasks” to refer to tasks like single-stimulus detection and discrimination (e.g., the tasks discussed in 3.1.1 and 3.1.2).

Even though 2-stimulus tasks were designed to remove observer bias, significant biases have been observed for them, too. While biases in spatial 2AFC tasks have received less attention, several suboptimalities have been documented for 2IFC tasks. For example, early research suggested that the second stimulus is more often selected as the one of higher intensity, a phenomenon called *time-order errors* (Fechner 1860; Osgood 1953). More recently Yeshurun et al. (2008) re-analyzed 2IFC data from seventeen previous experiments and found significant interval biases. The direction of the bias varied between the different experiments, suggesting that the specific experimental design has an influence on observers’ bias.

#### 3.1.4 Explaining suboptimality in 2-choice tasks

Why do people appear to have trouble setting appropriate criteria in 2-choice tasks? One possibility is that they have a tendency to give the same fixed response when uncertain [***decision rule***]. For example, a given observer may respond that they saw left (rather than right) motion every time they get distracted or had very low evidence for either choice. This could be due to a preference for one of the two stimuli or one of the two motor responses. Re-analysis of another previous study (Rahnev, Lau, and De Lange 2011), where we withheld the stimulus-response mapping until after the stimulus presentation, found that 12 of the 21 observers still showed a significant response bias for motion direction. Thus, a preference in motor behavior cannot fully account for this type of suboptimality.

Another possibility is that for many observers even ostensibly “equivalent” stimuli such as left and right motion give rise to measurement distributions with unequal variance [***likelihood function***]. In that case, an optimal decision rule would produce behavior that appears biased. Similarly, in 2-stimulus tasks, it is possible that the two stimuli are not given the same resources or that the internal representations for each stimulus are not independent of each other [***likelihood function***]. Finally, in the case of detection tasks, it is possible that some observers employ an idiosyncratic cost function by treating misses as less problematic than false alarms since the latter can be interpreted as lying [***cost function***].

### 3.2 Maintaining stable criteria

So far, we considered the optimality of the decision rule when all trials are considered together. We now turn our attention to whether observers’ behavior varies across trials or conditions (Figure 3).

**Figure 3.**
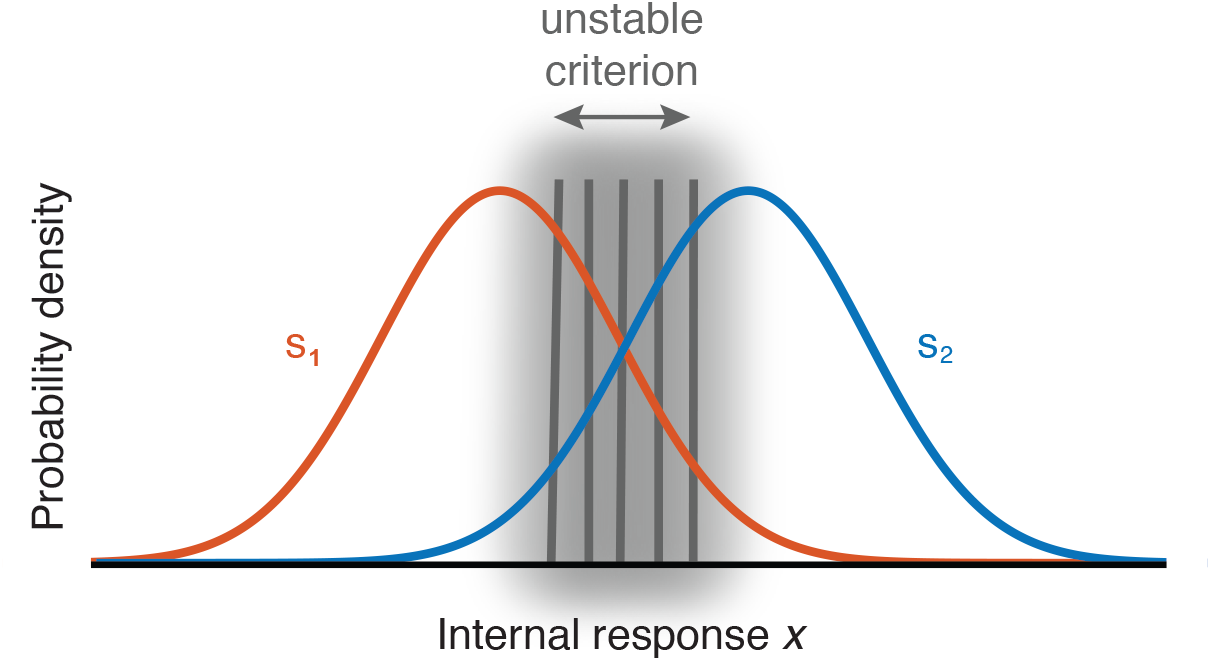
Depiction of a failure to maintain a stable criterion. The optimal criterion is shown in Figure 2 but observers often fail to maintain that criterion over the course of the experiment, resulting in a criterion that effectively varies over trials. Colored curves show measurement distributions.

#### 3.2.1 Sequential effects

Optimality in laboratory tasks requires that judgments are made based on the evidence from the current stimulus independently of previous stimuli. However, sequential effects are ubiquitous in perceptual tasks (Fischer and Whitney 2014; Fründ, Wichmann, and Macke 2014; Kaneko and Sakai 2015; Liberman, Fischer, and Whitney 2014; Norton et al. 2017; Tanner, Haller, and Atkinson 1967; Treisman and Faulkner 1984; Ward and Lockhead 1970; Yu and Cohen 2009). The general finding is that observers’ responses are positively autocorrelated such that the response on the current trial is likely to be the same as on the previous trial, though in some cases negative autocorrelations have also been reported (Tanner et al. 1967; Ward and Lockhead 1970). Further, observers are able to adjust to new trial-to-trial statistics but this adjustment is only strong in the direction of default biases and weak in the opposite direction (Abrahamyan et al. 2016). Similar effects have been observed in other species such as mice (Busse et al. 2011).

#### 3.2.2 Criterion attraction

Interleaving trials that require different criteria also hinders optimal criterion placement. Gorea & Sagi (2000) demonstrated that when high-contrast stimuli (optimally requiring a relatively liberal detection criterion) and low-contrast stimuli (optimally requiring a relatively conservative detection criterion) were presented simultaneously, observers tended to use the same compromised detection criterion that was suboptimal for both the high- and low-contrast stimuli. Observers used a suboptimal criterion despite the fact that on each trial they knew with 100% certainty which contrasts might have been present in each location. Similar criterion attraction has been shown in a variety of paradigms that involved using stimuli of different contrasts (Gorea, Caetta, and Sagi 2005; Gorea and Sagi 2001, 2002; Zak et al. 2012), attended vs. unattended stimuli (Morales et al. 2015; Rahnev, Maniscalco, et al. 2011), and central vs. peripheral stimuli (Solovey et al. 2015).

#### 3.2.3 Irrelevant reward influencing the criterion

The optimal decision rule is insensitive to multiplicative changes to the cost function. For example, rewarding all correct responses with $0.01 vs. $0.03, while wrong answers receive $0, should not alter the decision criterion; in both cases the optimal decision rule is the one that maximizes percent correct. However, both greater monetary rewards and punishments lead observers to adopt a more liberal detection criterion such that more stimuli are identified as targets (Reckless et al. 2013, 2014). Similar changes to the response criterion due to monetary motivation are obtained in a variety of paradigms (Henriques, Glowacki, and Davidson 1994; Taylor et al. 2004). To complicate matters, observers’ personality traits interact with the type of monetary reward in altering response criteria (Markman, Baldwin, and Maddox 2005).

#### 3.2.4 Explaining suboptimality in maintaining stable criteria

Why do people appear to shift their response criteria based on factors that should be irrelevant for the criterion placement? Sequential effects are typically explained in terms of an automatic tendency to exploit the continuity in our normal environment, even though such continuity is not present in most experimental setups (Fischer and Whitney 2014; Fritsche, Mostert, and de Lange 2017; Liberman et al. 2014). The visual system could have built-in mechanisms that bias new representations towards recent ones [***likelihood function***] or it may assume that a new stimulus is likely to be similar to a recent one [***prior***]. (Note that the alternative likelihoods or priors would need to be defined over pairs or sequences of trials.) Adopting a prior that the environment is autocorrelated may be a good strategy for maximizing reward: environments typically are autocorrelated and if they are not, such a prior may not hurt performance (Yu and Cohen 2009).

Criterion attraction likely stems from an inability to maintain two separate criteria simultaneously. This is equivalent to asserting that in certain situations observers cannot maintain a more complicated decision rule (e.g., different criteria for different conditions) and instead use a simpler one (e.g., single criterion for all conditions) [***decision rule***]. It is harder to explain why personality traits or task features such as increased monetary rewards (that should be irrelevant to the response criterion) change observers’ criteria.

### 3.3 Adjusting choice criteria

One of the most common ways to assess optimality in perceptual decision making is to manipulate the prior probability of the stimulus classes and/or provide unequal payoffs that bias responses towards one of the stimulus categories (Macmillan and Creelman 2005). Manipulating prior probabilities affects the prior *π*(*s*), while manipulating payoffs affects the cost function *L*(*s, a*). However, the two manipulations have an equivalent effect on the optimal decision rule: both require observers to shift their decision criterion by a factor dictated by the specific prior probability or reward structure (Figure 4).

**Figure 4.**
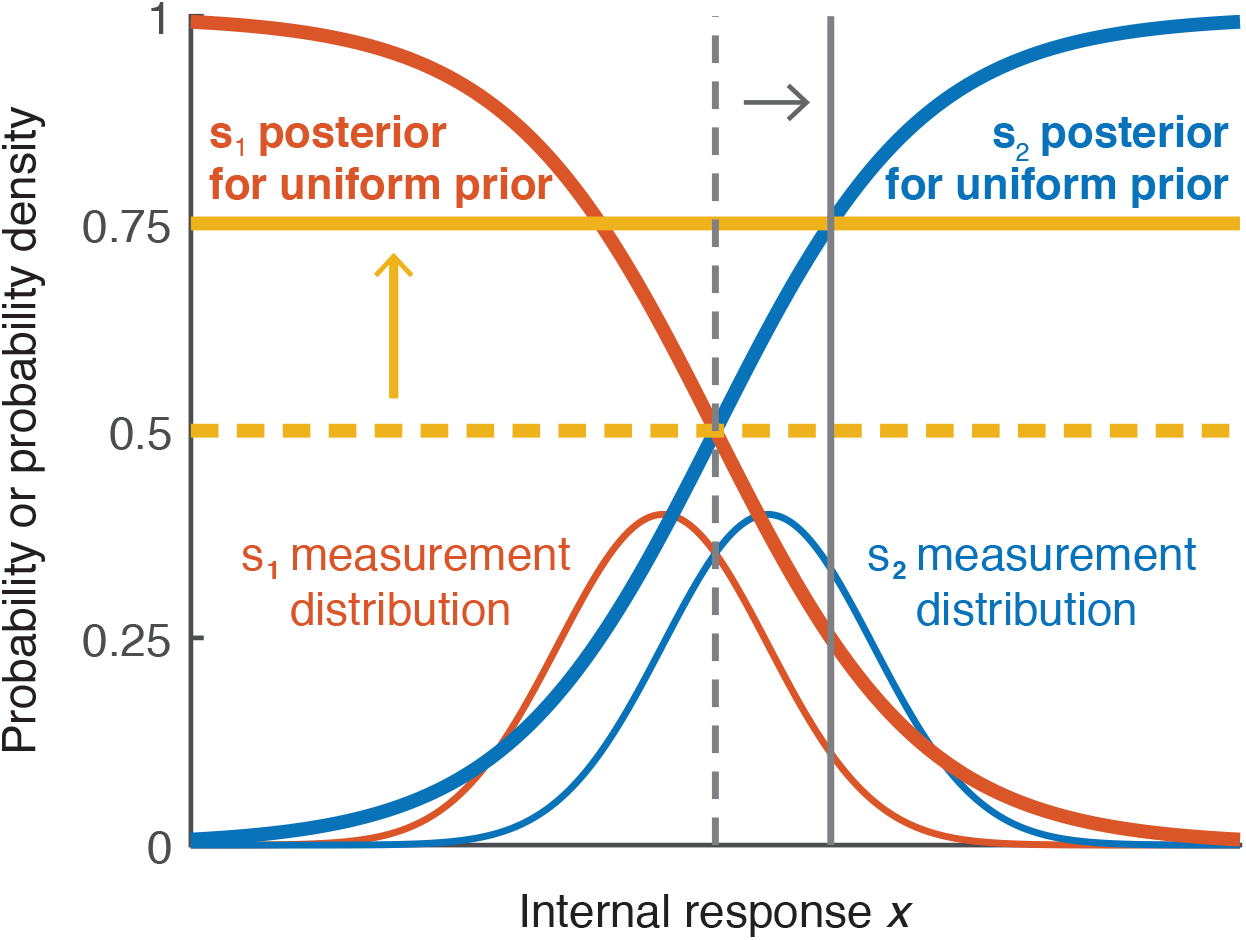
Depiction of optimal adjustment of choice criteria. In addition to the *s*_1_ and *s*_2_ measurement distributions (in thin red and blue lines), the figure shows the corresponding posterior probabilities as a function of x assuming uniform prior (in thick red and blue lines). The vertical criteria depict optimal criterion locations on x (thin gray lines) and correspond to the horizontal thresholds (thick yellow lines). Optimal criterion and threshold for equal prior probabilities and payoffs are shown in dashed lines. If unequal prior probability or unequal payoff is provided such that *s*_1_ ought to be chosen three times as often as *s*_2_, then the threshold would optimally be shifted to 0.75, corresponding to a shift in the criterion such that the horizontal threshold and vertical criterion intersect on the *s*_2_ posterior probability function. The y-axis is probability density for the measurement distributions, and probability for the posterior probability functions (the y-axis ticks refer to the posterior probability).

#### 3.3.1 Priors

Two main approaches have been used to determine whether observers can optimally adjust their criterion when one of two stimuli has a higher probability of occurrence. In base-rate manipulations, long blocks of the same occurrence frequency are used, and observers are typically not informed of the probabilities of occurrence in advance (e.g., Maddox, 1995). Most studies find that observers adjust their criterion to account for the unequal base rate, but this adjustment is smaller than what is required for optimal performance, resulting in a conservative criterion placement (Bohil and Maddox 2003a; Green and Swets 1966; Maddox and Bohil 2001, 2003, 2005; Maddox, Bohil, and Dodd 2003; Maddox and Dodd 2001; Tanner 1956; Tanner et al. 1967; Vincent 2011). Some studies have suggested that observers become progressively more suboptimal as the base-rate becomes progressively more extreme (Bohil and Maddox 2003a; Green and Swets 1966). However, a few studies have reported that certain conditions result in extreme criterion placement such that observers rely more on base-rate information than is optimal (Maddox and Bohil 1998b).

A second way to manipulate the probability of occurrence is to do it on a trial-by-trial basis and explicitly inform the observers about the stimulus probabilities before each trial. This approach also leads to conservative criterion placement such that observers do not shift their criterion enough (Ackermann and Landy 2015; de Lange et al. 2013; Rahnev, Lau, et al. 2011; Summerfield and Koechlin 2010; Ulehla 1966).

#### 3.3.2 Payoffs

The decision criterion can also be manipulated by giving different payoffs for different responses. The general finding with this manipulation is that observers, again, do not adjust their criterion enough (Ackermann and Landy 2015; Bohil and Maddox 2001, 2003a, 2003b; Busemeyer and Myung 1992; Maddox and Bohil 1998a, 2000, 2001, 2003, 2005; Maddox et al. 2003; Maddox and Dodd 2001; Markman et al. 2005; Taylor et al. 2004; Ulehla 1966) and, as with base rates, become more suboptimal for more extreme payoffs (Bohil and Maddox 2003a). Nevertheless, one study that involved a very large number of sessions with two monkeys reported extreme criterion changes (Feng et al. 2009).

Criterion adjustments in response to unequal payoffs are usually found to be more suboptimal compared to adjustments in response to unequal base-rates (Ackermann and Landy 2015; Bohil and Maddox 2001, 2003a; Busemeyer and Myung 1992; Healy and Kubovy 1981; Maddox 2002; Maddox and Bohil 1998a; Maddox and Dodd 2001) though the opposite pattern was found by Green and Swets (1966).

Finally, the exact payoff structure may also influence observers’ optimality. For instance, introducing a cost for incorrect answers leads to more suboptimal criterion placement compared to conditions with the same optimal criterion shift but without a cost for incorrect answers (Maddox and Bohil 2000; Maddox et al. 2003; Maddox and Dodd 2001).

#### 3.3.3 Explaining suboptimality in adjusting choice criteria

Why do people appear not to adjust their decision criteria optimally in response to priors and rewards? One possibility is that they do not have an accurate internal representation of the relevant probability implied by the prior or reward structure [***general***] (Acerbi, Vijayakumar, and Wolpert 2014; Ackermann and Landy 2015; Zhang and Maloney 2012). For example, Zhang & Maloney (2012) argued for the presence of “ubiquitous log odds” that systematically distort people’s probability judgments such that small values are overestimated while large values are underestimated (Brooke and MacRae 1977; Juslin, Nilsson, and Winman 2009; Kahneman and Tversky 1979; Varey, Mellers, and Birnbaum 1990).

A possible explanation for the suboptimality in base-rate experiments is the “flat-maxima” hypothesis, according to which the observer adjusts the decision criterion based on the change in reward and has trouble finding its optimal value if other criterion positions result in similar reward rates [***methodological***] (Bohil and Maddox 2003b; Busemeyer and Myung 1992; Maddox and Bohil 2001, 2003, 2004, 2005; Maddox et al. 2003; Maddox and Dodd 2001; von Winterfeldt and Edwards 1982). Another possibility is that the prior observers adopt in base-rate experiments comes from a separate process of Bayesian inference. If observers are uncertain about the true base rate, a prior assumption that it is likely to be unbiased would result in insufficient base-rate adjustment [***methodological***]. A central tendency bias can also arise when observers form a prior based on the sample of stimuli they have encountered so far, which are unlikely to cover the full range of the experimenter-defined stimulus distribution (Petzschner and Glasauer 2011). We classify these issues as methodological since if the observers have not been able to learn a particular LPC component, then they cannot adopt the optimal decision rule.

Finally, another possibility is that observers also place a premium on being correct rather than just maximizing reward [***cost function***]. Maddox & Bohil (1998a) posited the COmpetition Between Reward and Accuracy maximization (COBRA) hypothesis according to which observers attempt to maximize reward but also place a premium on accuracy (Maddox and Bohil 2004, 2005). This consideration applies to manipulations of payoffs but not of prior probabilities, and may explain why payoff manipulations typically lead to larger deviations from optimality than priors.

### 3.4 Tradeoff between speed and accuracy

In the examples above, the only variable of interest has been observers’ choice irrespective of their reaction times (RTs). However, if instructed, observers can provide responses faster at lower accuracy, a phenomenon known as speed-accuracy tradeoff (SAT; Fitts, 1966; Heitz, 2014). An important question here is whether observers can adjust their RTs optimally to achieve maximum reward in a given amount of time (Figure 5). A practical difficulty for studies attempting to address this question is that the accuracy/RT curve is not generally known and is likely to differ substantially between different tasks (Heitz 2014). Therefore, the only Standard Assumption here is that accuracy increases monotonically as a function of RT. Precise accuracy/RT curves can be constructed by assuming one of the many models from the sequential sampling modeling framework (Forstmann, Ratcliff, and Wagenmakers 2016), and there is a vibrant discussion about the optimal stopping rule depending on whether signal reliability is known or unknown (Bogacz 2007; Bogacz et al. 2006; Drugowitsch et al. 2012, 2015; Hanks et al. 2011; Hawkins et al. 2015; Thura et al. 2012). However, since different models predict different accuracy/RT curves, in what follows we only assume a monotonic relationship between accuracy and RT.

**Figure 5.**
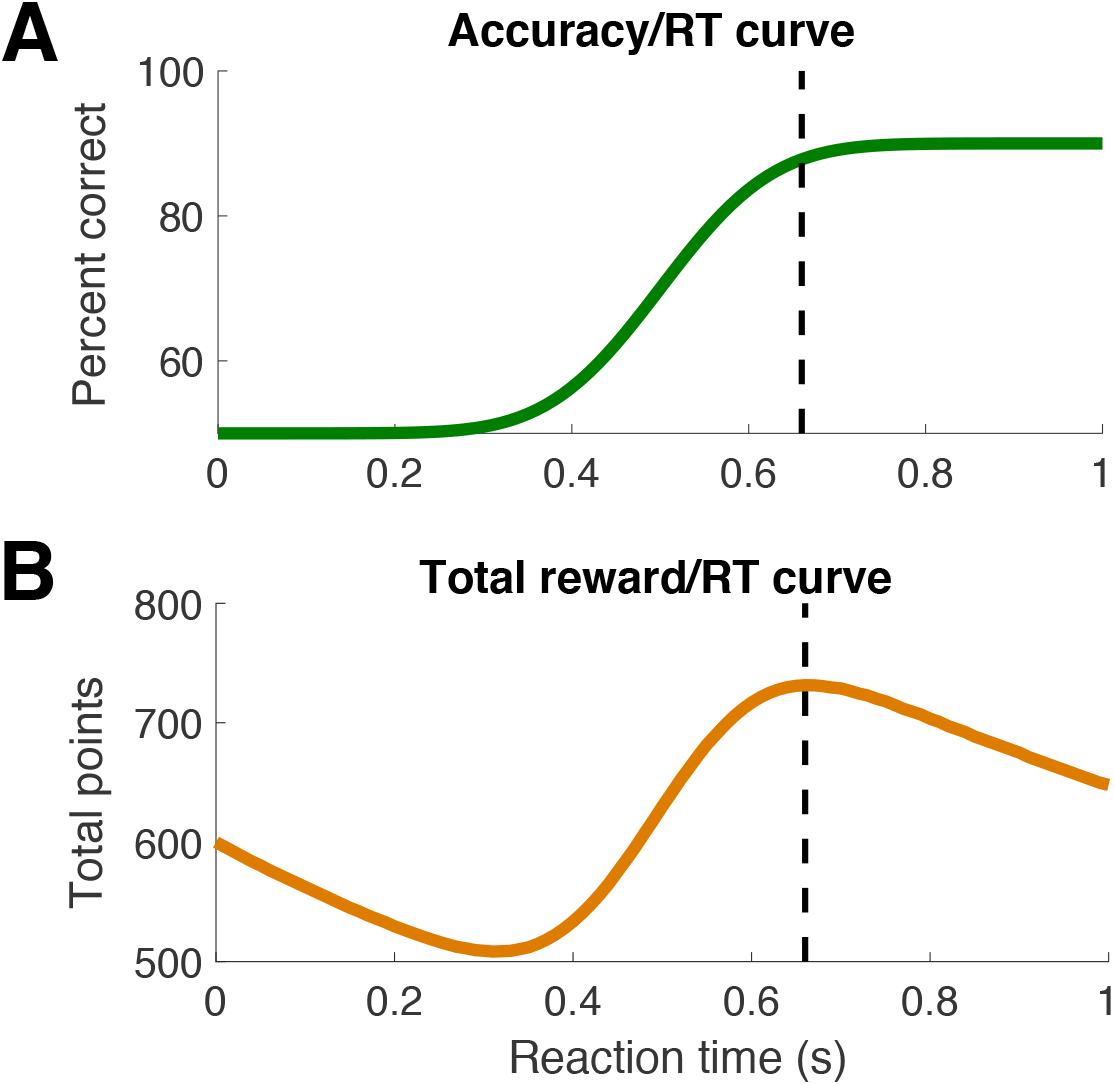
A. Depiction of one possible accuracy/RT curve. Percent correct responses increases monotonically as a function of RT and asymptotes at 90%. B. The total reward/RT curve for the accuracy/RT curve from panel A with the following additional assumptions: (i) observers complete as many trials as possible within a 30-minute window, (ii) completing a trial takes 1.5 seconds on top of the RT (due to stimulus presentation and between-trial breaks), and (iii) each correct answer results in 1 point, while incorrect answers result in 0 points. The optimal RT – the one which maximizes the total reward – is depicted with dashed lines.

#### 3.4.1 Trading off speed and accuracy

While observers are able to adjust their behavior to account for both accuracy and RT, they cannot do so optimally (Balcı et al. 2011; Bogacz et al. 2010; Simen et al. 2009; Starns and Ratcliff 2010, 2012; Tsetsos et al. 2015). In most cases observers take too long to decide, leading to slightly higher accuracy but substantially longer RTs than optimal (Bogacz et al. 2010; Simen et al. 2009; Starns and Ratcliff 2010, 2012). This effect occurs when observers have a fixed period of time to complete as many trials as possible (Bogacz et al. 2010; Simen et al. 2009; Starns and Ratcliff 2010, 2012), and in the more familiar design with a fixed number of trials per block (Starns and Ratcliff 2010, 2012). Further, observers take longer to decide for more difficult compared to easier conditions, even though optimizing the total reward demands that they always do the opposite (Oud et al. 2016; Starns and Ratcliff 2012). Older adults are even more suboptimal than college-age participants by this measure (Starns and Ratcliff 2010, 2012).

#### 3.4.2 Keeping a low error rate under implicit time pressure

Even though observers tend to overemphasize accuracy, they are also suboptimal in tasks that require an extreme emphasis on accuracy. This conclusion comes from a line of research on visual search in which observers are typically given unlimited amount of time to decide whether a target is present or not (Eckstein 2011). In certain situations, such as airport checkpoints or detecting tumors in mammograms, the goal is to keep a very low miss rate irrespective of RT, because misses can have dire consequences (Evans, Birdwell, and Wolfe 2013; Wolfe et al. 2013). The optimal RT can be derived from Figure 5A as the minimal RT that results in the desired accuracy rate. A series of studies by Wolfe and colleagues found that observers, even trained doctors and airport checkpoint screeners, are suboptimal in such tasks in that they allow overly high rates of misses (Evans et al. 2011, 2013; Wolfe et al. 2013; Wolfe, Horowitz, and Kenner 2005; Wolfe and Van Wert 2010). Further, this effect was robust and resistant to a variety of methods designed to help observers take longer in order to achieve higher accuracy (Wolfe et al. 2007) or correct motor errors (Van Wert, Horowitz, and Wolfe 2009). An explanation of this suboptimality based on capacity limits is rejected by two studies that found that observers can be induced to take longer time, and thus achieve higher accuracy, by first providing them with a block of high prevalence targets accompanied with feedback (Wolfe et al. 2007, 2013).

#### 3.4.3 Explaining suboptimality in the speed-accuracy tradeoff

Why do people appear to be unable to trade off speed and accuracy optimally? As above, it is possible to account for overly long RTs by postulating that, in addition to maximizing their total reward, observers place a premium on being accurate [***cost function***] (Balcı et al. 2011; Bogacz et al. 2010; Holmes and Cohen 2014). Another possibility is that observers’ judgments of elapsed time are noisy [***general***], and longer-than-optimal RTs lead to a higher reward rate than RTs that are shorter than optimal by the same amount (Simen et al. 2009; Zacksenhouse, Bogacz, and Holmes 2010). Finally, in some situations observers may also place a premium on speed [***cost function***], making it impossible to keep a very low error rate (Wolfe et al. 2013).

### 3.5 Confidence in one’s decision

The Bayesian approach prescribes how the posterior probability should be computed. While researchers typically examine the question of whether the stimulus with highest posterior probability is selected, it is also possible to examine whether observers can report the actual value of the posterior distribution or perform simple computations with it (Figure 6). In such cases, observers are asked to provide “metacognitive” confidence ratings about the accuracy of their decisions (Metcalfe and Shimamura 1994; Yeung and Summerfield 2012). Such studies rarely provide subjects with an explicit cost function (but see Kiani and Shadlen 2009; Rahnev et al. 2013), but in many cases reasonable assumptions can be made in order to derive optimal performance (see below).

**Figure 6.**
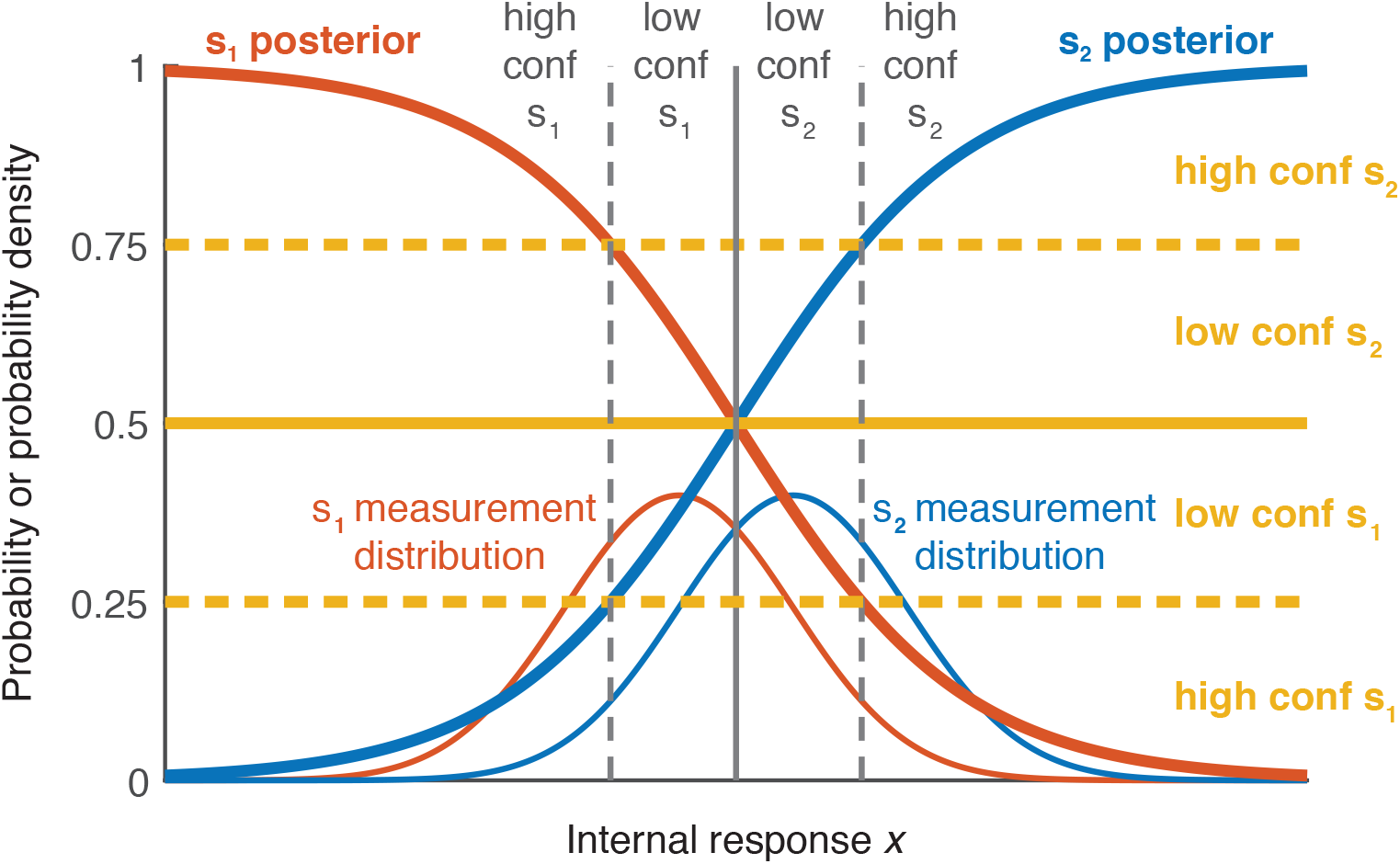
Depiction of how an observer should give confidence ratings. Similar to Figure 4, both the measurement distributions and posterior probabilities as a function of x assuming uniform prior are depicted. The confidence thresholds (depicted as yellow lines) correspond to criteria defined on x (depicted as thick gray lines). The horizontal thresholds and vertical criteria intersect on the posterior probability functions. The y-axis is probability density for the measurement distributions, and probability for the posterior probability functions (the y-axis ticks refer to the posterior probability).

#### 3.5.1 Over- and under-confidence (confidence calibration)

It is straightforward to construct a payoff structure for confidence ratings such that observers gain the most reward when their confidence reflects the posterior probability of being correct (e.g., Fleming et al. 2016; Massoni, Gajdos, and Vergnaud 2014). Most studies, however, do not provide observers with such a payoff structure, so assessing the optimality of the confidence ratings necessitates the further assumption that observers create a similar function internally. To test for optimality, we can then consider, for example, all trials in which an observer has 70% confidence of being correct, and test whether the average accuracy on those trials is indeed 70%. This type of relationship between confidence and accuracy is often referred to as confidence calibration (Baranski and Petrusic 1994). Studies of confidence have found that for certain tasks observers are overconfident (i.e., they overestimate their accuracy) (Adams 1957; Baranski and Petrusic 1994; Dawes 1980; Harvey 1997; Keren 1988; Koriat 2011) while for other tasks observers are underconfident (i.e., they underestimate their accuracy) (Baranski and Petrusic 1994; Björkman, Juslin, and Winman 1993; Dawes 1980; Harvey 1997; Winman and Juslin 1993). One pattern that emerges consistently is that overconfidence often occurs for difficult tasks while underconfidence appears in easy tasks (Baranski and Petrusic 1994, 1995, 1999), a phenomenon known as the *hard-easy effect* (Gigerenzer, Hoffrage, and Kleinbölting 1991). Similar results are seen for tasks outside of the perceptual domain such as answering general knowledge questions (Griffin and Tversky 1992). Over- and under-confidence are stable over different tasks (Ais et al. 2015; Song et al. 2011) and depend on non-perceptual factors such as one’s optimism bias (Ais et al. 2015).

#### 3.5.2 Dissociations of confidence and accuracy across different experimental conditions

While precise confidence calibration is computationally difficult, a weaker test of optimality examines whether experimental conditions that lead to the same performance are judged with the same level of confidence (even if this level is too high or too low). This test only requires that observers’ confidence ratings follow a consistent internal cost function across the two tasks. Many studies demonstrate dissociations between confidence and accuracy across tasks, thus showing that observers fail this weaker optimality test. For example, speeded responses can decrease accuracy but leave confidence unchanged (Baranski and Petrusic 1994; Vickers and Packer 1982), while slowed responses can lead to the same accuracy but lower confidence (Kiani, Corthell, and Shadlen 2014). Dissociations between confidence and accuracy have also been found in conditions that differ in attention (Rahnev, Bahdo, et al. 2012; Rahnev, Maniscalco, et al. 2011; Wilimzig et al. 2008), the variability of the perceptual signal (de Gardelle and Mamassian 2015; Koizumi, Maniscalco, and Lau 2015; Samaha et al. 2016; Song, Koizumi, and Lau 2015; Spence, Dux, and Arnold 2016; Zylberberg, Roelfsema, and Sigman 2014), the stimulus-onset asynchrony in metacontrast masking (Lau and Passingham 2006), the presence of unconscious information (Vlassova, Donkin, and Pearson 2014), and the relative timing of a concurrent saccade (Navajas, Sigman, and Kamienkowski 2014). Further, some of these biases seem to arise from individual differences that are stable across multiple sessions (de Gardelle and Mamassian 2015). Finally, dissociations between confidence and accuracy have been found in studies that applied transcranial magnetic stimulation (TMS) to the visual (Rahnev, Maniscalco, et al. 2012), premotor (Fleming et al. 2015), or frontal cortex (Chiang et al. 2014).

#### 3.5.3 Metacognitive sensitivity (confidence resolution)

The sections above were concerned with the average magnitude of confidence ratings over many trials. Another measure of interest is the degree of correspondence between confidence and accuracy on individual trials (Metcalfe and Shimamura 1994), called metacognitive sensitivity (Fleming and Lau 2014) or confidence resolution (Baranski and Petrusic 1994). Recently, Maniscalco & Lau (2012) developed a method to quantify how optimal an observer’s metacognitive sensitivity is. Their method computes *meta-d’*, a measure of how much information is available for metacognition, which can then be compared with the actual *d’* value. An optimal observer would have a *meta-d’*/*d’* ratio of 1. Maniscalco & Lau obtained a ratio of .77, suggesting a 23% loss of information for confidence judgments. Even though some studies that used the same measure but different perceptual paradigms found values close to 1 (Fleming et al. 2014), many others arrived at values substantially lower than 1 (Bang, Shekhar, and Rahnev 2017; Maniscalco and Lau 2015; Maniscalco, Peters, and Lau 2016; Massoni 2014; McCurdy et al. 2013; Schurger, Kim, and Cohen 2015; Sherman et al. 2015; Vlassova et al. 2014). Interestingly, at least one study has reported values significantly above 1, suggesting that in certain cases the metacognitive system has more information than was used for the primary decision (Charles et al. 2013), thus implying the presence of suboptimality in the perceptual decision.

#### 3.5.4 Confidence does not simply reflect the posterior probability of being correct

Another way of assessing the optimality of confidence ratings is to determine whether observers compute confidence in a manner consistent with the posterior probability of being correct. This is also a weaker condition than reporting the actual posterior probability of being correct, because it does not specify how observers should place decision boundaries between different confidence ratings, only that these boundaries should depend on the posterior probability of being correct. While one study found that confidence ratings are consistent with computations based on the posterior probability (Sanders, Hangya, and Kepecs 2016; but see Adler and Ma 2017b), others showed that either some (Aitchison et al. 2015; Navajas et al. 2017) or most (Adler and Ma 2017a; Denison et al. 2017) observers are described better by heuristic models in which confidence depends on uncertainty but not on the actual posterior probability of being correct.

Further, confidence judgments are influenced by a host of factors unrelated to the perceptual signal at hand and thus in violation of the principle that they should reflect the posterior probability of being correct. For example, emotional states, such as worry (Massoni 2014) and arousal (Allen et al. 2016), affect how sensory information relates to confidence ratings. Other factors, such as eye gaze stability (Schurger et al. 2015), working memory load (Maniscalco and Lau 2015) and age (Weil et al. 2013), affect the relationship between confidence and accuracy. Sequential effects have also been reported for confidence judgments such that a high confidence rating is more likely to follow a high, rather than low, confidence rating (Mueller and Weidemann 2008). Confidence dependencies exist even between different tasks, such as letter and color discrimination, that depend on different neural populations in the visual cortex (Rahnev et al. 2015). Inter-task confidence influences have been dubbed “confidence leak” and have been shown to be negatively correlated with observers’ metacognitive sensitivity (Rahnev et al. 2015).

Confidence has also been shown to exhibit a “positive evidence” bias (Maniscalco et al. 2016; Zylberberg, Barttfeld, and Sigman 2012). In 2-choice tasks, one can distinguish between sensory evidence in a trial that is congruent with the observer’s response on that trial (positive evidence) and sensory evidence that is incongruent with the response (negative evidence). Even though the perceptual decisions usually follow the optimal strategy of weighting equally both of these sources of evidence, confidence ratings are suboptimal in depending more heavily on the positive evidence (Koizumi et al. 2015; Maniscalco et al. 2016; Samaha et al. 2016; Song et al. 2015; Zylberberg et al. 2012).

#### 3.5.5 Explaining suboptimality in confidence ratings

Why do people appear to give inappropriate confidence ratings? Some components of over-and underconfidence can be explained by inappropriate transformation of internal evidence into probabilities [***general***] (Zhang and Maloney 2012), methodological considerations such as interleaving conditions with different difficulty levels, which can have inadvertent effects on the prior [***methodological***] (Drugowitsch, Moreno-Bote, and Pouget 2014), or even individual differences such as shyness about giving high confidence, which can be conceptualized as extra cost for high confidence responses [***cost function***]. Confidence-accuracy dissociations are often attributed to observers’ inability to maintain different criteria for different conditions, even if they are clearly distinguishable [***decision rule***] (Koizumi et al. 2015; Rahnev, Maniscalco, et al. 2011). The “positive evidence” bias [***decision rule***] introduced in the end of Section 3.5.4 can also account for many suboptimalities of confidence ratings.

More generally, it is possible that confidence ratings are not only based on the available perceptual evidence as assumed by most modeling approaches (Drugowitsch and Pouget 2012; Green and Swets 1966; Macmillan and Creelman 2005; Ratcliff and Starns 2009; Vickers 1979). Other theories postulate the existence of either different processing streams that contribute differentially to the perceptual decision and the subjective confidence judgment (Del Cul et al. 2009; Jolij and Lamme 2005; Weiskrantz 1996) or a second processing stage that determines the confidence judgment, and which builds upon the information in an earlier processing stage responsible for the perceptual decision (Bang et al. 2017; van den Berg, Yoo, and Ma 2017; Lau and Rosenthal 2011; Maniscalco and Lau 2010, 2016; Pleskac and Busemeyer 2010). Both types of models could be used to explain the various findings of suboptimal behavior and imply the existence of different measurement distributions for decision and confidence [***likelihood function***].

### 3.6 Comparing sensitivity in different tasks

The above sections discussed observers’ performance on a single task. Another way of examining optimality is to compare the performance on two related tasks. If the two tasks have a formal relationship, then an optimal observer’s sensitivity on the two tasks should follow that relationship.

#### 3.6.1 Comparing performance in 1-stimulus and 2-stimulus tasks

Visual sensitivity has traditionally been measured by employing either (1) a 1-stimulus (detection or discrimination) task in which a single stimulus from one of two stimulus classes is presented on each trial, or (2) a 2-stimulus task in which both stimulus classes are presented on each trial (see Section 1.3). Intuitively, 2-stimulus tasks are easier because the final decision is based on more perceptual information. Assuming independent processing of each stimulus, the relationship between the sensitivity on these two types of tasks can be mathematically defined: the sensitivity on the 2-stimulus task should be 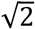 times higher than on the 1-stimulus task (Macmillan & Creelman, 2005; Figure 7). Nevertheless, empirical studies have often contradicted this predicted relationship: Many studies found sensitivity ratios smaller than 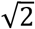 (Creelman and Macmillan 1979; Jesteadt 1974; Leshowitz 1969; Markowitz and Swets 1967; Pynn 1972; Schulman and Mitchell 1966; Swets and Green 1961; Viemeister 1970; Watson et al. 1973; Yeshurun, Carrasco, and Maloney 2008), though a few have found ratios larger than 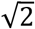 (Leshowitz 1969; Markowitz and Swets 1967; Swets and Green 1961).

**Figure 7.**
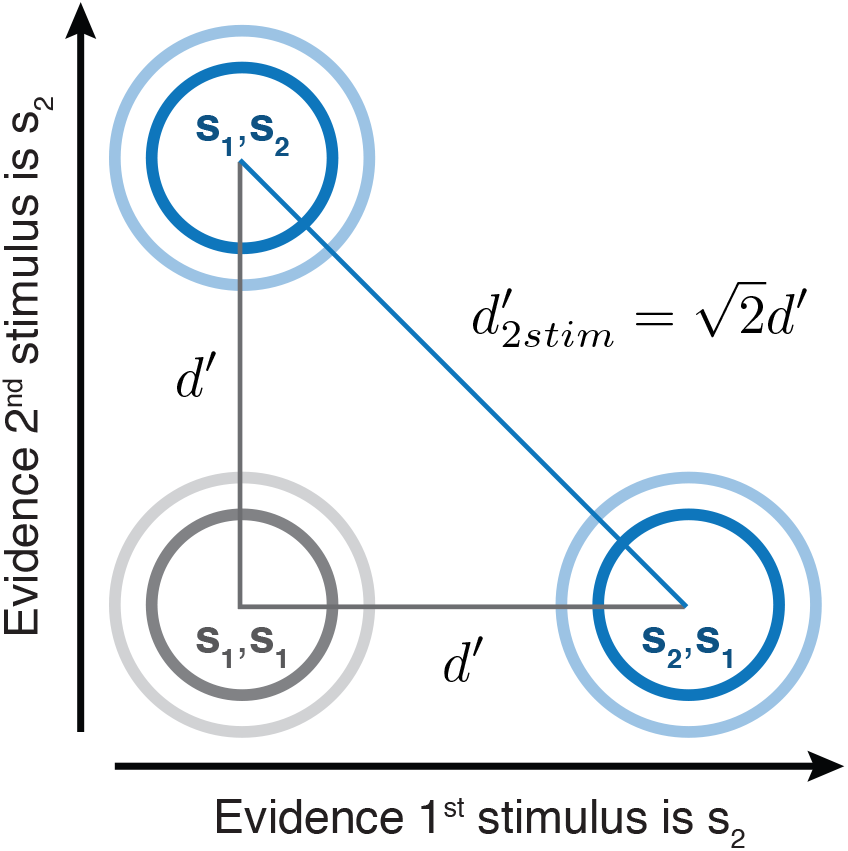
Depiction of the relationship between 1-stimulus and 2-stimulus tasks. Each axis corresponds to a 1-stimulus task (e.g., Figure 2). The three sets of concentric circles represent 2D circular Gaussian distributions corresponding to presenting two stimuli in a row (e.g., *s*_2_,*s*_1_ means that *s*_2_ was presented first and *s*_1_ was presented second). If the discriminability between *s*_1_ and *s*_2_ is d’ (1-stimulus task; gray lines in triangle), then the Pythagorean theorem gives us the expected discriminability between *s*_1_,*s*_2_ and *s*_2_,*s*_1_ (2-stimulus task; blue line in triangle).

#### 3.6.2 Comparing performance in other tasks

Many other comparisons between tasks have been performed. In temporal 2IFC tasks observers often have different sensitivity to the two stimulus intervals (García-Pérez and Alcalá-Quintana 2010, 2011; Yeshurun et al. 2008), suggesting an inability to distribute resources equally. Other studies find that longer inter-stimulus intervals in 2IFC tasks lead to decreases in sensitivity (Berliner and Durlach 1973; Kinchla and Smyzer 1967; Tanner 1961), presumably due to memory limitations. Further, choice variability on 3-choice tasks is greater than what would be predicted by a related 2-choice task (Drugowitsch et al. 2016). Creelman and Macmillan (1979) compared the sensitivity on 9 different psychophysical tasks and found a complex pattern of dependencies, many of which were at odds with optimal performance. Finally, Olzak (1985) demonstrated deviations from the expected relationship between detection and discrimination tasks.

An alternative approach to comparing an observer’s performance on different tasks is allowing observers to choose which tasks they prefer to complete and analyzing the optimality of these decisions. In particular, one can test for the presence of transitivity: if an observer prefers task A to task B and task B to task C, then the observer should choose task A to task C. Several studies suggest that human observers violate the transitivity principle both in choosing tasks (Zhang, Morvan, and Maloney 2010) and choosing stimuli (Tsetsos et al. 2016), though there is considerable controversy surrounding such findings (Davis-Stober et al. 2016; Kalenscher et al. 2010; Regenwetter et al. 2011, 2017; Regenwetter, Dana, and Davis-Stober 2010).

#### 3.6.3 Explaining suboptimality in between-task comparisons

Why does human performance on different tasks violate the expected relationship between these tasks? One possibility is that observers face certain capacity limits in one but not the other task that alter how the stimuli are encoded [***likelihood function***]. For example, compared to 1-stimulus tasks, the more complex 2-stimulus task requires the simultaneous processing of two stimuli. If limited resources hamper the processing of the second stimulus, then sensitivity in that task would fall short of what is predicted based on the 1-stimulus task.

In some experiments observers performed worse than expected on the 1-stimulus, rather than on the 2-stimulus task. A possible explanation of this effect is the presence of a larger “criterion jitter” in the 1-stimulus task, i.e., a larger variability in the decision criterion from trial to trial. Since 2-stimulus tasks involve the comparison of two stimuli on each trial, these tasks are less susceptible to criterion jitter. Such criterion variability, which could stem from sequential dependencies or even random criterion fluctuations (see Section 3.2), decreases the estimated stimulus sensitivity (Mueller and Weidemann 2008). The criterion jitter could also be due to computational imprecision [***general***] (Bays and Dowding 2017; Beck et al. 2012; Dayan 2014; Drugowitsch et al. 2016; Renart and Machens 2014; Whiteley and Sahani 2012; Wyart and Koechlin 2016). Such imprecision could arise from constraints at the neural level and may account for a large amount of choice suboptimality (Drugowitsch et al. 2016).

### 3.7 Cue combination

Studies of cue combination have been fundamental to the view that sensory perception is optimal (Trommershäuser, Körding, and Landy 2011). Cue combination (also called “cue integration”) is needed whenever different sensory features provide separate pieces of information about a single physical quantity. For example, auditory and visual signals can separately inform about the location of an object. Each cue provides imperfect information about the physical world, but different cues have different sources of variability. As a result, integrating the different cues can provide a more accurate and reliable estimate of the physical quantity of interest.

One can test for optimality in cue combination by comparing the perceptual estimate formed from two cues with the estimates formed from each cue individually. The optimal estimate is typically taken to be the one that maximizes precision (minimizes variability) across trials (Figure 8). When the variability for each cue is Gaussian and independent of the other cues, the maximum likelihood estimate (MLE) is a linear combination of the estimates from each cue, weighted by their individual reliabilities (Landy, Banks, and Knill 2011). Whether observers conform to this weighted sum formula can be readily tested psychophysically, and a large number of studies have done exactly this for different types of cues and tasks (see Ma, 2010 and Trommershäuser et al., 2011 for reviews).

**Figure 8.**
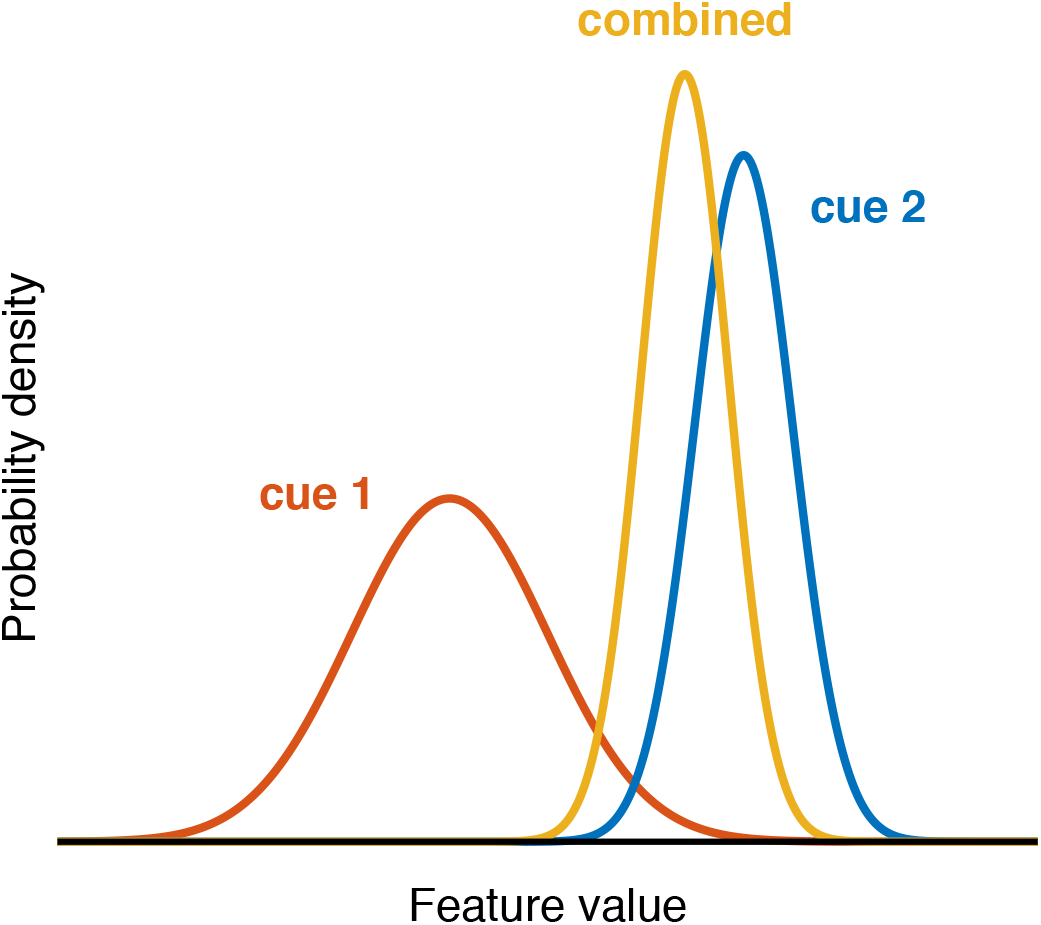
Optimal cue combination. Two cues that give independent information about the value of a sensory feature (red and blue curves) are combined to form a single estimate of the feature value (yellow curve). The combined cue distribution is narrower than both individual cue distributions, and its mean is closer to the mean of the distribution of the more informative cue.

In particular, the optimal mean perceptual estimate (*x*) after observing cue 1 (with feature estimate *x*_1_ and variance of 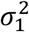) and cue 2 (with feature estimate *x*_2_ and variance of 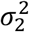) is:

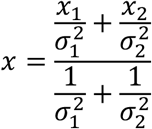

such that the feature estimate *x_i_* is weighted by its reliability 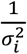 and the whole expression is normalized by the sum of the reliabilities. The optimal variance of the perceptual estimate (*σ*^2^) is:

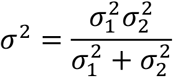

#### 3.7.1 Examples of optimality in cue combination

A classic example of cue combination is a study of visual-haptic cue combination by Ernst & Banks (2002). In this study, observers estimated the height of a rectangle using 1) only sight, 2) only touch, or 3) both sight and touch. Performance in the visual-haptic condition was well described by the MLE formula: the single cue measurements predicted both the reliability of the combined estimates and the weights given to each cue. Many studies have observed similar optimal cue combination behavior in a range of tasks estimating different physical quantities (Trommershäuser et al. 2011). These studies have investigated integration across two modalities (including vision, touch, audition, the vestibular sense, and proprioception; e.g., Alais & Burr, 2004; Ernst & Banks, 2002; Gu, Angelaki, & DeAngelis, 2008; van Beers, Sittig, & Denier van der Gon, 1996) and across two features in the same modality, such as various visual cues to depth (e.g., Jacobs 1999; Landy et al. 1995). Common among these experiments is that trained observers complete many trials of a psychophysical task, and the two cues provide similar estimates of the quantity of interest. Optimal cue combination has also been observed during sensory-motor integration (Maloney and Zhang 2010; Trommershäuser 2009; Wei and Körding 2011; Yeshurun et al. 2008).

#### 3.7.2 Examples of suboptimality in cue combination

Because optimality is often the hypothesized outcome in cue combination studies, findings of suboptimality may be underreported or underemphasized in the literature (Rosas and Wichmann 2011). Still, a number of studies have demonstrated suboptimal cue combination that violates some part of the MLE formula. These violations fall into two categories: 1) those in which the cues are integrated but are not weighted according to their independently-measured reliabilities, and 2) those in which estimates from two cues are no better than estimates from a single cue.

In the first category are findings from a wide range of combined modalities: visual-auditory (Battaglia, Jacobs, and Aslin 2003; Burr, Banks, and Morrone 2009; Maiworm and Röder 2011), visual-vestibular (Fetsch et al. 2012; Prsa, Gale, and Blanke 2012), visual-haptic (Battaglia, Kersten, and Schrater 2011; Rosas et al. 2005), and visual-visual (Knill and Saunders 2003; Rosas, Wichmann, and Wagemans 2007). For example, auditory and visual cues were not integrated according to the MLE rule in a localization task; instead, observers treated the visual cue as though it were more reliable than it really was (Battaglia et al. 2003). Similarly, visual and haptic texture cues were integrated according to their reliabilities, but observers underweighted the visual cue (Rosas et al. 2005). Suboptimal integration of visual and auditory cues was also found for patients with central vision loss, but not for patients with peripheral vision loss (Garcia et al. 2017).

In some of the above studies, cue misweighting was restricted to low-reliability cues: in a visual-vestibular heading task, observers overweighted vestibular cues when visual reliability was low (Fetsch et al. 2012), and in a visual-auditory temporal order judgment task, observers overweighted auditory cues when auditory reliability was low (Maiworm and Röder 2011). However, overweighting does not only occur within a limited range of reliabilities (e.g., Battaglia et al., 2003; Prsa et al., 2012).

Several studies have failed to find optimal cue combination in the temporal domain. In an audiovisual rate combination task, observers only partially integrated the auditory and visual cues, and they did not integrate them at all when the rates were very different (Roach, Heron, and McGraw 2006). Observers also overweight auditory cues in temporal order judgment tasks (Maiworm and Röder 2011) and temporal bisection tasks (Burr et al. 2009). It is well-established that when two cues give very different estimates, observers tend to discount one of them (Gepshtein et al. 2005; Jack and Thurlow 1973; Körding et al. 2007; Roach et al. 2006; Warren and Cleaves 1971), an effect which has been called “robust fusion” (Maloney and Landy 1989), which may arise from inferring that the two cues come from separate sources (Körding et al. 2007). However, in most of these studies, suboptimal cue combination was observed even when the cues gave similar estimates.

In the second category of suboptimal cue combination findings, two cues are no better than one (Chen and Tyler 2015; Drugowitsch, DeAngelis, et al. 2014; Landy and Kojima 2001; Oruç, Maloney, and Landy 2003; Rosas et al. 2005, 2007). (Note that some of these studies found a mix of optimal and suboptimal observers.) Picking the best cue is known as a “veto” type of cue combination (Bülthoff and Mallot 1988) and is considered a case of “strong fusion” (Clark and Yullie 1990; Landy et al. 1995). This is an even more serious violation of optimal cue combination, since it is as though no integration has taken place at all - the system either picks the best cue or in some cases does *worse* with two cues than with one.

Cues may also be mandatorily combined even when doing so is not suitable for the observer’s task. For example, texture and disparity information about slant is subsumed in a combined estimate, rendering the single cue estimates unrecoverable (Hillis et al. 2002). Interestingly, the single cue estimates are not lost in children, allowing them to outperform adults when the cues disagree (Nardini, Bedford, and Mareschal 2010). In a related finding, observers used multiple visual features to identify a letter even when the optimal strategy was to use only a single, relevant feature (Saarela and Landy 2015).

#### 3.7.3 Combining stimuli of the same type

So far, we have only considered cue combination studies in which the two cues come from different sensory modalities or dimensions. Suboptimal behavior has also been observed when combining cues from the same dimension. For example, Summerfield and colleagues showed that observers do not weight every sample stimulus equally in a decision (Summerfield and Tsetsos 2015). For simultaneous samples, observers underweight “outlier” stimuli lying far from the mean of the sample (de Gardelle and Summerfield 2011; Michael et al. 2015; Michael, de Gardelle, and Summerfield 2014; Vandormael et al. 2017). For sequential samples, observers overweight stimuli toward the end of the sequence (a recency effect) as well as stimuli that are similar to recently presented items (Bang and Rahnev 2017; Cheadle et al. 2014; Wyart, Myers, and Summerfield 2015). Observers also use only a subset of a sample of orientations to estimate its mean (Dakin 2001). More generally, accuracy on tasks with sequential samples is substantially lower than what would be predicted by sensory noise alone (Drugowitsch et al. 2016).

#### 3.7.4 Combining sensory and motor cues

Suboptimal cue integration has also been found in sensory-motor tasks. For example, when integrating the path of a pointing movement with online visual feedback, observers underestimate the uncertainty indicated by the feedback (Körding and Wolpert 2004). In a pointing task in which observers are rewarded for physically touching the correct visual target, observers underweight the difficulty of the motor task by aiming for a small target even though the perceptual information indicating the correct target was also uncertain (Fleming, Maloney, and Daw 2013). Similar biases were reported in related task (Landy et al. 2007). Within the action domain (and so beyond our focus on perception), Maloney and Zhang (2010) have reviewed studies showing both optimal and suboptimal behavior.

#### 3.7.5 Cue combination in children

Optimal cue integration takes time to develop. Children are suboptimal until around 10 years of age when combining multisensory (Gori et al. 2008; Nardini et al. 2008; Petrini et al. 2014) or visual (Dekker et al. 2015; Nardini et al. 2010) cues.

#### 3.7.6 Explaining suboptimal cue combination

Why do people sometimes appear to combine cues suboptimally? One possible explanation is that observers do not have accurate representations of the reliability of the cues (Knill and Saunders 2003; Rosas et al. 2005) because learning the reliability is difficult [***methodological***]. This methodological issue is particularly acute when the cues are new to the observer. For example, in one task for which cue combination was suboptimal, observers haptically explored a surface with a single finger to estimate its slant. However, observers may have little experience with single-finger slant estimation, since multiple fingers or the whole hand might ordinarily be used for such a task (Rosas et al. 2005). Alternatively, cue combination may be suboptimal when one cue provides all information in parallel but the other cue provides information serially (Plaisier et al. 2014). Reliability estimation might also be difficult when the reliability is very low. This possibility may apply to studies in which observers were optimal within a range of sensory reliability, but not outside it (Fetsch et al. 2012; Maiworm and Röder 2011).

Some authors suggest that another reason for over- or under-weighting a certain cue could be prior knowledge about how cues ought to be combined [***prior***]. This could include a prior assumption about how likely a cue is to be related to the desired physical property (Battaglia et al. 2011; Ganmor, Landy, and Simoncelli 2015), how likely two cue types are to correspond to one another (and thus be beneficial to integrate) (Roach et al. 2006), or a general preference to rely on a particular modality, such as audition in a timing task (Maiworm and Röder 2011).

For certain tasks, some researchers question the assumptions of the MLE model, such as Gaussian noise [***likelihood function***] (Burr et al. 2009) or the independence of the neural representations of the two cues [***likelihood function***] (Rosas et al. 2007). In other cases, it appears that observers use alternative cost functions by, for example, taking RT into account [***cost function***] (Drugowitsch, DeAngelis, et al. 2014).

“Robust averaging,” or down-weighting of outliers, has been observed when observers must combine multiple pieces of information that give very different perceptual estimates. Such down-weighting can stem from adaptive gain changes [***likelihood function***] that result in highest sensitivity to stimuli close to the mean of the sample (or in the sequential case, the subset of the sample that has been presented so far; Summerfield & Tsetsos, 2015). This adaptive gain mechanism is similar to models of sensory adaptation (Barlow 1990; Carandini and Heeger 2012; Wark, Lundstrom, and Fairhall 2007). By following principles of efficient coding that place the largest dynamic range at the center of the sample (Barlow 1961; Brenner, Bialek, and de Ruyter van Steveninck 2000; Wainwright 1999), different stimuli receive unequal weightings. Note that psychophysical studies in which stimulus variability is low would not be expected to show this kind of suboptimality (Cheadle et al. 2014).

It is debated whether suboptimal cue combination in children reflects a switching strategy (Adams 2016), immature neural mechanisms for integrating cues, or whether the developing brain is optimized for a different task, such as multisensory calibration or conflict detection (Gori et al. 2008; Nardini et al. 2010).

### 3.8 Other examples of suboptimality

Thus far we have specifically focused on tasks where the optimal behavior can be specified mathematically in a relatively uncontroversial manner (though see Section 4.2 below). However, the issue of optimality has been discussed in a variety of other contexts.

#### 3.8.1 Perceptual biases, illusions, and improbabilities

A number of basic visual biases have been documented. Some examples include repulsion of orientation or motion direction estimates away from cardinal directions (Figure 9A; Jastrow, 1892; Rauber & Treue, 1998), a bias to perceive speeds as slower than they are when stimuli are low contrast (Stone & Thompson, 1992; Thompson, 1982; but see Thompson, Brooks, & Hammett, 2006), a bias to perceive surfaces as convex (Langer and Bülthoff 2001; Sun and Perona 1997), and a bias to perceive visual stimuli closer to fixation than they are (whereas the opposite is true for auditory stimuli; Odegaard, Wozny, & Shams, 2015).

**Figure 9.**
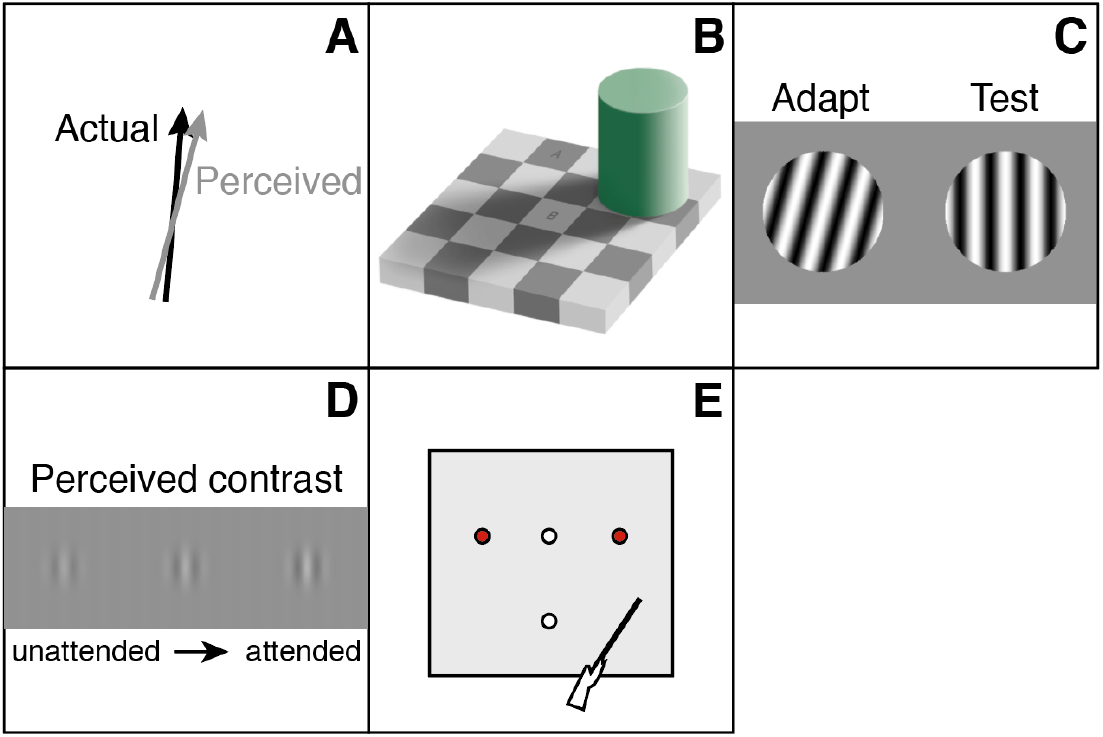
Examples of illusions and biases. A) Cardinal repulsion. A nearly vertical (or horizontal) line looks more tilted away from the cardinal axis than it is. B) Adelson’s checkerboard brightness illusion. Square B appears to be brighter than square A, even though the two squares have the same luminance. Figure courtesy of Michael Bach (http://www.michaelbach.de/ot/lum-adelsonCheckShadow/index.html). C) Tilt aftereffect. After viewing a tilted adapting grating (left), observers perceive a vertical test grating (right) to be tilted away from the adaptor. D) Effects of spatial attention on contrast appearance (Carrasco, Ling, and Read 2004). An attended grating appears to have higher contrast than the same grating when it is unattended. E) Effects of action affordances on perceptual judgments (Witt 2011). Observers judge an object to be closer (far white circle compared to near white circle) relative to the distance between two landmark objects (red circles) when they are holding a tool that allows them to reach that object than when they have no tool.

When biases, context, or other factors lead to something looking dramatically different from its physical reality, we might call it a visual illusion. A classic example is the brightness illusion (Figure 9B) in which two squares on a checkerboard appear to be different shades of gray even though they actually have the same luminance (Adelson 1993). Perceptual illusions persist even when the observer knows about the illusion and even after thousands of trials of exposure (Gold et al. 2000).

Some illusions are difficult to reconcile with existing theories of optimal perception. Anderson, O’Vari, & Barth (2011), for example, reported strong percepts of illusory surfaces that were improbable according to optimal frameworks for contour synthesis. In the size-weight illusion, smaller objects are perceived as heavier than larger objects of the same weight, even though the prior expectation is that smaller objects are lighter (Brayanov and Smith 2010; Peters, Ma, and Shams 2016).

#### 3.8.2 Adaptation

Adaptation is a widespread phenomenon in sensory systems in which responsiveness to prolonged or repeated stimuli is reduced (Webster 2015). As some researchers have discussed (Wei and Stocker 2015), adaptation could be seen as suboptimal from a Bayesian perspective because subsequent perceptual estimates tend to diverge from rather than conform to the prior stimulus. For example, after prolonged viewing of a line tilted slightly away from vertical, a vertical line looks tilted in the opposite direction (the “tilt aftereffect”, Figure 9C; Gibson & Radner, 1937). Or, after viewing motion in a certain direction, a stationary stimulus appears to drift in the opposite direction (Wohlgemuth 1911). After adapting to a certain color, perception is biased toward the complementary color (Sabra 1989; Turnbull 1961), and after adapting to a specific face, another face appears more different from that face than it would have otherwise (Webster et al. 2004; Webster and MacLeod 2011). In all of these examples, perception is repelled away from the prior stimulus, which, at least on the surface, appears suboptimal (but see Section 3.8.5).

#### 3.8.3 Appearance changes due to visual attention

The same physical stimulus can also be perceived in different ways depending on the state of visual attention. Directing spatial attention to a stimulus can make it appear larger (Anton-Erxleben, Henrich, and Treue 2007), faster (Anton-Erxleben, Herrmann, and Carrasco 2013; Fuller, Park, and Carrasco 2009; Turatto, Vescovi, and Valsecchi 2007), and brighter (Tse 2005), and to have higher spatial frequency (Abrams, Barbot, and Carrasco 2010; Gobell and Carrasco 2005) and higher contrast (Figure 9D; Carrasco et al. 2004; Liu, Abrams, and Carrasco 2009; Störmer, Mcdonald, and Hillyard 2009) than it would otherwise. Often attentional effects improve performance on a visual task, but sometimes they make performance worse (Ling and Carrasco 2006; Yeshurun and Carrasco 1998), demonstrating inflexibility in the system.

#### 3.8.4 Cognition-based biases

Other studies have documented visual biases associated with more cognitive factors, including action affordances (Witt 2011), motivation (Balcetis 2015), and language (Lupyan 2012). For example, when people will be reaching for an object with a tool that allows them to reach further, they report the object as looking closer than when they will be reaching without the tool (Figure 9E; Witt, Proffitt, & Epstein, 2005). In the linguistic domain, calling an object a “triangle” leads observers to report the object as having more equal sides than when the object is called “three-sided” (Lupyan 2016). How much these more cognitive factors affect perception *per se*, as opposed to post-perceptual judgments, and to what extent the observed visual biases are mediated by attention, remain controversial questions (Firestone and Scholl 2016).

#### 3.8.5 Explaining these other examples of apparent suboptimality

Why are people prone to certain biases and illusions? Some biases and illusions have been explained as arising from priors in the visual system [***prior***]. Misperceptions of motion direction (Weiss et al. 2002) and biases in reporting the speed of low contrast stimuli (Stocker and Simoncelli 2006a; Thompson 1982; Vintch and Gardner 2014) have been explained as optimal percepts for a visual system with a prior for slow motion (Stocker and Simoncelli 2006a; Weiss et al. 2002). Such a prior is motivated by the fact that natural objects tend to be still or move slowly but has been empirically challenged by subsequent research (Hammett et al. 2007; Hassan and Hammett 2015; Thompson et al. 2006; Vaziri-Pashkam and Cavanagh 2008). Priors have been be invoked to explain many other biases and illusions (Brainard et al. 2006; Girshick, Landy, and Simoncelli 2011; Glennerster et al. 2006; Raviv, Ahissar, and Loewenstein 2012). The suggestion is that these priors have been made stable over a lifetime and influence perception even when they do not apply (i.e., in a laboratory task).

Optimal decoding of sensory representations in one task can be accompanied by suboptimal biases in another task using the same stimuli. For example, in a fine motion discrimination task, observers seem to overweight the neurons tuned away from the discrimination boundary, because these neurons distinguish best between the two possible stimuli. This overweighting could explain why motion direction judgments in an interleaved estimation task are biased away from the boundary (Jazayeri and Movshon 2007). Another interpretation of these results is in terms of an improper decision rule (Zamboni et al. 2016). Specifically, observers may discard sensory information related to the rejected decision outcome [***decision rule***] (Bronfman et al. 2015; Fleming et al. 2013; Luu and Stocker 2016), an effect known as *self-consistency bias* (Stocker and Simoncelli 2008).

Various efforts have been made to explain adaptation in the framework of Bayesian optimality (Grzywacz and Balboa 2002; Hohwy, Roepstorff, and Friston 2008; Schwiedrzik et al. 2014; Snyder et al. 2015). One of the most well-developed lines of work explains the repulsive effects of adaptation as a consequence of efficient coding [***likelihood function***] (Stocker and Simoncelli 2006b). In this framework, a sensory system adapts to maximize its dynamic range around the value of previous input. This change in coding does not affect the prior (as might be expected in a Bayesian treatment of adaptation) but rather affects the likelihood function. Specifically, it skews new observations away from the adapted stimulus, giving rise to repulsive aftereffects. A similar principle has been suggested to explain why perceptual estimates are repelled from long-term priors, such as those determined by the statistics of natural images (Wei and Stocker 2012, 2015).

## 4. ASSESSING OPTIMALITY: NOT A USEFUL GOAL IN ITSELF

The extensive review in the previous section demonstrates that general claims about the optimality of human perceptual decision making are empirically false. But there are also theoretical reasons to turn away from assessing optimality as a primary research goal.

### 4.1 Challenges in defining optimality

Section 2 introduced a formal definition of optimality based on Bayesian Decision Theory. However, the question of what phenomena should be considered optimal vs. suboptimal quickly becomes complicated in many actual applications. There are at least two issues that are not straightforward to address.

The first issue concerns the exact form of the cost function. Bayesian Decision Theory postulates that observers should minimize the expected loss. However, observers may reasonably prefer to minimize the maximum loss, minimize the variability of the losses, or optimize some other quantity. Therefore, behavior that is suboptimal according to standard Bayesian Decision Theory may be optimal according to other definitions. A related, and deeper, problem is that some observers may also try to minimize other quantities such as time spent, level of boredom, or metabolic energy expended (Carrasco 2011; Lennie 2003). What appears to be a suboptimal decision on a specific task may be optimal when all these other variables are taken into account (Beck et al. 2012; Bowers and Davis 2012a). Even the clearest cases of suboptimal decision rules (e.g., the self-consistency bias) could be construed as part of a broader optimality (e.g., being self-consistent may be important for other goals). In a Bayesian framework, taking into account extra variables requires that each of the LPCD components is defined over all these variables. If one pursues this logic, it leads to a cost function that operates over our entire evolutionary history. We do not think efforts to understand such cost functions should be abandoned, but specifying them quantitatively is impossible given our current knowledge.

The second issue concerns whether optimality should depend on the likelihood, prior, and cost function adopted by the observer. In order to be able to review a large literature using consistent assumptions, we defined a set Standard Assumptions and labeled any deviation from these assumptions as suboptimal. This approach is by no means uncontroversial. For example, priors based on a lifetime of experience may be inflexible, so one could consider the Standard Assumption about following the experimenter-defined prior overly restrictive. An alternative view could be that suboptimal behavior concerns only deviations from the experimenter-defined quantities that are under observers’ control (Tenenbaum and Griffiths 2006; Yu and Cohen 2009). The problem with this definition is that it introduces a new variable to consider – what exactly is truly under observers’ control – which is often hard to determine. A third approach is to define optimality exclusively in terms of the decision rule regardless of what likelihood, prior, and cost function the observer adopts. In this view, observers are under no obligation to follow the experimenter’s instructions (e.g., they are free to bring in their own priors and cost function). The problem with this approach is that failing to adopt the proper prior or cost function can result in just as much missed objective reward as adopting an improper decision rule. Similar problems apply to “improper” likelihood functions: as an extreme example, a strategy in which the observer closes her eyes (resulting in a non-informative likelihood function) and chooses actions randomly has to be labeled “optimal” because the decision rule is optimal. The ambiguity regarding the role of the likelihood, prior, or cost function points to the difficulties in constructing a general-purpose definition of optimality.

In short, optimality is impossible to define in the abstract. It is only well-defined in the context of a set of specific assumptions, rendering general statements about the optimality (or suboptimality) of human perceptual decisions meaningless.

### 4.2 Optimality claims in and of themselves have limited value

The current emphasis on optimality is fueled by the belief that demonstrating optimality in perception provides us with important insight. On the contrary, simply stating that observers are optimal is of limited value for two main reasons.

First, it is unclear when a general statement about the optimality of perceptual decisions is supposed to apply. While most experimental work focuses on very simple tasks, it is widely recognized that the computational complexity of many real-world tasks makes optimality unachievable by the brain (Bossaerts and Murawski 2017; Cooper 1990; Gershman et al. 2015; van Rooij 2008; Tsotsos 1993). Further, in many situations, the brain cannot be expected to have complete knowledge of the likelihood function, which all but guarantees that the decision rule will be suboptimal (Beck et al. 2012). (It should be noted that attempting to incorporate observers’ computational capacities or knowledge brings back the problems related to how one defines optimality discussed in Section 4.1.) Therefore, general statements about optimality must be intended only for the simplest cases of perceptual decisions (although, as Section 3 demonstrated, even for these cases, suboptimality is ubiquitous).

Second, even for a specific task, statements about optimality alone are insufficient to predict behavior. Instead, to predict future perceptual decisions, one needs to specify each part of the process underlying the decision. Within the Bayesian framework, for example, one needs to specify each LPCD component, which goes well beyond a statement that “observers are optimal.”

Is it useless to compare human performance to optimal performance? Absolutely not. *Within the context of a specific model*, demonstrating optimal or suboptimal performance is immensely helpful (Goodman et al. 2015; Tauber et al. 2017). Such demonstrations can support or challenge components of the model and suggest ways to alter the model to accommodate actual behavior. But the critical part here is the model, not the optimality.

## 5. TOWARD A STANDARD OBSERVER MODEL

If there are so many empirical examples of suboptimality (Section 3) and optimality can be challenging even to define (Section 4), then what is the way forward?

### 5.1 Creating and testing observer models

Psychophysics has a long history of creating ideal observer models (Geisler 2011; Green and Swets 1966; Ulehla 1966). These models specify a set of assumptions about how sensory information is represented internally and add an optimal decision rule in order to generate predictions about behavior. The motivation behind these models has been to test the collective set of assumptions incorporated into the model. However, over time, the “ideal” part of the ideal observer models has become dominant, culminating in the current outsized emphasis on demonstrating the optimality of the decision rule – what we call the *optimality approach*. Even frameworks such as “bounded rationality” (Gigerenzer and Selten 2002; Simon 1957) or “computational rationality” (Gershman et al. 2015), which explicitly concern themselves with the limitations of the decision-making process, still place the greatest emphasis on the optimality of the decision rule.

The emphasis on the decision rule in the optimality approach has led to an overly flexible treatment of the other LPCD components (Bowers and Davis 2012a). This issue is especially problematic due to the inherent degeneracy of Bayesian Decision Theory (Acerbi 2014): different combinations of the likelihood, prior, cost, and decision rule can lead to the same expected loss. Further, for any likelihood, cost function, and decision rule, a prior can be found for which that decision rule is optimal (“complete class theorem”) (Berger 1985; Jaynes 2003).

To eliminate the flexibility of the optimality approach, the field should return to the original intention of building ideal observer models, namely to test the collective set of assumptions incorporated into such models. To this end, we propose that researchers drop the “ideal” and shift emphasis to building, simply, “observer models.” Creating observer models should differ from the current optimality approach in two critical ways. First, whether or not the decision rule is optimal should be considered irrelevant. Second, the nature of the decision rule should not be considered more important than the nature of the other components.

These two simple changes address the pitfalls of the optimality approach. Within the optimality approach, a new finding is often modeled using flexibly-chosen LPCD components (Bowers and Davis 2012a). Then, depending on the inferred decision rule, a conclusion is reached that observers are optimal (or suboptimal). At this point, the project is considered complete and a general claim is made about optimality (or suboptimality). As others have pointed out, this approach has led to many “just-so stories” (Bowers and Davis 2012a), since the assumptions of the model are not rigorously tested. On the contrary, when building observer models (e.g., in the Bayesian framework), a new finding is used to generate *hypotheses* about a particular LPCD component (Maloney and Mamassian 2009). Hypotheses about the likelihood, prior, or cost function are considered as important as hypotheses about the decision rule. Critically, unlike in the optimality approach, this step is considered just the beginning of the process! The hypotheses are then examined in detail while evidence is gathered for or against them. Researchers can formulate alternative hypotheses to explain a given dataset and evaluate them using model comparison techniques. In addition, researchers can conduct follow-up experiments in which they test their hypotheses using different tasks, stimuli, and observers. There are researchers who already follow this approach, and we believe the field would benefit from adopting it as the standard practice. In Box 1, we list specific steps for implementing observer models within a Bayesian framework (the steps will be similar regardless of the framework).

#### Box 1. Implementing observer models within a Bayesian framework

1. Describe the complete generative model including assumptions about what information the observer is using to perform the task (e.g., stimulus properties, training, experimenter’s instructions, feedback, explicit vs. implicit rewards, response time pressure, etc.).
2. Specify the assumed likelihood function, prior, and cost function. If multiple options are plausible, test them in different models.
3. Derive both the optimal decision rule, as well as plausible alternative decision rules. Compare their ability to fit the data.
4. Interpret the results with respect to what has been learned about each LPCD component, not optimality per se. Specify how the conclusions depend on the assumptions about the other LPCD components.
5. Most importantly, follow up on any new hypotheses about LPCD components with additional studies in order to avoid “just-so stories.”
6. New hypotheses that prove to be general eventually become part of the Standard Observer Model (see Section 5.2).

Two examples demonstrate the process of implementing observer models. A classic example concerns the existence of Gaussian variability in the measurement distribution. This assumption has been extensively tested for decades (Green and Swets 1966; Macmillan and Creelman 2005), thus eventually earning its place among the Standard Assumptions in the field. A second example comes from the literature on speed perception. A classic finding is that reducing the contrast of a slow-moving stimulus reduces its apparent speed (Stone and Thompson 1992; Thompson 1982). A popular Bayesian explanation for this effect is that most objects in natural environments are stationary, so the visual system has a prior for slow speeds. Consequently, when sensory information is uncertain, as occurs at low contrasts, slow-biased speed perception could be considered “optimal” (Weiss et al. 2002). Importantly, rather than stopping at this claim, researchers have investigated the hypothetical slow motion prior in follow-up studies. One study quantitatively inferred observers’ prior speed distributions under the assumption of a Bayesian decision rule (Stocker and Simoncelli 2006a). Other researchers tested the slow motion prior and found that, contrary to its predictions, high speed motion at low contrast can appear to move *faster* than its physical speed (Hammett et al. 2007; Hassan and Hammett 2015; Thompson et al. 2006). These latter studies challenged the generality of the slow motion prior hypothesis.

### 5.2 Creating a Standard Observer Model

We believe that an overarching goal of the practice of creating and testing observer models is the development of a Standard Observer Model that predicts observers’ behavior on a wide variety of perceptual tasks. Such a model would be a significant achievement for the science of perceptual decision making. It is difficult – perhaps impossible – to anticipate what form the Standard Observer Model will take. It may be a Bayesian model (Maloney and Mamassian 2009), a “bag of tricks” (Ramachandran 1990), a neural network (Yamins et al. 2014), etc. However, regardless of the framework in which they were originally formulated, hypotheses with overwhelming empirical support will become part of the Standard Observer Model. In this context, perhaps the most damaging aspect of the current outsized emphasis on optimality is that while it has generated many hypotheses, few of them have received sufficient subsequent attention to justify inclusion in (or exclusion from) the eventual Standard Observer Model.

We suggest that immediate progress can be made by a concerted effort to test the hypotheses that have already been proposed to explain suboptimal decisions. To facilitate this effort, here we compile the hypotheses generated in the course of explaining the findings from Section 3.

Within a Bayesian framework, these hypotheses relate to the likelihood function, prior, cost function, or decision rule (the LPCD components). Further, a few of them are general and apply to several LPCD components, and a few are methodological considerations. In some cases, essentially the same hypothesis was offered in the context of several different empirical effects. We summarize these hypotheses in Table 1. Note that the table by no means exhaustively covers all existing hypotheses that deserve to be thoroughly tested.

**Table 1.**
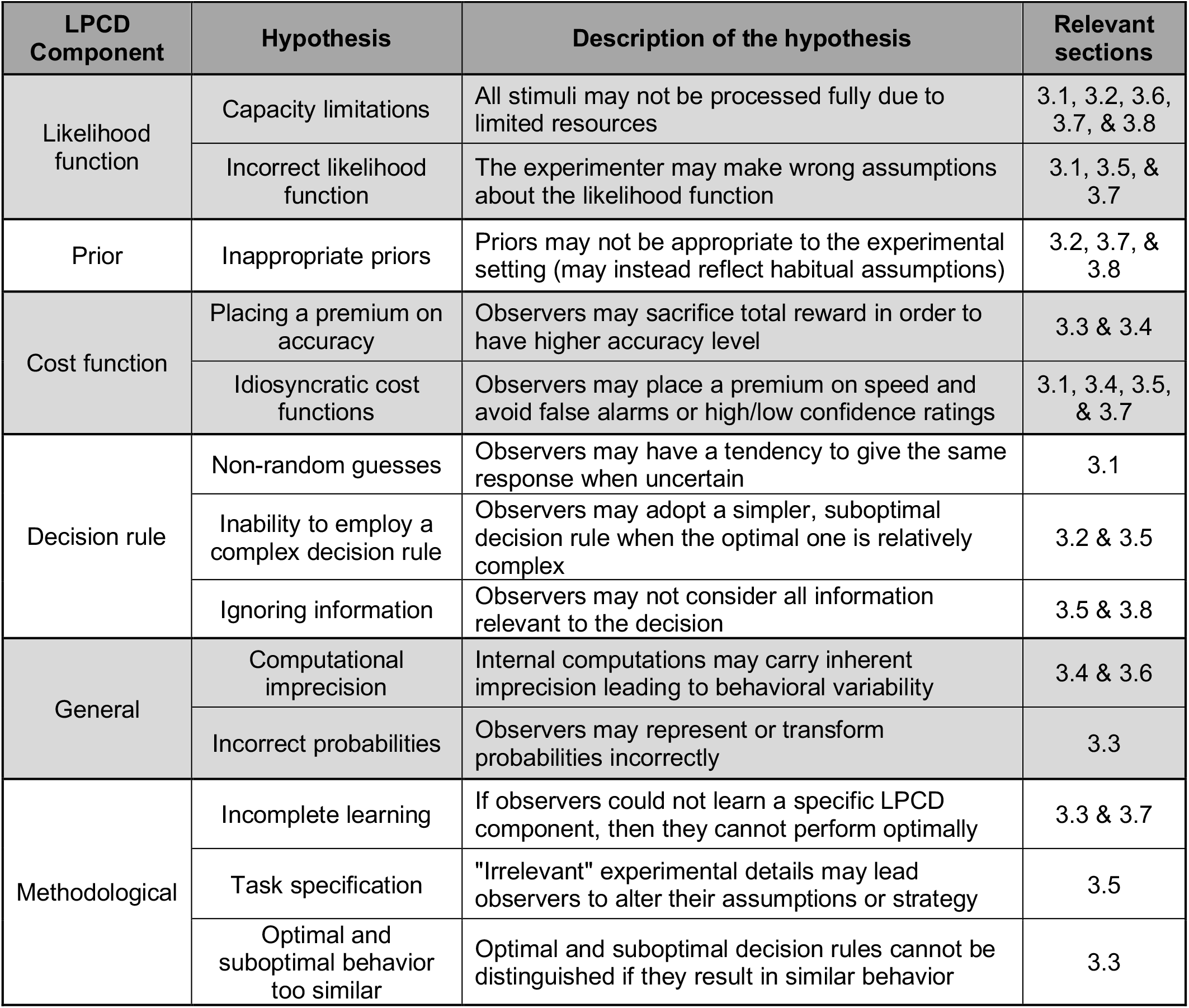
Summary of hypotheses proposed to account for suboptimal decisions.

Table 1 classifies instances of deficient learning as methodological issues. This choice is not to downplay the problem of learning. Questions of how observers acquire their priors and cost functions are of utmost importance, and meaningful progress has already been made on this front (Acerbi et al. 2014; Acerbi, Wolpert, and Vijayakumar 2012; Beck et al. 2012; Geisler and Najemnik 2013; Gekas et al. 2013; Seriès and Seitz 2013). Here we categorize deficient learning as a methodological issue when, due to the experimental setup, an observer cannot acquire the relevant knowledge even though they have the capacity to do so.

Future research should avoid the methodological issues from Table 1. In particular, great care must be taken to ensure that observers’ assumptions in performing a task match exactly the assumptions implicit in the analysis.

We have stated the hypotheses in Table 1 at a fairly high level to succinctly capture the broad categories from our review. Much of the work ahead will be to break each high-level hypothesis down into multiple, specific hypotheses and incorporate these hypotheses into observer models. For example, statements about “inappropriate priors” or “capacity limitations” prompt more finegrained hypotheses about specific priors or limitations whose ability to predict behavior can be tested. Some hypotheses, like capacity limitations, have already been investigated extensively – for example in studies of attention and working memory (e.g., Carrasco 2011; Cowan 2005). Turning our existing knowledge of these phenomena into concrete observer models that predict perceptual decisions is an exciting direction for the field. Other hypotheses, like placing a premium on accuracy, have not been tested extensively and therefore should still be considered “just-so stories” (Bowers and Davis 2012a). Thus, the real work ahead lies in verifying, rejecting, and expanding the hypotheses generated from findings of suboptimal perceptual decisions.

### 5.3 Implications of abandoning the optimality approach

Abandoning the optimality approach has at least two immediate implications for research practices.

First, researchers should stop focusing on optimality. What should be advertised in the title and abstract of a paper is not the optimality but what is learned about the components of the perceptual process. One of the central questions in perceptual decision making is how best to characterize the sources that corrupt decisions (Beck et al. 2012; Drugowitsch et al. 2016; Hanks and Summerfield 2017; Wyart and Koechlin 2016). By shifting attention away from optimality, the effort to build complete observer models sharpens the focus on this question.

Second, new model development should not unduly emphasize optimal models. According to some Bayesian theorists, models that assume optimal behavior are intrinsically preferable to models that do not. This preference stems from the argument that because people can approximate optimal behavior on some tasks, they must possess the machinery for fully optimal decisions (Drugowitsch & Pouget, 2012). Many models have been judged positively for supporting optimal decision rules: probabilistic population codes for allowing optimal cue combination (Ma et al. 2006), neural sampling models for allowing marginalization (which is needed in many optimal decision rules) (Fiser et al. 2010), and drift diffusion models for allowing optimal integration of information across time (Bogacz 2007). The large body of findings of suboptimality reviewed here, however, should make this reasoning suspect: if the brain is built to make optimal decisions, then why does it produce so many suboptimal ones? It is also important to remember that close-to-optimal behavior can also be produced by suboptimal decision rules (Bowers and Davis 2012a; Maloney and Mamassian 2009; Shen and Ma 2016). Influential theories postulate that evolutionary pressures produced heuristic but useful, rather than normative, behavior (Gigerenzer and Brighton 2009; Juslin et al. 2009; Simon 1956). Thus, models should be judged solely based on their ability to describe actual behavior and not on their ability to support optimal decision rules.

## 6. CONCLUSION

Are perceptual decisions optimal? A substantial body of research appears to answer this question in the affirmative. Here we showed instead that every category of perceptual tasks that lends itself to optimality analysis features numerous findings of suboptimality. Perceptual decisions cannot therefore be claimed to be optimal in general. In addition, independent of the empirical case against optimality, we questioned whether a focus on optimality per se can lead to any real progress. Instead, we advocated for a return to building complete observer models with an equal focus on all model components. Researchers should aim for their models to capture all the systematic weirdness of human behavior rather than preserve an aesthetic ideal. To facilitate this effort, we compiled the hypotheses generated in the effort to explain the findings of suboptimality reviewed here. The real work ahead lies in testing these hypotheses, with the ultimate goal of developing a Standard Observer Model of perceptual decision making.

## Acknowledgements

We thank Luigi Acerbi, William Adler, Stephanie Badde, Michael Landy, Wei Ji Ma, Larry Maloney, Brian Maniscalco, Jonathan Winawer, and five reviewers for many helpful comments and discussions. D.R. was supported by a startup grant from Georgia Institute of Technology. R.N.D. was supported by National Institutes of Health National Eye Institute F32 EY025533 and T32 EY007136 to New York University.

## References

Abrahamyan, Arman, Laura Luz Silva, Steven C. Dakin, Matteo Carandini, and Justin L. Gardner. 2016. “Adaptable History Biases in Human Perceptual Decisions.” Proceedings of the National Academy of Sciences 113(25):E3548–57. Retrieved August 2, 2016 (http://www.pnas.org/lookup/doi/10.1073/pnas.1518786113).

Abrams, Jared, Antoine Barbot, and Marisa Carrasco. 2010. “Voluntary Attention Increases Perceived Spatial Frequency.” Attention, Perception, & Psychophysics 72(6):1510–21. Retrieved (http://eutils.ncbi.nlm.nih.gov/entrez/eutils/elink.fcgi?dbfrom=pubmed&id=20675797&retmode=ref&cmd=prlinks).

Acerbi Luigi. 2014. “Complex Internal Representations in Sensorimotor Decsision Making: A Bayesian Investigation.” University of Edinburgh. Retrieved (https://www.era.lib.ed.ac.uk/bitstream/handle/1842/16233/Acerbi2015.pdf?sequence=1&isAllowed=y).

Acerbi, Luigi, Sethu Vijayakumar, and Daniel M. Wolpert. 2014. “On the Origins of Suboptimality in Human Probabilistic Inference.” edited by J. Beck. PLoS computational biology 10(6):e1003661. Retrieved July 18, 2014 (http://dx.plos.org/10.1371/journal.pcbi.1003661).

Acerbi, Luigi, Daniel M. Wolpert, and Sethu Vijayakumar. 2012. “Internal Representations of Temporal Statistics and Feedback Calibrate Motor-Sensory Interval Timing.” PLoS computational biology 8(11):e1002771. Retrieved September 28, 2015 (http://journals.plos.org/ploscompbiol/article?id=10.1371/journal.pcbi.1002771).

Ackermann John F. and Michael S. Landy. 2015. “Suboptimal Decision Criteria Are Predicted by Subjectively Weighted Probabilities and Rewards.” Attention, perception & psychophysics 77(2):638–58. Retrieved December 29, 2014 (http://www.ncbi.nlm.nih.gov/pubmed/25366822).

Adams J. K. 1957. “A Confidence Scale Defined in Terms of Expected Percentages.” The American journal of psychology 70(3):432–36.

Adams Wendy J. 2016. “The Development of Audio-Visual Integration for Temporal Judgements” edited by U. R. Beierholm. PLOS Computational Biology 12(4):e1004865. Retrieved January 18, 2018 (http://dx.plos.org/10.1371/journal.pcbi.1004865).

Adelson E. H. 1993. “Perceptual Organization and the Judgment of Brightness.” Science 262(5142):2042–44.

Adler William T. and Wei Ji Ma. 2017a. “Human Confidence Reports Account for Sensory Uncertainty but in a Non-Bayesian Way.” bioRxiv 93203. Retrieved January 18, 2018 (https://www.biorxiv.org/content/early/2017/05/18/093203).

Adler William T. and Wei Ji Ma. 2017b. “Limitations of Proposed Signatures of Bayesian Confidence.” bioRxiv 218222. Retrieved January 18, 2018 (https://www.biorxiv.org/content/early/2017/11/16/218222).

Ais, Joaquín, Ariel Zylberberg, Pablo Barttfeld, and Mariano Sigman. 2015. “Individual Consistency in the Accuracy and Distribution of Confidence Judgments.” Cognition 146:377–86. Retrieved November 4, 2015 (http://www.ncbi.nlm.nih.gov/pubmed/26513356).

Aitchison, Laurence, Dan Bang, Bahador Bahrami, and Peter E. Latham. 2015. “Doubly Bayesian Analysis of Confidence in Perceptual Decision-Making.” PLoS computational biology 11(10):e1004519.

Alais, David and David Burr. 2004. “The Ventriloquist Effect Results from near-Optimal Bimodal Integration.” Current biology 14(3):257–62. Retrieved June 20, 2016 (http://www.ncbi.nlm.nih.gov/pubmed/14761661).

Allen, Micah et al. 2016. “Unexpected Arousal Modulates the Influence of Sensory Noise on Confidence.” eLife 5:e18103. Retrieved (http://elifesciences.org/lookup/doi/10.7554/eLife.18103).

Anderson Barton L., Judit O’Vari, and Hilary Barth. 2011. “Non-Bayesian Contour Synthesis.” Current Biology 21(6):492–96. Retrieved (http://linkinghub.elsevier.com/retrieve/pii/S0960982211001746).

Anton-Erxleben Katharina, Christian Henrich, and Stefan Treue. 2007. “Attention Changes Perceived Size of Moving Visual Patterns.” Journal of Vision 7(11):5.1-9. Retrieved (http://eutils.ncbi.nlm.nih.gov/entrez/eutils/elink.fcgi?dbfrom=pubmed&id=17997660&retmode=ref&cmd=prlinks).

Anton-Erxleben Katharina, Katrin Herrmann, and Marisa Carrasco. 2013. “Independent Effects of Adaptation and Attention on Perceived Speed.” Psychological science 24(2):150–59. Retrieved (http://eutils.ncbi.nlm.nih.gov/entrez/eutils/elink.fcgi?dbfrom=pubmed&id=23241456&retmode=ref&cmd=prlinks).

Balcetis E. 2015. “Approach and Avoidance as Organizing Structures for Motivated Distance Perception.” Emotion Review. Retrieved (http://emr.sagepub.com/content/early/2015/05/29/1754073915586225.abstract).

Balcı Fuat et al. 2011. “Acquisition of Decision Making Criteria: Reward Rate Ultimately Beats Accuracy.” Attention, perception & psychophysics 73(2):640–57. Retrieved September 11, 2015 (http://www.pubmedcentral.nih.gov/articlerender.fcgi?artid=3383845&tool=pmcentrez&rendertype=abstract).

Bang, Ji Won and Dobromir Rahnev. 2017. “Stimulus Expectation Alters Decision Criterion but Not Sensory Signal in Perceptual Decision Making.” Scientific Reports 7:17072. Retrieved (http://www.nature.com/articles/s41598-017-16885-2).

Bang Ji Won, Medha Shekhar, and Dobromir Rahnev. 2017. “Sensory Noise Increases Metacognitive Efficiency.” bioRxiv 189399. Retrieved January 29, 2018 (https://www.biorxiv.org/content/early/2017/09/15/189399).

Baranski J. V and W. M. Petrusic. 1994. “The Calibration and Resolution of Confidence in Perceptual Judgments.” Perception & psychophysics 55(4):412–28. Retrieved April 15, 2014 (http://www.ncbi.nlm.nih.gov/pubmed/8036121).

Baranski J. V and W. M. Petrusic. 1995. “On the Calibration of Knowledge and Perception.” Canadian journal of experimental psychology 49(3):397–407. Retrieved April 15, 2014 (http://www.ncbi.nlm.nih.gov/pubmed/9183984).

Baranski J. V. and W. M. Petrusic. 1999. “Realism of Confidence in Sensory Discrimination.” Perception & psychophysics 61(7):1369–83. Retrieved April 15, 2014 (http://www.ncbi.nlm.nih.gov/pubmed/10572465).

Barlow H. B. 1961. “Possible Principles Underlying the Transformation of Sensory Messages.” in Sensory Communication, edited by W. A. Rosenblith. MIT Press.

Barlow H. B. 1990. “A Theory about the Functional Role and Synaptic Mechanism of Visual after-Effects.” Vision: Coding and efficiency. Retrieved (http://books.google.com/books?hl=en&lr=&id=xGJ_DxN3eygC&oi=fnd&pg=PA363&dq=a+theory+about+the+functional+role+and+synaptic+mechanism+of+visual+after+effects&ots=VsSUzK0vpB&sig=lZX28LU68XpGk9T8zoLwY8WOJBs).

Battaglia Peter W., Robert A. Jacobs, and Richard N. Aslin. 2003. “Bayesian Integration of Visual and Auditory Signals for Spatial Localization.” Journal of the Optical Society of America A, Optics and image science 20(7):1391–97.

Battaglia Peter W., Daniel Kersten, and Paul R. Schrater. 2011. “How Haptic Size Sensations Improve Distance Perception.” PLoS computational biology 7(6):e1002080. Retrieved (http://eutils.ncbi.nlm.nih.gov/entrez/eutils/elink.fcgi?dbfrom=pubmed&id=21738457&retmode=ref&cmd=prlinks).

Bays Paul M. and Ben A. Dowding. 2017. “Fidelity of the Representation of Value in Decision-Making” edited by J. O’Reilly. PLOS Computational Biology 13(3):e1005405. Retrieved March 14, 2017 (http://www.ncbi.nlm.nih.gov/pubmed/28248958).

Beck Jeffrey M., Wei Ji Ma, Xaq Pitkow, Peter E. Latham, and Alexandre Pouget. 2012. “Not Noisy, Just Wrong: The Role of Suboptimal Inference in Behavioral Variability.” Neuron 74(1):30–39. Retrieved February 28, 2013 (http://www.ncbi.nlm.nih.gov/pubmed/22500627).

van Beers R. J., A. C. Sittig, and J. J. Denier van der Gon. 1996. “How Humans Combine Simultaneous Proprioceptive and Visual Position Information.” Experimental brain research 111(2):253–61.

van den Berg, Ronald, Aspen H. Yoo, and Wei Ji Ma. 2017. “Fechner’s Law in Metacognition: A Quantitative Model of Visual Working Memory Confidence.” Psychological Review 124(2):197–214.

Berger James O. 1985. Statistical Decision Theory and Bayesian Analysis. New York: Springer.

Berliner J. E. and N. I. Durlach. 1973. “Intensity Perception. IV. Resolution in Roving-Level Discrimination.” The Journal of the Acoustical Society of America 53(5):1270–87. Retrieved September 30, 2015 (http://www.ncbi.nlm.nih.gov/pubmed/4712555).

Björkman M., P. Juslin, and A. Winman. 1993. “Realism of Confidence in Sensory Discrimination: The Underconfidence Phenomenon.” Perception & psychophysics 54(1):75–81. Retrieved March 24, 2016 (http://www.ncbi.nlm.nih.gov/pubmed/8351190).

Bogacz Rafal. 2007. “Optimal Decision-Making Theories: Linking Neurobiology with Behaviour.” Trends in cognitive sciences 11(3):118–25. Retrieved September 18, 2014 (http://www.sciencedirect.com/science/article/pii/S1364661307000290).

Bogacz, Rafal, Eric Brown, Jeff Moehlis, Philip Holmes, and Jonathan D. Cohen. 2006. “The Physics of Optimal Decision Making: A Formal Analysis of Models of Performance in Two-Alternative Forced-Choice Tasks.” Psychological Review 113(4):700–765. Retrieved (http://doi.apa.org/getdoi.cfm?doi=10.1037/0033-295X.113.4.700).

Bogacz, Rafal, Peter T. Hu, Philip J. Holmes, and Jonathan D. Cohen. 2010. “Do Humans Produce the Speed-Accuracy Trade-off That Maximizes Reward Rate?” Quarterly journal of experimental psychology (2006) 63(5):863–91. Retrieved September 5, 2015 (http://www.pubmedcentral.nih.gov/articlerender.fcgi?artid=2908414&tool=pmcentrez&rendertype=abstract).

Bohil Corey J. and W. Todd Maddox. 2001. “Category Discriminability, Base-Rate, and Payoff Effects in Perceptual Categorization.” Perception & psychophysics 63(2):361–76.

Bohil Corey J. and W. Todd Maddox. 2003a. “A Test of the Optimal Classifier’s Independence Assumption in Perceptual Categorization.” Perception & psychophysics 65(3):478–93.

Bohil Corey J. and W. Todd Maddox. 2003b. “On the Generality of Optimal versus Objective Classifier Feedback Effects on Decision Criterion Learning in Perceptual Categorization.” Memory & cognition 31(2):181–98.

Bossaerts, Peter and Carsten Murawski. 2017. “Computational Complexity and Human Decision-Making.” Trends in Cognitive Sciences 21(12):917–29. Retrieved (http://dx.doi.org/10.1016/j.tics.2017.09.005).

Bowers Jeffrey S. and Colin J. Davis. 2012a. “Bayesian Just-so Stories in Psychology and Neuroscience.” Psychological bulletin 138(3):389–414. Retrieved September 5, 2015 (http://eutils.ncbi.nlm.nih.gov/entrez/eutils/elink.fcgi?dbfrom=pubmed&id=22545686&retmode=ref&cmd=prlinks).

Bowers Jeffrey S. and Colin J. Davis. 2012b. “Is That What Bayesians Believe? Reply to Griffiths, Chater, Norris, and Pouget (2012).” Psychological Bulletin 138(3):423–26. Retrieved (http://doi.apa.org/getdoi.cfm?doi=10.1037/a0027750).

Brainard David H. et al. 2006. “Bayesian Model of Human Color Constancy.” Journal of Vision 6(11):1267–81.

Brayanov J. B. and M. A. Smith. 2010. “Bayesian and ‘Anti-Bayesian’ Biases in Sensory Integration for Action and Perception in the Size-Weight Illusion.” Journal of Neurophysiology 103(3):1518–31. Retrieved (http://jn.physiology.org/cgi/doi/10.1152/jn.00814.2009).

Brenner N., W. Bialek, and R. de Ruyter van Steveninck. 2000. “Adaptive Rescaling Maximizes Information Transmission.” Neuron 26(3):695–702. Retrieved (http://eutils.ncbi.nlm.nih.gov/entrez/eutils/elink.fcgi?dbfrom=pubmed&id=10896164&retmode=ref&cmd=prlinks).

Bronfman Zohar Z. et al. 2015. “Decisions Reduce Sensitivity to Subsequent Information.” Proceedings of the Royal Society B 282(1810):20150228. Retrieved March 9, 2017 (http://www.ncbi.nlm.nih.gov/pubmed/26108628).

Brooke J. B. and A. W. MacRae. 1977. “Error Patterns in the Judgment and Production of Numerical Proportions.” Perception & Psychophysics 21(4):336–40. Retrieved October 2, 2015 (http://www.springerlink.com/index/10.3758/BF03199483).

Bülthoff H. H. and H. A. Mallo. 1988. “Integration of Depth Modules: Stereo and Shading.” Journal of the Optical Society of America A, Optics and image science 5(10):1749–58. Retrieved (http://eutils.ncbi.nlm.nih.gov/entrez/eutils/elink.fcgi?dbfrom=pubmed&id=3204438&retmode=ref&cmd=prlinks).

Burr, David, Martin S. Banks, and Maria Concetta Morrone. 2009. “Auditory Dominance over Vision in the Perception of Interval Duration.” Experimental Brain Research 198(1):49–57. Retrieved (http://eutils.ncbi.nlm.nih.gov/entrez/eutils/elink.fcgi?dbfrom=pubmed&id=19597804&retmode=ref&cmd=prlinks).

Busemeyer Jerome R. and In Jae Myung. 1992. “An Adaptive Approach to Human Decision Making: Learning Theory, Decision Theory, and Human Performance.” Journal of Experimental Psychology: General 121(2):177–94. Retrieved December 30, 2014 (http://psycnet.apa.org/journals/xge/121/2/177.html).

Busse L. et al. 2011. “The Detection of Visual Contrast in the Behaving Mouse.” Journal of Neuroscience 31(31):11351–61. Retrieved August 25, 2016 (http://www.jneurosci.org/cgi/doi/10.1523/JNEUROSCI.6689-10.2011).

Carandini, Matteo and David J. Heeger. 2012. “Normalization as a Canonical Neural Computation.” Nature reviews. Neuroscience 13(1):51–62. Retrieved February 28, 2013 (http://www.pubmedcentral.nih.gov/articlerender.fcgi?artid=3273486&tool=pmcentrez&rendertype=abstract).

Carrasco Marisa. 2011. “Visual Attention: The Past 25 Years.” Vision research 51(13):1484–1525. Retrieved September 17, 2013 (http://www.pubmedcentral.nih.gov/articlerender.fcgi?artid=3390154&tool=pmcentrez&rendertype=abstract).

Carrasco, Marisa, Sam Ling, and Sarah Read. 2004. “Attention Alters Appearance.” Nature neuroscience 7(3):308–13. Retrieved February 28, 2013 (http://www.ncbi.nlm.nih.gov/pubmed/14966522).

Charles, Lucie, Filip Van Opstal, Sébastien Marti, and Stanislas Dehaene. 2013. “Distinct Brain Mechanisms for Conscious versus Subliminal Error Detection.” NeuroImage 73:80–94. Retrieved September 1, 2015 (http://www.ncbi.nlm.nih.gov/pubmed/23380166).

Cheadle, Sam et al. 2014. “Adaptive Gain Control during Human Perceptual Choice.” Neuron 81(6):1429–41. Retrieved March 24, 2014 (http://www.cell.com/article/S0896627314000518/fulltext).

Chen, Chien-Chung and Christopher William Tyler. 2015. “Shading Beats Binocular Disparity in Depth from Luminance Gradients: Evidence against a Maximum Likelihood Principle for Cue Combination.” PloS one 10(8):e0132658. Retrieved March 24, 2016 (http://journals.plos.org/plosone/article?id=10.1371/journal.pone.0132658).

Chiang, Tzu-Ching, Ru-Band Lu, Shulan Hsieh, Yun-Hsuan Chang, and Yen-Kuang Yang. 2014. “Stimulation in the Dorsolateral Prefrontal Cortex Changes Subjective Evaluation of Percepts.” edited by M. Gray. PloS one 9(9):e106943. Retrieved September 29, 2014 (http://dx.plos.org/10.1371/journal.pone.0106943).

Clark J. J. and Alan L. Yullie. 1990. Data Fusion for Sensory Information Processing. Boston: Kluwer Academic Publishers.

Cooper Gregory F. 1990. “The Computational Complexity of Probabilistic Inference Using Bayesian Belief Networks.” Artificial Intelligence 42(2–3):393–405.

Cowan Nelson. 2005. Working Memory Capacity. New York: Psychology Press.

Creelman, C. D. and Neil A. Macmillan. 1979. “Auditory Phase and Frequency Discrimination: A Comparison of Nine Procedures.” Journal of experimental psychology. Human perception and performance 5(1):146–56. Retrieved September 30, 2015 (http://www.ncbi.nlm.nih.gov/pubmed/528924).

Del Cul, A, S. Dehaene, P. Reyes, E. Bravo, and A. Slachevsky. 2009. “Causal Role of Prefrontal Cortex in the Threshold for Access to Consciousness.” Brain 132(Pt 9):2531–40. Retrieved March 7, 2013 (http://www.ncbi.nlm.nih.gov/pubmed/19433438).

Dakin S. C. 2001. “Information Limit on the Spatial Integration of Local Orientation Signals.” Journal of the Optical Society of America A, Optics and image science 18(5):1016–26. Retrieved (http://eutils.ncbi.nlm.nih.gov/entrez/eutils/elink.fcgi?dbfrom=pubmed&id=11336204&retmode=ref&cmd=prlinks).

Davis-Stober Clintin P., Sanghyuk Park, Nicholas Brown, and Michel Regenwetter. 2016. “Reported Violations of Rationality May Be Aggregation Artifacts.” Proceedings of the National Academy of Sciences of the United States of America 113(33):E4761-3. Retrieved March 9, 2017 (http://www.ncbi.nlm.nih.gov/pubmed/27462103).

Dawes R. M. 1980. “Confidence in Intellectual vs. Confidence in Perceptual Judgments.” Pp. 327–45 in Similarity and choice: Papers in honor of Clyde Coombs, edited by E. D. Lantermann and H. Fege. Bern Han Huber.

Dayan, Peter. 2014. “Rationalizable Irrationalities of Choice.” Topics in cognitive science 6(2):204–28. Retrieved November 13, 2014 (http://doi.wiley.com/10.1111/tops.12082).

Dekker Tessa M. et al. 2015. “Late Development of Cue Integration Is Linked to Sensory Fusion in Cortex.” Current biology 25(21):2856–61. Retrieved (http://eutils.ncbi.nlm.nih.gov/entrez/eutils/elink.fcgi?dbfrom=pubmed&id=26480841&retmode=ref&cmd=prlinks).

Denison Rachel N., William T. Adler, Marisa Carrasco, and Wei Ji Ma. 2017. “Humans Flexibly Incorporate Attention-Dependent Uncertainty into Perceptual Decisions and Confidence.” bioRxiv 175075. Retrieved January 19, 2018 (https://www.biorxiv.org/content/early/2017/08/10/175075).

Drugowitsch, Jan, Gregory C. DeAngelis, Dora E. Angelaki, and Alexandre Pouget. 2015. “Tuning the Speed-Accuracy Trade-off to Maximize Reward Rate in Multisensory Decision-Making.” eLife 4:e06678. Retrieved March 14, 2017 (http://www.ncbi.nlm.nih.gov/pubmed/26090907).

Drugowitsch, Jan, Gregory C. DeAngelis, Eliana M. Klier, Dora E. Angelaki, and Alexandre Pouget. 2014. “Optimal Multisensory Decision-Making in a Reaction-Time Task.” eLife 3:e03005. Retrieved April 24, 2015 (http://elifesciences.org/content/early/2014/06/14/eLife.03005.abstract).

Drugowitsch, Jan, Rubén Moreno-Bote, Anne K. Churchland, Michael N. Shadlen, and Alexandre Pouget. 2012. “The Cost of Accumulating Evidence in Perceptual Decision Making.” The Journal of neuroscience 32(11):3612–28. Retrieved February 28, 2013 (http://www.pubmedcentral.nih.gov/articlerender.fcgi?artid=3329788&tool=pmcentrez&rendertype=abstract).

Drugowitsch, Jan, Rubén Moreno-Bote, and Alexandre Pouget. 2014. “Relation between Belief and Performance in Perceptual Decision Making.” edited by S. Ben Hamed. PloS one 9(5):e96511. Retrieved July 12, 2014 (http://dx.plos.org/10.1371/journal.pone.0096511).

Drugowitsch, Jan and Alexandre Pouget. 2012. “Probabilistic vs. Non-Probabilistic Approaches to the Neurobiology of Perceptual Decision-Making.” Current opinion in neurobiology 22(6):963–69. Retrieved September 19, 2014 (http://www.pubmedcentral.nih.gov/articlerender.fcgi?artid=3513621&tool=pmcentrez&rendertype=abstract).

Drugowitsch, Jan, Valentin Wyart, Anne-Dominique Devauchelle, and Etienne Koechlin. 2016. “Computational Precision of Mental Inference as Critical Source of Human Choice Suboptimality.” Neuron 92(6):1398–1411. Retrieved (http://dx.doi.org/10.1016/j.neuron.2016.11.005).

Eberhardt, Frederick and David Danks. 2011. “Confirmation in the Cognitive Sciences: The Problematic Case of Bayesian Models.” Minds and Machines 21(3):389–410. Retrieved (http://link.springer.com/10.1007/s11023-011-9241-3).

Eckstein M. P. 2011. “Visual Search: A Retrospective.” Journal of Vision 11(5):14–14. Retrieved December 29, 2014 (http://www.journalofvision.org/content/11/5/14.abstract).

Ernst Marc O. and Martin S. Banks. 2002. “Humans Integrate Visual and Haptic Information in a Statistically Optimal Fashion.” Nature 415(6870):429–33. Retrieved November 7, 2014 (http://dx.doi.org/10.1038/415429a).

Evans Karla K., Robyn L. Birdwell, and Jeremy M. Wolfe. 2013. “If You Don’t Find It Often, You Often Don’t Find It: Why Some Cancers Are Missed in Breast Cancer Screening.” PloS one 8(5):e64366. Retrieved December 28, 2014 (http://www.pubmedcentral.nih.gov/articlerender.fcgi?artid=3667799&tool=pmcentrez&rendertype=abstract).

Evans Karla K., Rosemary H. Tambouret, Andrew Evered, David C. Wilbur, and Jeremy M. Wolfe. 2011. “Prevalence of Abnormalities Influences Cytologists’ Error Rates in Screening for Cervical Cancer.” Archives of pathology & laboratory medicine 135(12):1557–60. Retrieved December 29, 2014 (http://www.pubmedcentral.nih.gov/articlerender.fcgi?artid=3966132&tool=pmcentrez&rendertype=abstract).

Fechner G. T. 1860. Elemente Der Psychophysik. Leipzig: Breitkopf und Härtel.

Feng, Samuel, Philip Holmes, Alan Rorie, and William T. Newsome. 2009. “Can Monkeys Choose Optimally When Faced with Noisy Stimuli and Unequal Rewards?” PLoS computational biology 5(2):e1000284. Retrieved December 2, 2014 (http://www.pubmedcentral.nih.gov/articlerender.fcgi?artid=2631644&tool=pmcentrez&rendertype=abstract).

Fetsch Christopher R., Alexandre Pouget, Gregory C. Deangelis, and Dora E. Angelaki. 2012. “Neural Correlates of Reliability-Based Cue Weighting during Multisensory Integration.” Nature Neuroscience 15(1):146–54. Retrieved (http://eutils.ncbi.nlm.nih.gov/entrez/eutils/elink.fcgi?dbfrom=pubmed&id=22101645&retmode=ref&cmd=prlinks).

Firestone, Chaz and Brian J. Scholl. 2016. “Cognition Does Not Affect Perception: Evaluating the Evidence for ‘Top-Down’ Effects.” Behavioral and brain sciences 39:e229. Retrieved July 22, 2015 (http://www.ncbi.nlm.nih.gov/pubmed/26189677).

Fischer, Jason and David Whitney. 2014. “Serial Dependence in Visual Perception.” Nature Neuroscience 17(5):738–43. Retrieved April 1, 2014 (http://dx.doi.org/10.1038/nn.3689).

Fiser, József, Pietro Berkes, Gergo Orbán, and Máté Lengyel. 2010. “Statistically Optimal Perception and Learning: From Behavior to Neural Representations.” Trends in cognitive sciences 14(3):119–30. Retrieved July 16, 2014 (http://www.pubmedcentral.nih.gov/articlerender.fcgi?artid=2939867&tool=pmcentrez&rendertype=abstract).

Fitts Paul M. 1966. “Cognitive Aspects of Information Processing: III. Set for Speed versus Accuracy.” Journal of Experimental Psychology 71(6):849–57.

Fleming Stephen M. et al. 2015. “Action-Specific Disruption of Perceptual Confidence.” Psychological science 26(1):89–98. Retrieved November 27, 2014 (http://www.ncbi.nlm.nih.gov/pubmed/25425059).

Fleming Stephen M. and Hakwan Lau. 2014. “How to Measure Metacognition.” Frontiers in Human Neuroscience 8. Retrieved July 16, 2014 (http://journal.frontiersin.org/Journal/10.3389/fnhum.2014.00443/abstract).

Fleming Stephen M., Laurence T. Maloney, and Nathaniel D. Daw. 2013. “The Irrationality of Categorical Perception.” Journal of Neuroscience 33(49):19060–70. Retrieved December 4, 2013 (http://eutils.ncbi.nlm.nih.gov/entrez/eutils/elink.fcgi?dbfrom=pubmed&id=24305804&retmode=ref&cmd=prlinks).

Fleming Stephen M., Sébastien Massoni, Thibault Gajdos, and Jean-Christophe Vergnaud. 2016. “Metacognition about the Past and Future: Quantifying Common and Distinct Influences on Prospective and Retrospective Judgments of Self-Performance.” Neuroscience of Consciousness 2016(1):niw018. Retrieved January 18, 2018 (https://academic.oup.com/nc/article-lookup/doi/10.1093/nc/niw018).

Fleming Stephen M., J. Ryu, J. G. Golfinos, and K. E. Blackmon. 2014. “Domain-Specific Impairment in Metacognitive Accuracy Following Anterior Prefrontal Lesions.” Brain. Retrieved August 7, 2014 (http://brain.oxfordjournals.org/content/early/2014/08/06/brain.awu221.long).

Forstmann Birte U., Roger Ratcliff, and Eric-Jan Wagenmakers. 2016. “Sequential Sampling Models in Cognitive Neuroscience: Advantages, Applications, and Extensions.” Annual review of psychology 67:641–66. Retrieved (http://www.annualreviews.org/eprint/2stAyEdsCkSk9MpsHMDV/full/10.1146/annurev-psych-122414-033645).

Fritsche, Matthias, Pim Mostert, and Floris P. de Lange. 2017. “Opposite Effects of Recent History on Perception and Decision.” Current Biology. Retrieved (http://dx.doi.org/10.1016/j.cub.2017.01.006).

Fründ, Ingo, Felix A. Wichmann, and Jakob H. Macke. 2014. “Quantifying the Effect of Intertrial Dependence on Perceptual Decisions.” Journal of vision 14(7). Retrieved July 11, 2014 (http://www.ncbi.nlm.nih.gov/pubmed/24944238).

Fuller, Stuart, Yunsoo Park, and Marisa Carrasco. 2009. “Cue Contrast Modulates the Effects of Exogenous Attention on Appearance.” Vision Research 49(14):1825–37. Retrieved (http://eutils.ncbi.nlm.nih.gov/entrez/eutils/elink.fcgi?dbfrom=pubmed&id=19393260&retmode=ref&cmd=prlinks).

Ganmor, Elad, Michael S. Landy, and Eero P. Simoncelli. 2015. “Near-Optimal Integration of Orientation Information across Saccades.” Journal of vision 15(16):8. Retrieved January 18, 2018 (http://jov.arvojournals.org/article.aspx?doi=10.1167/15.16.8).

García-Pérez Miguel A. and Rocío Alcalá-Quintana. 2010. “The Difference Model with Guessing Explains Interval Bias in Two-Alternative Forced-Choice Detection Procedures.” Journal of Sensory Studies 25(6):876–98. Retrieved (http://doi.wiley.com/10.1111/j.1745-459X.2010.00310.x).

García-Pérez Miguel A. and Rocío Alcalá-Quintana. 2011. “Interval Bias in 2AFC Detection Tasks: Sorting out the Artifacts.” Attention, Perception, & Psychophysics 73(7):2332–52. Retrieved August 25, 2016 (http://www.springerlink.com/index/10.3758/s13414-011-0167-x).

Garcia Sara E. et al. 2017. “Multisensory Cue Combination After Sensory Loss: Audio-Visual Localization in Patients With Progressive Retinal Disease.” Journal of Experimental Psychology: Human Perception and Performance. Retrieved February 17, 2017 (http://doi.apa.org/getdoi.cfm?doi=10.1037/xhp0000344).

de Gardelle, Vincent and Pascal Mamassian. 2015. “Weighting Mean and Variability during Confidence Judgments.” PloS one 10(3):e0120870. Retrieved March 26, 2015 (http://www.pubmedcentral.nih.gov/articlerender.fcgi?artid=4368758&tool=pmcentrez&rendertype=abstract).

de Gardelle, Vincent and Christopher Summerfield. 2011. “Robust Averaging during Perceptual Judgment.” Proceedings of the National Academy of Sciences 108(32):13341–46. Retrieved October 3, 2015 (http://www.pnas.org/content/108/32/13341.abstract).

Geisler Wilson S. 2011. “Contributions of Ideal Observer Theory to Vision Research.” Vision Research 51(7):771–81.

Geisler Wilson S. and J. Najemnik. 2013. “Optimal and Non-Optimal Fixation Selection in Visual Search.” Perception ECVP abstract 42:226. Retrieved October 2, 2015 (http://www.perceptionweb.com/abstract.cgi?id=v130805).

Gekas N., M. Chalk, A. R. Seitz, and P. Serie. 2013. “Complexity and Specificity of Experimentally-Induced Expectations in Motion Perception.” Journal of Vision 13(4):8–8. Retrieved September 28, 2015 (http://jov.arvojournals.org/article.aspx?articleid=2121832).

Gepshtein, Sergei, Johannes Burge, Marc O. Ernst, and Martin S. Banks. 2005. “The Combination of Vision and Touch Depends on Spatial Proximity.” Journal of Vision 5(11):1013–23. Retrieved (http://eutils.ncbi.nlm.nih.gov/entrez/eutils/elink.fcgi?dbfrom=pubmed&id=16441199&retmode=ref&cmd=prlinks).

Gershman S. J., E. J. Horvitz, and Joshua B. Tenenbaum. 2015. “Computational Rationality: A Converging Paradigm for Intelligence in Brains, Minds, and Machines.” Science 349(6245):273–78. Retrieved July 17, 2015 (http://www.ncbi.nlm.nih.gov/pubmed/26185246).

Gibson J. J. and M. Radner. 1937. “Adaptation, after-Effect and Contrast in the Perception of Tilted Lines. I. Quantitative Studies.” Journal of experimental psychology 20(5):453–67. Retrieved (http://doi.apa.org/getdoi.cfm?doi=10.1037/h0059826).

Gigerenzer, Gerd and Henry Brighton. 2009. “Homo Heuristicus: Why Biased Minds Make Better Inferences.” Topics in Cognitive Science 1(1):107–43.

Gigerenzer, Gerd, U. Hoffrage, and H. Kleinbölting. 1991. “Probabilistic Mental Models: A Brunswikian Theory of Confidence.” Psychological review 98(4):506–28. Retrieved January 10, 2015 (http://www.ncbi.nlm.nih.gov/pubmed/1961771).

Gigerenzer, Gerd and Reinhard Selten. 2002. Bounded Rationality. Cambridge, MA: MIT Press.

Girshick Ahna R., Michael S. Landy, and Eero P. Simoncelli. 2011. “Cardinal Rules: Visual Orientation Perception Reflects Knowledge of Environmental Statistics.” Nature neuroscience 14(7):926–32. Retrieved August 20, 2015 (http://www.pubmedcentral.nih.gov/articlerender.fcgi?artid=3125404&tool=pmcentrez&rendertype=abstract).

Glennerster, Andrew, Lili Tcheang, Stuart J. Gilson, Andrew W. Fitzgibbon, and Andrew J. Parker. 2006. “Humans Ignore Motion and Stereo Cues in Favor of a Fictional Stable World.” Current Biology 16(4):428–32. Retrieved (http://eutils.ncbi.nlm.nih.gov/entrez/eutils/elink.fcgi?dbfrom=pubmed&id=16488879&retmode=ref&cmd=prlinks).

Gobell, Joetta and Marisa Carrasco. 2005. “Attention Alters the Appearance of Spatial Frequency and Gap Size.” Psychological science 16(8):644–51. Retrieved (http://eutils.ncbi.nlm.nih.gov/entrez/eutils/elink.fcgi?dbfrom=pubmed&id=16102068&retmode=ref&cmd=prlinks).

Gold Jason M., Richard F. Murray, Patrick J. Bennett, and Allison B. Sekuler. 2000. “Deriving Behavioural Receptive Fields for Visually Completed Contours.” Current Biology 10(11):663–66. Retrieved September 30, 2015 (http://www.sciencedirect.com/science/article/pii/S0960982200005236).

Goodman N. D. et al. 2015. “Relevant and Robust: A Response to Marcus and Davis (2013).” Psychological Science 26(4):539–41. Retrieved (http://pss.sagepub.com/lookup/doi/10.1177/0956797614559544).

Gorea, Andrei, Florent Caetta, and Dov Sagi. 2005. “Criteria Interactions across Visual Attributes.” Vision research 45(19):2523–32. Retrieved September 23, 2014 (http://www.ncbi.nlm.nih.gov/pubmed/15950255).

Gorea, Andrei and Dov Sagi. 2000. “Failure to Handle More than One Internal Representation in Visual Detection Tasks.” Proceedings of the National Academy of Sciences of the United States of America 97(22):12380–84. Retrieved September 23, 2014 (http://www.pubmedcentral.nih.gov/articlerender.fcgi?artid=17350&tool=pmcentrez&rendertype=abstract).

Gorea, Andrei and Dov Sagi. 2001. “Disentangling Signal from Noise in Visual Contrast Discrimination.” Nature neuroscience 4(11):1146–50. Retrieved September 23, 2014 (http://www.ncbi.nlm.nih.gov/pubmed/11687818).

Gorea, Andrei and Dov Sagi. 2002. “Natural Extinction: A Criterion Shift Phenomenon.” Visual Cognition 9(8):913–36.

Gori, Monica, Michela Del Viva, Giulio Sandini, and David C. Burr. 2008. “Young Children Do Not Integrate Visual and Haptic Form Information.” Current Biology 18(9):694–98. Retrieved (http://eutils.ncbi.nlm.nih.gov/entrez/eutils/elink.fcgi?dbfrom=pubmed&id=18450446&retmode=ref&cmd=prlinks).

Green D. M. and John A. Swets. 1966. Signal Detection Theory and Psychophysics. New York: John Wiley & Sons Ltd.

Griffin, Dale and Amos Tversky. 1992. “The Weighing of Evidence and the Determinants of Confidence.” Cognitive Psychology 24(3):411–35. Retrieved December 20, 2014 (http://www.sciencedirect.com/science/article/pii/001002859290013R).

Griffiths Thomas L., Nick Chater, Dennis Norris, and Alexandre Pouget. 2012. “How the Bayesians Got Their Beliefs (and What Those Beliefs Actually Are): Comment on Bowers and Davis (2012).” Psychological bulletin 138(3):415–22. Retrieved (http://doi.apa.org/getdoi.cfm?doi=10.1037/a0026884).

Griffiths Thomas L., Falk Lieder, and Noah D. Goodman. 2015. “Rational Use of Cognitive Resources: Levels of Analysis Between the Computational and the Algorithmic.” Topics in Cognitive Science 7(2):217–29. Retrieved (http://doi.wiley.com/10.1111/tops.12142).

Grzywacz Norberto M. and Rosario M. Balboa. 2002. “A Bayesian Framework for Sensory Adaptation.” Neural computation 14(3):543–59. Retrieved (http://eutils.ncbi.nlm.nih.gov/entrez/eutils/elink.fcgi?dbfrom=pubmed&id=11860682&retmode=ref&cmd=prlinks).

Gu, Yong, Dora E. Angelaki, and Gregory C. DeAngelis. 2008. “Neural Correlates of Multisensory Cue Integration in Macaque MSTd.” Nature Neuroscience 11(10):1201–10. Retrieved June 20, 2016 (http://www.nature.com/doifinder/10.1038/nn.2191).

Hammett Stephen T., Rebecca A. Champion, Peter G. Thompson, and Antony B. Morland. 2007. “Perceptual Distortions of Speed at Low Luminance: Evidence Inconsistent with a Bayesian Account of Speed Encoding.” Vision Research 47(4):564–68. Retrieved (http://www.ncbi.nlm.nih.gov/pubmed/17011014).

Hanks Timothy D., Mark E. Mazurek, Roozbeh Kiani, Elisabeth Hopp, and Michael N. Shadlen. 2011. “Elapsed Decision Time Affects the Weighting of Prior Probability in a Perceptual Decision Task.” The Journal of neuroscience 31(17):6339–52. Retrieved February 28, 2013 (http://www.pubmedcentral.nih.gov/articlerender.fcgi?artid=3356114&tool=pmcentrez&rendertype=abstract).

Hanks Timothy D. and Christopher Summerfield. 2017. “Perceptual Decision Making in Rodents, Monkeys, and Humans.” Neuron 93(1):15–31. Retrieved (http://dx.doi.org/10.1016/j.neuron.2016.12.003).

Harvey N. 1997. “Confidence in Judgment.” Trends in cognitive sciences 1(2):78–82. Retrieved (http://www.ncbi.nlm.nih.gov/pubmed/21223868).

Hassan, Omar and Stephen T. Hammett. 2015. “Perceptual Biases Are Inconsistent with Bayesian Encoding of Speed in the Human Visual System.” Journal of vision 15(2):9. Retrieved March 24, 2016 (http://jov.arvojournals.org/article.aspx?articleid=2213273).

Hawkins Guy E., Birte U. Forstmann, Eric-Jan Wagenmakers, Roger Ratcliff, and Scott D. Brown. 2015. “Revisiting the Evidence for Collapsing Boundaries and Urgency Signals in Perceptual Decision-Making.” Journal of Neuroscience 35(6):2476–84. Retrieved February 12, 2015 (http://www.jneurosci.org/content/35/6/2476.full).

Healy Alice F. and Michael Kubovy. 1981. “Probability Matching and the Formation of Conservative Decision Rules in a Numerical Analog of Signal Detection.” Journal of Experimental Psychology: Human Learning and Memory 7(5):344–54.

Heitz Richard P. 2014. “The Speed-Accuracy Tradeoff: History, Physiology, Methodology, and Behavior.” Frontiers in neuroscience 8:150. Retrieved May 19, 2015 (http://www.pubmedcentral.nih.gov/articlerender.fcgi?artid=4052662&tool=pmcentrez&rendertype=abstract).

Helmholtz H. L. F. 1856. Treatise on Physiological Optics. Bristol: Thoemmes Continuum.

Henriques, J. B., J. M. Glowacki, and R. J. Davidson. 1994. “Reward Fails to Alter Response Bias in Depression.” Journal of abnormal psychology 103(3):460–66. Retrieved December 26, 2014 (http://www.ncbi.nlm.nih.gov/pubmed/7930045).

Hillis J. M., Marc O. Ernst, Martin S. Banks, and Michael S. Landy. 2002. “Combining Sensory Information: Mandatory Fusion Within, but Not Between, Senses.” Science 298(5598):1627–30. Retrieved (http://www.sciencemag.org/cgi/doi/10.1126/science.1075396).

Hohwy, Jakob, Andreas Roepstorff, and Karl Friston. 2008. “Predictive Coding Explains Binocular Rivalry: An Epistemological Review.” Cognition 108(3):687–701. Retrieved (http://eutils.ncbi.nlm.nih.gov/entrez/eutils/elink.fcgi?dbfrom=pubmed&id=18649876&retmode=ref&cmd=prlinks).

Holmes, Philip and Jonathan D. Cohen. 2014. “Optimality and Some of Its Discontents: Successes and Shortcomings of Existing Models for Binary Decisions.” Topics in cognitive science 6(2):258–78. Retrieved November 6, 2014 (http://www.ncbi.nlm.nih.gov/pubmed/24648411).

Jack C. E. and W. R. Thurlow. 1973. “Effects of Degree of Visual Association and Angle of Displacement on the “ventriloquism” Effect.” Perceptual and motor skills 37(3):967–79. Retrieved (http://eutils.ncbi.nlm.nih.gov/entrez/eutils/elink.fcgi?dbfrom=pubmed&id=4764534&retmode=ref&cmd=prlinks).

Jacobs R. A. 1999. “Optimal Integration of Texture and Motion Cues to Depth.” Vision Research 39(21):3621–29. Retrieved (http://eutils.ncbi.nlm.nih.gov/entrez/eutils/elink.fcgi?dbfrom=pubmed&id=10746132&retmode=ref&cmd=prlinks).

Jastrow Joseph. 1892. “Studies from the University of Wisconsin: On the Judgment of Angles and Positions of Lines.” The American Journal of Psychology 5(2):214. Retrieved (http://www.jstor.org/stable/1410867?origin=crossref).

Jaynes Edwin T. 2003. Probability Theory: The Logic of Sicence. Cambridge University Press.

Jazayeri, Mehrdad and J. Anthony Movshon. 2007. “A New Perceptual Illusion Reveals Mechanisms of Sensory Decoding.” Nature 446(7138):912–15. Retrieved November 26, 2014 (http://www.nature.com/doifinder/10.1038/nature05739).

Jesteadt Walt. 1974. “Intensity and Frequency Discrimination in One- and Two-Interval Paradigms.” The Journal of the Acoustical Society of America 55(6):1266. Retrieved September 30, 2015 (http://scitation.aip.org/content/asa/journal/jasa/55/6/10.1121/1.1914696).

Jolij, Jacob and Victor A. F. Lamme. 2005. “Repression of Unconscious Information by Conscious Processing: Evidence from Affective Blindsight Induced by Transcranial Magnetic Stimulation.” Proceedings of the National Academy of Sciences of the United States of America 102(30):10747–51. Retrieved September 15, 2015 (http://www.pnas.org/content/102/30/10747.abstract).

Jones, Matt and Bradley C. Love. 2011. “Bayesian Fundamentalism or Enlightenment? On the Explanatory Status and Theoretical Contributions of Bayesian Models of Cognition.” Behavioral and Brain Sciences 34(4):169–88. Retrieved (http://www.journals.cambridge.org/abstract_S0140525X10003134).

Juslin, Peter, Håkan Nilsson, and Anders Winman. 2009. “Probability Theory, Not the Very Guide of Life.” Psychological Review 116(4):856–74.

Kahneman, Daniel and Amos Tversky. 1979. “Prospect Theory: An Analysis of Decision under Risk.” Econometrica 47(2):263. Retrieved March 11, 2017 (http://www.jstor.org/stable/1914185?origin=crossref).

Kalenscher, Tobias, Philippe N. Tobler, Willem Huijbers, Sander M. Daselaar, and Cyriel Pennartz. 2010. “Neural Signatures of Intransitive Preferences.” Frontiers in Human Neuroscience 4:49. Retrieved March 9, 2017 (http://journal.frontiersin.org/article/10.3389/fnhum.2010.00049/abstract).

Kaneko, Yoshiyuki and Katsuyuki Sakai. 2015. “Dissociation in Decision Bias Mechanism between Probabilistic Information and Previous Decision.” Frontiers in Human Neuroscience 9. Retrieved April 28, 2015 (http://journal.frontiersin.org/article/10.3389/fnhum.2015.00261/abstract).

Keren Gideon. 1988. “On the Ability of Monitoring Non-Veridical Perceptions and Uncertain Knowledge: Some Calibration Studies.” Acta Psychologica 67(2):95–119. Retrieved March 24, 2016 (http://www.sciencedirect.com/science/article/pii/0001691888900078).

Kiani, Roozbeh, Leah Corthell, and Michael N. Shadlen. 2014. “Choice Certainty Is Informed by Both Evidence and Decision Time.” Neuron 84(6):1329–42. Retrieved December 17, 2014 (http://www.sciencedirect.com/science/article/pii/S0896627314010964).

Kiani, Roozbeh and Michael N. Shadlen. 2009. “Representation of Confidence Associated with a Decision by Neurons in the Parietal Cortex.” Science 324(5928):759–64. Retrieved February 28, 2013 (http://www.pubmedcentral.nih.gov/articlerender.fcgi?artid=2738936&tool=pmcentrez&rendertype=abstract).

Kinchla R. A. and F. Smyzer. 1967. “A Diffusion Model of Perceptual Memory.” Perception & Psychophysics 2(6):219–29. Retrieved September 2, 2015 (http://www.springerlink.com/index/10.3758/BF03212471).

Knill David C. and Alexandre Pouget. 2004. “The Bayesian Brain: The Role of Uncertainty in Neural Coding and Computation.” Trends in neurosciences 27(12):712–19. Retrieved July 10, 2014 (http://www.ncbi.nlm.nih.gov/pubmed/15541511).

Knill David C. and Jeffrey A. Saunders. 2003. “Do Humans Optimally Integrate Stereo and Texture Information for Judgments of Surface Slant?” Vision Research 43(24):2539–58.

Koizumi, Ai, Brian Maniscalco, and Hakwan Lau. 2015. “Does Perceptual Confidence Facilitate Cognitive Control?” Attention, perception & psychophysics. Retrieved March 11, 2015 (http://www.ncbi.nlm.nih.gov/pubmed/25737256).

Körding Konrad P. et al. 2007. “Causal Inference in Multisensory Perception” edited by O. Sporns. PLoS ONE 2(9):e943. Retrieved June 20, 2016 (http://dx.plos.org/10.1371/journal.pone.0000943).

Körding Konrad P. and Daniel M. Wolpert. 2004. “Bayesian Integration in Sensorimotor Learning.” Nature 427(6971):244–47. Retrieved September 16, 2015 (http://dx.doi.org/10.1038/nature02169).

Körding Konrad P. and Daniel M. Wolpert. 2006. “Bayesian Decision Theory in Sensorimotor Control.” Trends in cognitive sciences 10(7):319–26. Retrieved July 2, 2015 (http://www.ncbi.nlm.nih.gov/pubmed/16807063).

Koriat Asher. 2011. “Subjective Confidence in Perceptual Judgments: A Test of the Self-Consistency Model.” Journal of Experimental Psychology: General 140(1):117–39. Retrieved March 18, 2013 (http://www.ncbi.nlm.nih.gov/pubmed/21299320).

Landy Michael S., Martin S. Banks, and David C. Knill. 2011. “Ideal-Observer Models of Cue Integration.” Pp. 5–29 in Sensory Cue Integration, edited by J. Trommershäuser, K. P. Körding, and M. S. Land. New York: Oxford University Press.

Landy Michael S., Ross Goutcher, Julia Trommershäuser, and Pascal Mamassian. 2007. “Visual Estimation under Risk.” Journal of vision 7(6):4. Retrieved November 26, 2014 (http://www.pubmedcentral.nih.gov/articlerender.fcgi?artid=2638507&tool=pmcentrez&rendertype=abstract).

Landy Michael S. and H. Kojima. 2001. “Ideal Cue Combination for Localizing Texture-Defined Edges.” Journal of the Optical Society of America A, Optics and image science 18(9):2307–20. Retrieved (http://www.cns.nyu.edu/~msl/papers/landykojima01.pdf).

Landy Michael S., L. Maloney, E. B. Johnston, and M. Youn. 1995. “Measurement and Modeling of Depth Cue Combination: In Defense of Weak Fusion.” Vision Research 35(3):389–412.

de Lange, Floris P., Dobromir Rahnev, Tobias H. Donner, and Hakwan Lau. 2013. “Prestimulus Oscillatory Activity over Motor Cortex Reflects Perceptual Expectations.” The Journal of neuroscience 33(4):1400–1410. Retrieved March 2, 2013 (http://www.ncbi.nlm.nih.gov/pubmed/23345216).

Langer M. S. and H. H. Bülthoff. 2001. “A Prior for Global Convexity in Local Shape-from-Shading.” Perception 30(4):403–10. Retrieved (http://eutils.ncbi.nlm.nih.gov/entrez/eutils/elink.fcgi?dbfrom=pubmed&id=11383189&retmode=ref&cmd=prlinks).

Lau, Hakwan and Richard E. Passingham. 2006. “Relative Blindsight in Normal Observers and the Neural Correlate of Visual Consciousness.” Proceedings of the National Academy of Sciences of the United States of America 103(49):18763–68. Retrieved (http://www.pubmedcentral.nih.gov/articlerender.fcgi?artid=1693736&tool=pmcentrez&rendertype=abstract).

Lau, Hakwan and David Rosenthal. 2011. “Empirical Support for Higher-Order Theories of Conscious Awareness.” Trends in cognitive sciences 15(8):365–73. Retrieved November 11, 2013 (http://www.ncbi.nlm.nih.gov/pubmed/21737339).

Lennie Peter. 2003. The Cost of Cortical Computation.

Leshowitz, Barry. 1969. “Comparison of ROC Curves from One- and Two-Interval Rating-Scale Procedures.” The Journal of the Acoustical Society of America 46(2B):399. Retrieved September 30, 2015 (http://scitation.aip.org/content/asa/journal/jasa/46/2B/10.1121/1.1911703).

Liberman, Alina, Jason Fischer, and David Whitney. 2014. “Serial Dependence in the Perception of Faces.” Current biology 24(21):2569–74. Retrieved October 8, 2014 (http://www.ncbi.nlm.nih.gov/pubmed/25283781).

Ling, Sam and Marisa Carrasco. 2006. “When Sustained Attention Impairs Perception.” Nature Neuroscience 9(10):1243–45. Retrieved (http://eutils.ncbi.nlm.nih.gov/entrez/eutils/elink.fcgi?dbfrom=pubmed&id=16964254&retmode=ref&cmd=prlinks).

Liu, Taosheng, Jared Abrams, and Marisa Carrasco. 2009. “Voluntary Attention Enhances Contrast Appearance.” Psychological science 20(3):354–62. Retrieved (http://eutils.ncbi.nlm.nih.gov/entrez/eutils/elink.fcgi?dbfrom=pubmed&id=19254239&retmode=ref&cmd=prlinks).

Lupyan Gary. 2012. “Linguistically Modulated Perception and Cognition: The Label-Feedback Hypothesis.” Frontiers in psychology 3:54. Retrieved (http://eutils.ncbi.nlm.nih.gov/entrez/eutils/elink.fcgi?dbfrom=pubmed&id=22408629&retmode=ref&cmd=prlinks).

Lupyan Gary. 2016. “The Paradox of the Universal Triangle: Concepts, Language, and Prototypes.” Quarterly journal of experimental psychology (2006) 1–69. Retrieved (http://eutils.ncbi.nlm.nih.gov/entrez/eutils/elink.fcgi?dbfrom=pubmed&id=26731302&retmode=ref&cmd=prlinks).

Luu, Long and Alan A. Stocker. 2016. “Choice-Induced Biases in Perception.” bioRxiv. Retrieved (http://biorxiv.org/content/early/2016/04/01/043224.abstract).

Ma, Wei Ji. 2010. “Signal Detection Theory, Uncertainty, and Poisson-like Population Codes.” Vision Research 50(22):2308–19. Retrieved November 26, 2014 (http://www.sciencedirect.com/science/article/pii/S004269891000430X).

Ma, Wei Ji, Jeffrey M. Beck, Peter E. Latham, and Alexandre Pouget. 2006. “Bayesian Inference with Probabilistic Population Codes.” Nature neuroscience 9(11):1432–38. Retrieved July 11, 2014 (http://www.ncbi.nlm.nih.gov/pubmed/17057707).

Macmillan, Neil A. and C. Douglas Creelman. 2005. Detection Theory: A User’s Guide. 2nd ed. Mahwah, NJ: Erlbaum.

Maddox, W.Todd. 1995. “Base-Rate Effects in Multidimensional Perceptual Categorization.” Journal of experimental psychology. Learning, memory, and cognition 21(2):288–301. Retrieved June 20, 2016 (http://www.ncbi.nlm.nih.gov/pubmed/7738501).

Maddox, W.Todd. 2002. “Toward a Unified Theory of Decision Criterion Learning in Perceptual Categorization.” Journal of the Experimental Analysis of Behavior 78(3):567–95. Retrieved June 20, 2016 (http://www.ncbi.nlm.nih.gov/pmc/articles/PMC1284916/).

Maddox, W.Todd and Corey J. Bohil. 1998a. “Base-Rate and Payoff Effects in Multidimensional Perceptual Categorization.” Journal of experimental psychology. Learning, memory, and cognition 24(6):1459–82.

Maddox, W.Todd and Corey J. Bohil. 1998b. “Overestimation of Base-Rate Differences in Complex Perceptual Categories.” Perception & psychophysics 60(4):575–92.

Maddox, W.Todd and Corey J. Bohil. 2000. “Costs and Benefits in Perceptual Categorization.” Memory & cognition 28(4):597–615.

Maddox, W.Todd and Corey J. Bohil. 2001. “Feedback Effects on Cost-Benefit Learning in Perceptual Categorization.” Memory & cognition 29(4):598–615.

Maddox, W.Todd and Corey J. Bohil. 2003. “A Theoretical Framework for Understanding the Effects of Simultaneous Base-Rate and Payoff Manipulations on Decision Criterion Learning in Perceptual Categorization.” Journal of experimental psychology. Learning, memory, and cognition 29(2):307–20.

Maddox, W.Todd and Corey J. Bohil. 2004. “Probability Matching, Accuracy Maximization, and a Test of the Optimal Classifier’s Independence Assumption in Perceptual Categorization.” Perception & psychophysics 66(1):104–18.

Maddox, W.Todd and Corey J. Bohil. 2005. “Optimal Classifier Feedback Improves Cost-Benefit but Not Base-Rate Decision Criterion Learning in Perceptual Categorization.” Memory & cognition 33(2):303–19.

Maddox, W.Todd, Corey J. Bohil, and Jeffrey L. Dodd. 2003. “Linear Transformations of the Payoff Matrix and Decision Criterion Learning in Perceptual Categorization.” Journal of experimental psychology. Learning, memory, and cognition 29(6):1174–93.

Maddox, W.Todd and J. L. Dod. 2001. “On the Relation between Base-Rate and Cost-Benefit Learning in Simulated Medical Diagnosis.” Journal of experimental psychology. Learning, memory, and cognition 27(6):1367–84. Retrieved December 31, 2014 (http://www.ncbi.nlm.nih.gov/pubmed/11713873).

Maiworm M. and B. Röder. 2011. “Suboptimal Auditory Dominance in Audiovisual Integration of Temporal Cues.” Tsinghua Science & Technology 16(2):121–32.

Maloney, Laurence T. and Michael S. Landy. 1989. “A Statistical Framework for Robust Fusion of Depth Information.” Pp. 1154–63 in Proceedings of SPIE, edited by W. A. Pearlman. Retrieved June 20, 2016 (http://proceedings.spiedigitallibrary.org/proceeding.aspx?articleid=1262206).

Maloney, Laurence T. and Pascal Mamassian. 2009. “Bayesian Decision Theory as a Model of Human Visual Perception: Testing Bayesian Transfer.” Visual neuroscience 26(1):147–55. Retrieved (http://www.ncbi.nlm.nih.gov/pubmed/19193251).

Maloney, Laurence T. and Hang Zhang. 2010. “Decision-Theoretic Models of Visual Perception and Action.” Vision research 50(23):2362–74. Retrieved June 20, 2016 (http://www.ncbi.nlm.nih.gov/pubmed/20932856).

Maniscalco, Brian and Hakwan Lau. 2010. “Comparing Signal Detection Models of Perceptual Decision Confidence.” Journal of Vision 10(7):213–213. Retrieved September 15, 2015 (http://jov.arvojournals.org/article.aspx?articleid=2138292).

Maniscalco, Brian and Hakwan Lau. 2012. “A Signal Detection Theoretic Approach for Estimating Metacognitive Sensitivity from Confidence Ratings.” Consciousness and cognition 21(1):422–30. Retrieved February 27, 2013 (http://www.ncbi.nlm.nih.gov/pubmed/22071269).

Maniscalco, Brian and Hakwan Lau. 2015. “Manipulation of Working Memory Contents Selectively Impairs Metacognitive Sensitivity in a Concurrent Visual Discrimination Task.” Neuroscience of Consciousness 2015(1):niv002. Retrieved September 8, 2015 (http://nc.oxfordjournals.org/content/2015/1/niv002.abstract).

Maniscalco, Brian and Hakwan Lau. 2016. “The Signal Processing Architecture Underlying Subjective Reports of Sensory Awareness.” Neuroscience of Consciousness (November 2015):1–17.

Maniscalco, Brian, Megan A. K. Peters, and Hakwan Lau. 2016. “Heuristic Use of Perceptual Evidence Leads to Dissociation between Performance and Metacognitive Sensitivity.” Attention, perception & psychophysics 78(3):923–37. Retrieved February 24, 2016 (http://www.ncbi.nlm.nih.gov/pubmed/26791233).

Marcus, Gary F. and Ernest Davis. 2013. “How Robust Are Probabilistic Models of Higher-Level Cognition?” Psychological science 24(12):2351–60. Retrieved August 21, 2014 (http://pss.sagepub.com/content/24/12/2351.abstract?ijkey=42fdf6a62d20a7c5e573d149a973e121f7ae2626&keytype2=tf_ipsecsha).

Marcus, Gary F. and Ernest Davis. 2015. “Still Searching for Principles: A Response to Goodman et Al. (2015).” Psychological Science 26(4):542–44. Retrieved (http://pss.sagepub.com/lookup/doi/10.1177/0956797614568433).

Markman Arthur B., Grant C. Baldwin, and W. Todd Maddox. 2005. “The Interaction of Payoff Structure and Regulatory Focus in Classification.” Psychological science 16(11):852–55. Retrieved December 30, 2014 (http://www.ncbi.nlm.nih.gov/pubmed/16262768).

Markowitz, Joseph and John A. Swets. 1967. “Factors Affecting the Slope of Empirical ROC Curves: Comparison of Binary and Rating Responses.” Perception & Psychophysics 2(3):91–100. Retrieved September 30, 2015 (http://www.springerlink.com/index/10.3758/BF03210301).

Massoni Sébastien. 2014. “Emotion as a Boost to Metacognition: How Worry Enhances the Quality of Confidence.” Consciousness and cognition 29:189–98. Retrieved October 7, 2014 (http://www.ncbi.nlm.nih.gov/pubmed/25286128).

Massoni, Sébastien, Thibault Gajdos, and Jean-Christophe Vergnaud. 2014. “Confidence Measurement in the Light of Signal Detection Theory.” Frontiers in Psychology 5:1455. Retrieved January 18, 2018 (http://www.ncbi.nlm.nih.gov/pubmed/25566135).

McCurdy, Li Yan et al. 2013. “Anatomical Coupling between Distinct Metacognitive Systems for Memory and Visual Perception.” The Journal of neuroscience 33(5):1897–1906. Retrieved March 5, 2013 (http://www.ncbi.nlm.nih.gov/pubmed/23365229).

Metcalfe, Janet and Arthur P. Shimamura. 1994. Metacognition: Knowing about Knowing. Cambridge, MA: MIT Press.

Michael, Elizabeth, Vincent de Gardelle, Alejo Nevado-Holgado, and Christopher Summerfield. 2015. “Unreliable Evidence: 2 Sources of Uncertainty During Perceptual Choice.” Cerebral Cortex 25(4):937–47. Retrieved November 22, 2013 (http://www.ncbi.nlm.nih.gov/pubmed/24122138).

Michael, Elizabeth, Vincent de Gardelle, and Christopher Summerfield. 2014. “Priming by the Variability of Visual Information.” Proceedings of the National Academy of Sciences 111(21):7873–78. Retrieved May 14, 2014 (http://www.ncbi.nlm.nih.gov/pubmed/24821803).

Morales, Jorge et al. 2015. “Low Attention Impairs Optimal Incorporation of Prior Knowledge in Perceptual Decisions.” Attention, perception & psychophysics 77(6):2021–36. Retrieved August 28, 2015 (http://www.ncbi.nlm.nih.gov/pubmed/25836765).

Mozer Michael C., Harold Pashler, and Hadjar Homaei. 2008. “Optimal Predictions in Everyday Cognition: The Wisdom of Individuals or Crowds?” Cognitive science 32(7):1133–47.

Mueller S. T. and C. T. Weideman. 2008. “Decision Noise: An Explanation for Observed Violations of Signal Detection Theory.” Psychonomic Bulletin & Review 15(3):465–94. Retrieved May 14, 2013 (http://www.springerlink.com/index/10.3758/PBR.15.3.465).

Nardini, Marko, Rachael Bedford, and Denis Mareschal. 2010. “Fusion of Visual Cues Is Not Mandatory in Children.” Proceedings of the National Academy of Sciences 107(39):17041–46. Retrieved (http://eutils.ncbi.nlm.nih.gov/entrez/eutils/elink.fcgi?dbfrom=pubmed&id=20837526&retmode=ref&cmd=prlinks).

Nardini, Marko, Peter Jones, Rachael Bedford, and Oliver Braddick. 2008. “Development of Cue Integration in Human Navigation.” Current Biology 18(9):689–93. Retrieved (http://eutils.ncbi.nlm.nih.gov/entrez/eutils/elink.fcgi?dbfrom=pubmed&id=18450447&retmode=ref&cmd=prlinks).

Navajas, Joaquin et al. 2017. “The Idiosyncratic Nature of Confidence.” Nature Human Behaviour 1(11):810–18. Retrieved November 14, 2017 (http://www.nature.com/articles/s41562-017-0215-1).

Navajas, Joaquin, Mariano Sigman, and Juan E. Kamienkowski. 2014. “Dynamics of Visibility, Confidence, and Choice during Eye Movements.” Journal of experimental psychology: Human perception and performance 40(3):1213–27. Retrieved April 18, 2014 (http://www.ncbi.nlm.nih.gov/pubmed/24730743).

Norton Elyse H., Stephen M. Fleming, Nathaniel D. Daw, and Michael S. Landy. 2017. “Suboptimal Criterion Learning in Static and Dynamic Environments.” PLoS computational biology 13(1):e1005304.

Odegaard, Brian, David R. Wozny, and Ladan Shams. 2015. “Biases in Visual, Auditory, and Audiovisual Perception of Space.” PLoS computational biology 11(12):e1004649. Retrieved January 8, 2016 (http://journals.plos.org/ploscompbiol/article?id=10.1371/journal.pcbi.1004649).

Olzak L. A. 1985. “Interactions between Spatially Tuned Mechanisms: Converging Evidence.” Journal of the Optical Society of America. A, Optics and image science 2(9):1551–9.

Oruç, Ipek, Laurence T. Maloney, and Michael S. Landy. 2003. “Weighted Linear Cue Combination with Possibly Correlated Error.” Vision Research 43(23):2451–68. Retrieved (http://eutils.ncbi.nlm.nih.gov/entrez/eutils/elink.fcgi?dbfrom=pubmed&id=12972395&retmode=ref&cmd=prlinks).

Osgood, Charles Egerton. 1953. Method and Theory in Experimental Psychology. Oxford University Press.

Oud, Bastiaan et al. 2016. “Irrational Time Allocation in Decision-Making.” Proceedings of the Royal Society B: Biological Sciences 283(1822):20151439. Retrieved (http://www.ncbi.nlm.nih.gov/pubmed/26763695).

Peters, Megan A. K., Wei Ji Ma, and Ladan Shams. 2016. “The Size-Weight Illusion Is Not Anti-Bayesian after All: A Unifying Bayesian Account.” PeerJ 4:e2124. Retrieved June 6, 2017 (http://www.ncbi.nlm.nih.gov/pubmed/27350899).

Petrini, Karin, Alicia Remark, Louise Smith, and Marko Nardini. 2014. “When Vision Is Not an Option: Children’s Integration of Auditory and Haptic Information Is Suboptimal.” Developmental science 17(3):376–87. Retrieved (http://onlinelibrary.wiley.com/doi/10.1111/desc.12127/full).

Petzschner, Frederike H. and Stefan Glasauer. 2011. “Iterative Bayesian Estimation as an Explanation for Range and Regression Effects: A Study on Human Path Integration.” Journal of Neuroscience 31(47). Retrieved July 13, 2017 (http://www.jneurosci.org/content/31/47/17220).

Plaisier M. A., L. C. J. van Dam, C. Glowania, and M. O. Erns. 2014. “Exploration Mode Affects Visuohaptic Integration of Surface Orientation.” Journal of Vision 14(13):22–22. Retrieved January 18, 2018 (http://www.ncbi.nlm.nih.gov/pubmed/25413627).

Pleskac, Timothy J. and Jerome R. Busemeyer. 2010. “Two-Stage Dynamic Signal Detection: A Theory of Choice, Decision Time, and Confidence.” Psychological review 117(3):864–901. Retrieved March 2, 2013 (http://www.ncbi.nlm.nih.gov/pubmed/20658856).

Prsa, Mario, Steven Gale, and Olaf Blanke. 2012. “Self-Motion Leads to Mandatory Cue Fusion across Sensory Modalities.” Journal of Neurophysiology 108(8):2282–91. Retrieved (http://eutils.ncbi.nlm.nih.gov/entrez/eutils/elink.fcgi?dbfrom=pubmed&id=22832567&retmode=ref&cmd=prlinks).

Pynn C. T. 1972. “Intensity Perception. III. Resolution in Small-Range Identification.” The Journal of the Acoustical Society of America 51(2B):559. Retrieved September 30, 2015 (http://scitation.aip.org/content/asa/journal/jasa/51/2B/10.1121/1.1912878).

Rahnev, Dobromir, Brian Maniscalco, et al. 2011. “Attention Induces Conservative Subjective Biases in Visual Perception.” Nature Neuroscience 14(12):1513–15. Retrieved (http://www.ncbi.nlm.nih.gov/pubmed/22019729).

Rahnev, Dobromir et al. 2013. “Continuous Theta Burst Transcranial Magnetic Stimulation Reduces Resting State Connectivity between Visual Areas.” Journal of neurophysiology 110(8):1811–21. Retrieved September 30, 2013 (http://www.ncbi.nlm.nih.gov/pubmed/23883858).

Rahnev, Dobromir, Linda Bahdo, Floris P. de Lange, and Hakwan Lau. 2012. “Prestimulus Hemodynamic Activity in Dorsal Attention Network Is Negatively Associated with Decision Confidence in Visual Perception.” Journal of neurophysiology 108(5):1529–36. Retrieved April 5, 2013 (http://www.ncbi.nlm.nih.gov/pubmed/22723670).

Rahnev, Dobromir, Ai Koizumi, Li Yan McCurdy, Mark D’Esposito, and Hakwan Lau. 2015. “Confidence Leak in Perceptual Decision Making.” Psychological Science 26(11):1664–80. Retrieved (http://pss.sagepub.com/lookup/doi/10.1177/0956797615595037).

Rahnev, Dobromir, Hakwan Lau, and Floris P. De Lange. 2011. “Prior Expectation Modulates the Interaction between Sensory and Prefrontal Regions in the Human Brain.” Journal of Neuroscience 31(29):10741–48.

Rahnev, Dobromir, Brian Maniscalco, Bruce Luber, Hakwan Lau, and Sarah H. Lisanby. 2012. “Direct Injection of Noise to the Visual Cortex Decreases Accuracy but Increases Decision Confidence.” Journal of neurophysiology 107(6):1556–63. Retrieved March 15, 2013 (http://www.ncbi.nlm.nih.gov/pubmed/22170965).

Rahnev, Dobromir, Derek Evan Nee, Justin Riddle, Alina Sue Larson, and Mark D’Esposito. 2016. “Causal Evidence for Frontal Cortex Organization for Perceptual Decision Making.” Proceedings of the National Academy of Sciences 113(20):6059–6064. Retrieved May 10, 2016 (http://www.pnas.org/content/early/2016/05/04/1522551113.full?tab=metrics).

Ramachandran V. 1990. “Interactions between Motion, Depth, Color and Form: The Utilitarian Theory of Perception.” Pp. 346–360 in Vision: Coding and efficiency, edited by C. Blakemore. Cambridge, England: Cambridge University Press.

Ratcliff, Roger and Jeffrey J. Starns. 2009. “Modeling Confidence and Response Time in Recognition Memory.” Psychological review 116(1):59–83. Retrieved July 22, 2014 (http://www.pubmedcentral.nih.gov/articlerender.fcgi?artid=2693899&tool=pmcentrez&rendertype=abstract).

Rauber H. J. and S. Treu. 1998. “Reference Repulsion When Judging the Direction of Visual Motion.” Perception 27(4):393–402. Retrieved (http://eutils.ncbi.nlm.nih.gov/entrez/eutils/elink.fcgi?dbfrom=pubmed&id=9797918&retmode=ref&cmd=prlinks).

Raviv, Ofri, Merav Ahissar, and Yonatan Loewenstein. 2012. “How Recent History Affects Perception: The Normative Approach and Its Heuristic Approximation.” PLoS computational biology 8(10):e1002731. Retrieved (http://eutils.ncbi.nlm.nih.gov/entrez/eutils/elink.fcgi?dbfrom=pubmed&id=23133343&retmode=ref&cmd=prlinks).

Reckless, Greg E. et al. 2014. “The Left Inferior Frontal Gyrus Is Involved in Adjusting Response Bias during a Perceptual Decision-Making Task.” Brain and behavior 4(3):398–407. Retrieved December 26, 2014 (http://www.pubmedcentral.nih.gov/articlerender.fcgi?artid=4055190&tool=pmcentrez&rendertype=abstract).

Reckless Greg E., Ingeborg Bolstad, Per H. Nakstad, Ole a Andreassen, and Jimmy Jensen. 2013. “Motivation Alters Response Bias and Neural Activation Patterns in a Perceptual Decision-Making Task.” Neuroscience 238:135–47. Retrieved March 5, 2013 (http://www.ncbi.nlm.nih.gov/pubmed/23428623).

Regenwetter, Michel et al. 2017. “Heterogeneity and Parsimony in Intertemporal Choice.” Decision. Retrieved March 9, 2017 (http://doi.apa.org/getdoi.cfm?doi=10.1037/dec0000069).

Regenwetter, Michel, Jason Dana, and Clintin P. Davis-Stober. 2010. “Testing Transitivity of Preferences on Two-Alternative Forced Choice Data.” Frontiers in Psychology 1:148. Retrieved March 9, 2017 (http://journal.frontiersin.org/article/10.3389/fpsyg.2010.00148/abstract).

Regenwetter, Michel, Jason Dana, Clintin P. Davis-Stober, and Ying Guo. 2011. “Parsimonious Testing of Transitive or Intransitive Preferences: Reply to Birnbaum (2011).” Psychological Review 118(4):684–88. Retrieved March 9, 2017 (http://doi.apa.org/getdoi.cfm?doi=10.1037/a0025291).

Renart, Alfonso and Christian K. Machens. 2014. “Variability in Neural Activity and Behavior.” Current Opinion in Neurobiology 25:211–20. Retrieved (http://dx.doi.org/10.1016/j.conb.2014.02.013).

Roach Neil W., James Heron, and Paul V McGraw. 2006. “Resolving Multisensory Conflict: A Strategy for Balancing the Costs and Benefits of Audio-Visual Integration.” Proceedings Biological sciences / The Royal Society 273(1598):2159–68. Retrieved (http://eutils.ncbi.nlm.nih.gov/entrez/eutils/elink.fcgi?dbfrom=pubmed&id=16901835&retmode=ref&cmd=prlinks).

van Rooij, Iris. 2008. “The Tractable Cognition Thesis.” Cognitive Science 32(6):939–84. Retrieved December 21, 2017 (http://doi.wiley.com/10.1080/03640210801897856).

Rosas, Pedro, Johan Wagemans, Marc O. Ernst, and Felix A. Wichmann. 2005. “Texture and Haptic Cues in Slant Discrimination: Reliability-Based Cue Weighting without Statistically Optimal Cue Combination.” Journal of the Optical Society of America A, Optics and image science 22(5):801–9.

Rosas, Pedro and Felix A. Wichmann. 2011. “Cue Combination: Beyond Optimality.” Pp. 144–52 in Sensory Cue Integration, edited by J. Trommershäuser, K. P. Körding, and M. S. Land. New York: Oxford University Press.

Rosas, Pedro, Felix A. Wichmann, and Johan Wagemans. 2007. “Texture and Object Motion in Slant Discrimination: Failure of Reliability-Based Weighting of Cues May Be Evidence for Strong Fusion.” Journal of Vision 7(6):3. Retrieved (http://eutils.ncbi.nlm.nih.gov/entrez/eutils/elink.fcgi?dbfrom=pubmed&id=17685786&retmode=ref&cmd=prlinks).

Saarela, Toni P. and Michael S. Landy. 2015. “Integration Trumps Selection in Object Recognition.” Current Biology 25(7):920–27. Retrieved March 24, 2016 (http://www.ncbi.nlm.nih.gov/pubmed/25802154).

Sabra A. I. 1989. “The Optics of Ibn Al-Haytham. Books I-III. On Direct Vision.”.

Samaha, Jason, John J. Barrett, Andrew D. Sheldon, Joshua J. LaRocque, and Bradley R. Postle. 2016. “Dissociating Perceptual Confidence from Discrimination Accuracy Reveals No Influence of Metacognitive Awareness on Working Memory.” Frontiers in Psychology 7:851. Retrieved August 2, 2016 (http://journal.frontiersin.org/Article/10.3389/fpsyg.2016.00851/abstract).

Sanders Joshua I., Balázs Hangya, and Adam Kepecs. 2016. “Signatures of a Statistical Computation in the Human Sense of Confidence.” Neuron 90(3):499–506. Retrieved May 5, 2016 (http://www.cell.com/article/S0896627316300162/fulltext).

Schulman A. I. and R. R. Mitchel. 1966. “Operating Characteristics from Yes-No and Forced-Choice Procedures.” The Journal of the Acoustical Society of America 40(2):473–77. Retrieved September 30, 2015 (http://www.ncbi.nlm.nih.gov/pubmed/5911357).

Schurger, Aaron, Min-Soo Kim, and Jonathan D. Cohen. 2015. “Paradoxical Interaction between Ocular Activity, Perception, and Decision Confidence at the Threshold of Vision.” PloS one 10(5):e0125278. Retrieved August 3, 2015 (http://journals.plos.org/plosone/article?id=10.1371/journal.pone.0125278).

Schwiedrzik, Caspar M. et al. 2014. “Untangling Perceptual Memory: Hysteresis and Adaptation Map into Separate Cortical Networks.” Cerebral Cortex 24(5):1152–64. Retrieved (http://eutils.ncbi.nlm.nih.gov/entrez/eutils/elink.fcgi?dbfrom=pubmed&id=23236204&retmode=ref&cmd=prlinks).

See Judi E., Joel S. Warm, William N. Dember, and Steven R. Howe. 1997. “Vigilance and Signal Detection Theory: An Empirical Evaluation of Five Measures of Response Bias.” Human Factors 39(1):14–29. Retrieved June 17, 2016 (http://hfs.sagepub.com/cgi/doi/10.1518/001872097778940704).

Seriès, Peggy and Aaron R. Seitz. 2013. “Learning What to Expect (in Visual Perception).” Frontiers in human neuroscience 7:668. Retrieved (http://eutils.ncbi.nlm.nih.gov/entrez/eutils/elink.fcgi?dbfrom=pubmed&id=24187536&retmode=ref&cmd=prlinks).

Shen, Shan and Wei Ji Ma. 2016. “A Detailed Comparison of Optimality and Simplicity in Perceptual Decision Making.” Psychological review. Retrieved June 13, 2016 (http://www.ncbi.nlm.nih.gov/pubmed/27177259).

Sherman M. T., A. K. Seth, A. B. Barrett, and R. Kana. 2015. “Prior Expectations Facilitate Metacognition for Perceptual Decision.” Consciousness and cognition 35:53–65. Retrieved May 21, 2015 (http://www.sciencedirect.com/science/article/pii/S1053810015000926).

Simen, Patrick et al. 2009. “Reward Rate Optimization in Two-Alternative Decision Making: Empirical Tests of Theoretical Predictions.” Journal of experimental psychology. Human perception and performance 35(6):1865–97. Retrieved September 11, 2015 (http://www.pubmedcentral.nih.gov/articlerender.fcgi?artid=2791916&tool=pmcentrez&rendertype=abstract).

Simon H. A. 1956. “Rational Choice and the Structure of the Environment.” Psychological Review 63(2):129–38.

Simon Herbert. 1957. “A Behavioral Model of Rational Choice.” in Models of Man, Social and Rational: Mathematical Essays on Rational Human Behavior in a Social Setting. New York: Wiley.

Snyder Joel S., Caspar M. Schwiedrzik, A. Davi Vitela, and Lucia Melloni. 2015. “How Previous Experience Shapes Perception in Different Sensory Modalities.” Frontiers in human neuroscience 9:594. Retrieved (http://eutils.ncbi.nlm.nih.gov/entrez/eutils/elink.fcgi?dbfrom=pubmed&id=26582982&retmode=ref&cmd=prlinks).

Solovey, Guillermo, Guy Gerard Graney, and Hakwan Lau. 2015. “A Decisional Account of Subjective Inflation of Visual Perception at the Periphery.” Attention, perception & psychophysics 77(1):258–71. Retrieved August 3, 2015 (http://www.ncbi.nlm.nih.gov/pubmed/25248620).

Song, Amanda, Ai Koizumi, and Hakwan Lau. 2015. “A Behavioral Method to Manipulate Metacognitive Awareness Independent of Stimulus Awareness.” in Behavioral Methods in Consciousness Research, edited by M. Overgaard. Oxford: Oxford University Press.

Song, Chen et al. 2011. “Relating Inter-Individual Differences in Metacognitive Performance on Different Perceptual Tasks.” Consciousness and cognition 20(4):1787–92. Retrieved March 7, 2013 (http://www.pubmedcentral.nih.gov/articlerender.fcgi?artid=3203218&tool=pmcentrez&rendertype=abstract).

Spence Morgan L., Paul E. Dux, and Derek H. Arnold. 2016. “Computations Underlying Confidence in Visual Perception.” Journal of experimental psychology. Human perception and performance 42(5):671–82. Retrieved December 3, 2015 (http://www.ncbi.nlm.nih.gov/pubmed/26594876).

Starns, Jeffrey J. and Roger Ratcliff. 2010. “The Effects of Aging on the Speed-Accuracy Compromise: Boundary Optimality in the Diffusion Model.” Psychology and aging 25(2):377–90. Retrieved September 11, 2015 (http://www.pubmedcentral.nih.gov/articlerender.fcgi?artid=2896207&tool=pmcentrez&rendertype=abstract).

Starns, Jeffrey J. and Roger Ratcliff. 2012. “Age-Related Differences in Diffusion Model Boundary Optimality with Both Trial-Limited and Time-Limited Tasks.” Psychonomic bulletin & review 19(1):139–45. Retrieved September 11, 2015 (http://www.ncbi.nlm.nih.gov/pubmed/22144142).

Stocker, Alan A. and Eero P. Simoncelli. 2006a. “Noise Characteristics and Prior Expectations in Human Visual Speed Perception.” Nature neuroscience 9(4):578–85. Retrieved November 24, 2014 (http://dx.doi.org/10.1038/nn1669).

Stocker, Alan A. and Eero P. Simoncelli. 2006b. “Sensory Adaptation within a Bayesian Framework for Perception.” Advances in Neural Information Processing Systems 18 18:1291–98.

Stocker, Alan A. and Eero P. Simoncelli. 2008. “A Bayesian Model of Conditioned Perception.” Pp. 1409–16 in Advances in Neural Information Processing Systems, vol. 20, edited by J. C. Platt, D. Koller, Y. Singer, and S. Roweis. Cambridge, MA: MIT Press.

Stone L. S. and Peter Thompson. 1992. “Human Speed Perception Is Contrast Dependent.” Vision research 32(8):1535–49. Retrieved September 28, 2015 (http://www.ncbi.nlm.nih.gov/pubmed/1455726).

Störmer Viola S., John J. Mcdonald, and Steven A. Hillyard. 2009. “Cross-Modal Cueing of Attention Alters Appearance and Early Cortical Processing of Visual Stimuli.” Proceedings of the National Academy of Sciences 106(52):22456–61. Retrieved (http://eutils.ncbi.nlm.nih.gov/entrez/eutils/elink.fcgi?dbfrom=pubmed&id=20007778&retmode=ref&cmd=prlinks).

Summerfield, Christopher and Etienne Koechlin. 2010. “Economic Value Biases Uncertain Perceptual Choices in the Parietal and Prefrontal Cortices.” Frontiers in human neuroscience 4:208. Retrieved September 2, 2015 (http://www.pubmedcentral.nih.gov/articlerender.fcgi?artid=3024559&tool=pmcentrez&rendertype=abstract).

Summerfield, Christopher and Konstantinos Tsetsos. 2015. “Do Humans Make Good Decisions?” Trends in Cognitive Sciences 19(1):27–34. Retrieved September 5, 2015 (http://www.sciencedirect.com/science/article/pii/S1364661314002381).

Sun J. and P. Peron. 1997. “Shading and Stereo in Early Perception of Shape and Reflectance.” Perception 26(4):519–29. Retrieved (http://eutils.ncbi.nlm.nih.gov/entrez/eutils/elink.fcgi?dbfrom=pubmed&id=9404497&retmode=ref&cmd=prlinks).

Swets, John A. and D. M. Green. 1961. “Sequential Observations by Human Observers of Signals in Noise.” Pp. 177–95 in Information theory: Proceedings of the fourth London symposium, edited by C. Cherry. London: Butterworth.

Swets, John A., W. P. Tanner, and T. G. Birdsall. 1961. “Decision Processes in Perception.” Psychological review 68(5):301–40. Retrieved July 1, 2015 (http://www.ncbi.nlm.nih.gov/pubmed/13774292).

Tanner T. A., R. W. Haller, and R. C. Atkinson. 1967. “Signal Recognition as Influenced by Presentation Schedules.” Perception & Psychophysics 2(8):349–58. Retrieved September 4, 2015 (http://www.springerlink.com/index/10.3758/BF03210070).

Tanner W. P. 1956. “Theory of Recognition.” Journal of the Acoustical Society of America 28:882–88.

Tanner W. P. 1961. “Physiological Implications of Psychophysical Data.” Annals of the New York Academy of Sciences 89:752–65. Retrieved September 30, 2015 (http://www.ncbi.nlm.nih.gov/pubmed/13775211).

Tauber, Sean, Daniel J. Navarro, Amy Perfors, and Mark Steyvers. 2017. “Bayesian Models of Cognition Revisited: Setting Optimality aside and Letting Data Drive Psychological Theory.” Psychological Review 124(4):410–41.

Taylor, Stephan F. et al. 2004. “A Functional Neuroimaging Study of Motivation and Executive Function.” NeuroImage 21(3):1045–54. Retrieved December 16, 2014 (http://www.ncbi.nlm.nih.gov/pubmed/15006672).

Tenenbaum, Joshua B. and Thomas L. Griffiths. 2006. “Optimal Predictions in Everyday Cognition.” Psychological Science 17(9):767–73.

Tenenbaum Joshua B., Charles Kemp, Thomas L. Griffiths, and Noah D. Goodman. 2011. “How to Grow a Mind: Statistics, Structure, and Abstraction.” Science 331(6022):1279–85. Retrieved February 28, 2013 (http://www.ncbi.nlm.nih.gov/pubmed/21393536).

Thompson Peter. 1982. “Perceived Rate of Movement Depends on Contrast.” Vision research 22(3):377–80. Retrieved September 28, 2015 (http://www.ncbi.nlm.nih.gov/pubmed/7090191).

Thompson, Peter, Kevin Brooks, and Stephen T. Hammett. 2006. “Speed Can Go up as Well as down at Low Contrast: Implications for Models of Motion Perception.” Vision Research 46(6–7):782–86. Retrieved (http://eutils.ncbi.nlm.nih.gov/entrez/eutils/elink.fcgi?dbfrom=pubmed&id=16171842&retmode=ref&cmd=prlinks).

Thura, David, Julie Beauregard-Racine, Charles-William Fradet, and Paul Cisek. 2012. “Decision Making by Urgency Gating: Theory and Experimental Support.” Journal of neurophysiology 108(11):2912–30. Retrieved December 14, 2014 (http://www.ncbi.nlm.nih.gov/pubmed/22993260).

Treisman, Michel and Andrew Faulkner. 1984. “The Setting and Maintenance of Criteria Representing Levels of Confidence.” Journal of Experimental Psychology: Human Perception and Performance 10(1):119–39. Retrieved September 15, 2015 (http://discovery.ucl.ac.uk/20033/).

Trommershäuser Julia. 2009. “Biases and Optimality of Sensory-Motor and Cognitive Decisions.” Progress in brain research 174:267–78. Retrieved (http://eutils.ncbi.nlm.nih.gov/entrez/eutils/elink.fcgi?dbfrom=pubmed&id=19477345&retmode=ref&cmd=prlinks).

Trommershäuser, Julia, Konrad P. Körding, and Michael S. Landy, eds. 2011. Sensory Cue Integration. New York: Oxford University Press.

Tse, Peter U. 2005. “Voluntary Attention Modulates the Brightness of Overlapping Transparent Surfaces.” Vision Research 45(9):1095–98. Retrieved (http://eutils.ncbi.nlm.nih.gov/entrez/eutils/elink.fcgi?dbfrom=pubmed&id=15707917&retmode=ref&cmd=prlinks).

Tsetsos, Konstantinos et al. 2016. “Economic Irrationality Is Optimal during Noisy Decision Making.” Proceedings of the National Academy of Sciences of the United States of America 113(11):3102–7. Retrieved March 2, 2016 (http://www.pnas.org/content/early/2016/02/24/1519157113.long).

Tsetsos, Konstantinos, Thomas Pfeffer, Pia Jentgens, and Tobias H. Donner. 2015. “Action Planning and the Timescale of Evidence Accumulation.” PloS one 10(6):e0129473.

Tsotsos, John K. 1993. “The Role of Computational Complexity in Perceptual Theory.” Advances in Psychology 99:261–96.

Turatto, Massimo, Massimo Vescovi, and Matteo Valsecchi. 2007. “Attention Makes Moving Objects Be Perceived to Move Faster.” Vision Research 47(2):166–78. Retrieved (http://eutils.ncbi.nlm.nih.gov/entrez/eutils/elink.fcgi?dbfrom=pubmed&id=17116314&retmode=ref&cmd=prlinks).

Turnbull W. H. 1961. The Correspondence of Isaac Newton, Vol. 3, 1688–1694. Cambridge: Cambridge University Press.

Ulehla Z. J. 1966. “Optimality of Perceptual Decision Criteria.” Journal of experimental psychology 71(4):564–69. Retrieved December 30, 2014 (http://www.ncbi.nlm.nih.gov/pubmed/5909083).

Vandormael, Hildward, Santiago Herce Castañón, Jan Balaguer, Vickie Li, and Christopher Summerfield. 2017. “Robust Sampling of Decision Information during Perceptual Choice.” Proceedings of the National Academy of Sciences 114(10):2771–76. Retrieved (http://www.pnas.org/lookup/doi/10.1073/pnas.1613950114).

Varey C. A., B. A. Mellers, and M. H. Birnbaum. 1990. “Judgments of Proportions.” Journal of experimental psychology. Human perception and performance 16(3):613–25. Retrieved October 2, 2015 (http://www.ncbi.nlm.nih.gov/pubmed/2144575).

Vaziri-Pashkam M. and P. Cavanagh. 2008. “Apparent Speed Increases at Low Luminance.” Journal of Vision 8(16):9. Retrieved (http://www.ncbi.nlm.nih.gov/pubmed/19146275).

Vickers D. 1979. Decision Processes in Visual Perception. New York: Academic Press.

Vickers, D. and J. Packer. 1982. “Effects of Alternating Set for Speed or Accuracy on Response Time, Accuracy and Confidence in a Unidimensional Discrimination Task.” Acta psychologica 50(2):179–97.

Viemeister N. F. 1970. “Intensity Discrilnination: Performance in Three Paradigms.” Perception & Psychophysics 8(6):417–19. Retrieved September 30, 2015 (http://www.springerlink.com/index/10.3758/BF03207037).

Vincent B. 2011. “Covert Visual Search: Prior Beliefs Are Optimally Combined with Sensory Evidence.” Journal of Vision 11(13):25–25.

Vintch, Brett and Justin L. Gardner. 2014. “Cortical Correlates of Human Motion Perception Biases.” Journal of Neuroscience 34(7):2592–2604. Retrieved (http://eutils.ncbi.nlm.nih.gov/entrez/eutils/elink.fcgi?dbfrom=pubmed&id=24523549&retmode=ref&cmd=prlinks).

Vlassova A., C. Donkin, and J. Pearson. 2014. “Unconscious Information Changes Decision Accuracy but Not Confidence.” Proceedings of the National Academy of Sciences 111(45):16214–16218. Retrieved October 28, 2014 (http://www.pnas.org/content/early/2014/10/24/1403619111.short).

Vul, Edward, Noah Goodman, Thomas L. Griffiths, and Joshua B. Tenenbaum. 2014. “One and Done? Optimal Decisions from Very Few Samples.” Cognitive science 38(4):599–637. Retrieved February 17, 2015 (http://www.ncbi.nlm.nih.gov/pubmed/24467492).

Wainwright M. J. 1999. “Visual Adaptation as Optimal Information Transmission.” Vision Research 39(23):3960–74. Retrieved (http://eutils.ncbi.nlm.nih.gov/entrez/eutils/elink.fcgi?dbfrom=pubmed&id=10748928&retmode=ref&cmd=prlinks).

Ward, Lawrence M. and G. R. Lockhead. 1970. “Sequential Effects and Memory in Category Judgments.” Journal of Experimental Psychology 84(1):27–34. Retrieved September 15, 2015 (https://scholars.duke.edu/display/pub651252).

Wark, Barry, Brian Nils Lundstrom, and Adrienne Fairhall. 2007. “Sensory Adaptation.” Current Opinion in Neurobiology 17(4):423–29. Retrieved (http://eutils.ncbi.nlm.nih.gov/entrez/eutils/elink.fcgi?dbfrom=pubmed&id=17714934&retmode=ref&cmd=prlinks).

Warren D. H. and W. T. Cleave. 1971. “Visual-Proprioceptive Interaction under Large Amounts of Conflict.” Journal of experimental psychology 90(2):206–14. Retrieved (http://eutils.ncbi.nlm.nih.gov/entrez/eutils/elink.fcgi?dbfrom=pubmed&id=5134326&retmode=ref&cmd=prlinks).

Watson Charles S., Steven C. Kellogg, David T. Kawanishi, and Patrick A. Lucas. 1973. “The Uncertain Response in Detection-Oriented Psychophysics.” Journal of Experimental Psychology 99(2):180–85.

Webster, Michael A. 2015. “Visual Adaptation.” Annual Review of Vision Science 1:547–67. Retrieved (http://eutils.ncbi.nlm.nih.gov/entrez/eutils/elink.fcgi?dbfrom=pubmed&id=26858985&retmode=ref&cmd=prlinks).

Webster Michael A., Daniel Kaping, Yoko Mizokami, and Paul Duhamel. 2004. “Adaptation to Natural Facial Categories.” Nature 428(6982):557–61. Retrieved (http://eutils.ncbi.nlm.nih.gov/entrez/eutils/elink.fcgi?dbfrom=pubmed&id=15058304&retmode=ref&cmd=prlinks).

Webster, Michael A. and Donald I. A. MacLeod. 2011. “Visual Adaptation and Face Perception.” Philosophical Transactions of the Royal Society B: Biological Sciences 366(1571):1702–25. Retrieved (http://eutils.ncbi.nlm.nih.gov/entrez/eutils/elink.fcgi?dbfrom=pubmed&id=21536555&retmode=ref&cmd=prlinks).

Wei, Kunlin and Konrad P. Körding. 2011. “Causal Inference in Sensorimotor Learning and Control.” Pp. 30–45 in Sensory Cue Integration, edited by J. Trommershäuser, K. Kording, and M. S. Land. New York.

Wei, Xue-Xin and Alan A. Stocker. 2012. “Efficient Coding Provides a Direct Link between Prior and Likelihood in Perceptual Bayesian Inference.” Advances in Neural Information Processing Systems 25 1313–21.

Wei, Xue-Xin and Alan A. Stocker. 2015. “A Bayesian Observer Model Constrained by Efficient Coding Can Explain ‘Anti-Bayesian’ Percepts.” Nature Neuroscience 18(10):1509–17. Retrieved September 7, 2015 (http://dx.doi.org/10.1038/nn.4105).

Weil, Leonora G. et al. 2013. “The Development of Metacognitive Ability in Adolescence.” Consciousness and cognition 22(1):264–71. Retrieved February 13, 2014 (http://www.pubmedcentral.nih.gov/articlerender.fcgi?artid=3719211&tool=pmcentrez&rendertype=abstract).

Weiskrantz L. 1996. “Blindsight Revisited.” Current opinion in neurobiology 6(2):215–20. Retrieved September 15, 2015 (http://www.ncbi.nlm.nih.gov/pubmed/8725963).

Weiss, Yair, Eero P. Simoncelli, and Edward H. Adelson. 2002. “Motion Illusions as Optimal Percepts.” Nature Neuroscience 5(6):598–604. Retrieved (http://www.nature.com/neuro/journal/v5/n6/full/nn858.html).

Van Wert, Michael J., Todd S. Horowitz, and Jeremy M. Wolfe. 2009. “Even in Correctable Search, Some Types of Rare Targets Are Frequently Missed.” Attention, perception & psychophysics 71(3):541–53. Retrieved September 16, 2015 (http://www.pubmedcentral.nih.gov/articlerender.fcgi?artid=2701252&tool=pmcentrez&rendertype=abstract).

Whiteley, Louise and Maneesh Sahani. 2008. “Implicit Knowledge of Visual Uncertainty Guides Decisions with Asymmetric Outcomes.” Journal of vision 8(3):2.1-15. Retrieved November 26, 2014 (http://www.journalofvision.org/content/8/3/2).

Whiteley, Louise and Maneesh Sahani. 2012. “Attention in a Bayesian Framework.” Frontiers in Human Neuroscience 6(June):100. Retrieved (http://eutils.ncbi.nlm.nih.gov/entrez/eutils/elink.fcgi?dbfrom=pubmed&id=22712010&retmode=ref&cmd=prlinks%5Cnpapers3://publication/doi/10.3389/fnhum.2012.00100).

Wilimzig, Claudia, Naotsugu Tsuchiya, Manfred Fahle, Wolfgang Einhäuser, and Christof Koch. 2008. “Spatial Attention Increases Performance but Not Subjective Confidence in a Discrimination Task.” Journal of vision 8(5):1–10. Retrieved November 22, 2013 (http://www.ncbi.nlm.nih.gov/pubmed/18842078).

Winman, Anders and Peter Juslin. 1993. “Calibration of Sensory and Cognitive Judgments: Two Different Accounts.” Scandinavian Journal of Psychology 34(2):135–48. Retrieved March 24, 2016 (http://doi.wiley.com/10.1111/j.1467-9450.1993.tb01109.x).

von Winterfeldt, Detlof and Ward Edwards. 1982. “Costs and Payoffs in Perceptual Research.” Psychological Bulletin 91(3):609–22.

Witt, Jessica K. 2011. “Action’s Effect on Perception.” Current Directions in Psychological Science. Retrieved (http://cdp.sagepub.com/content/20/3/201.short).

Witt Jessica K., Dennis R. Proffitt, and William Epstein. 2005. “Tool Use Affects Perceived Distance, but Only When You Intend to Use It.” Journal of Experimental Psychology: Human Perception and Performance 31(5):880–88. Retrieved (http://eutils.ncbi.nlm.nih.gov/entrez/eutils/elink.fcgi?dbfrom=pubmed&id=16262485&retmode=ref&cmd=prlinks).

Wohlgemuth A. 1911. On the After-Effect of Seen Movement. Cambridge University Press. Retrieved (https://books.google.com/books?id=Z6AhAQAAIAAJ).

Wolfe, Jeremy M. et al. 2007. “Low Target Prevalence Is a Stubborn Source of Errors in Visual Search Tasks.” Journal of experimental psychology. General 136(4):623–38. Retrieved December 10, 2014 (http://www.pubmedcentral.nih.gov/articlerender.fcgi?artid=2662480&tool=pmcentrez&rendertype=abstract).

Wolfe Jeremy M., David N. Brunelli, Joshua Rubinstein, and Todd S. Horowitz. 2013. “Prevalence Effects in Newly Trained Airport Checkpoint Screeners: Trained Observers Miss Rare Targets, Too.” Journal of vision 13(3):33. Retrieved December 29, 2014 (http://www.pubmedcentral.nih.gov/articlerender.fcgi?artid=3848386&tool=pmcentrez&rendertype=abstract).

Wolfe Jeremy M., Todd S. Horowitz, and Naomi M. Kenner. 2005. “Cognitive Psychology: Rare Items Often Missed in Visual Searches.” Nature 435(7041):439–40. Retrieved December 28, 2014 (http://dx.doi.org/10.1038/435439a).

Wolfe, Jeremy M. and Michael J. Van Wert. 2010. “Varying Target Prevalence Reveals Two Dissociable Decision Criteria in Visual Search.” Current biology 20(2):121–24. Retrieved December 4, 2014 (http://www.pubmedcentral.nih.gov/articlerender.fcgi?artid=2818748&tool=pmcentrez&rendertype=abstract).

Wyart, Valentin and Etienne Koechlin. 2016. “Choice Variability and Suboptimality in Uncertain Environments.” Current Opinion in Behavioral Sciences 11:109–15. Retrieved (http://dx.doi.org/10.1016/j.cobeha.2016.07.003).

Wyart, Valentin, Nicholas E. Myers, and Christopher Summerfield. 2015. “Neural Mechanisms of Human Perceptual Choice under Focused and Divided Attention.” The Journal of neuroscience 35(8):3485–98. Retrieved February 27, 2015 (http://www.jneurosci.org/content/35/8/3485.abstract?etoc).

Yamins, Daniel L. K. et al. 2014. “Performance-Optimized Hierarchical Models Predict Neural Responses in Higher Visual Cortex.” Proceedings of the National Academy of Sciences of the United States of America 111(23):8619–24. Retrieved August 28, 2017 (http://www.ncbi.nlm.nih.gov/pubmed/24812127).

Yeshurun, Yaffa and Marisa Carrasco. 1998. “Attention Improves or Impairs Visual Performance by Enhancing Spatial Resolution.” Nature 396(6706):72–75. Retrieved (http://eutils.ncbi.nlm.nih.gov/entrez/eutils/elink.fcgi?dbfrom=pubmed&id=9817201&retmode=ref&cmd=prlinks).

Yeshurun, Yaffa, Marisa Carrasco, and Laurence T. Maloney. 2008. “Bias and Sensitivity in Two-Interval Forced Choice Procedures: Tests of the Difference Model.” Vision Research 48(17):1837–51. Retrieved September 30, 2015 (http://www.sciencedirect.com/science/article/pii/S0042698908002599).

Yeung, Nick and Christopher Summerfield. 2012. “Metacognition in Human Decision-Making: Confidence and Error Monitoring.” Philosophical transactions of the Royal Society of London. Series B, Biological sciences 367(1594):1310–21. Retrieved March 2, 2013 (http://www.pubmedcentral.nih.gov/articlerender.fcgi?artid=3318764&tool=pmcentrez&rendertype=abstract).

Yu, Angela J. and Jonathan D. Cohen. 2009. “Sequential Effects: Superstition or Rational Behavior?” Advances in Neural Information Processing Systems 21:1873–80.

Zacksenhouse M., R. Bogacz, and P. Holme. 2010. “Robust versus Optimal Strategies for Two-Alternative Forced Choice Tasks.” Journal of mathematical psychology 54(2):230–46. Retrieved September 16, 2015 (http://www.pubmedcentral.nih.gov/articlerender.fcgi?artid=3505075&tool=pmcentrez&rendertype=abstract).

Zak, Ido, Mikhail Katkov, Andrei Gorea, and Dov Sagi. 2012. “Decision Criteria in Dual Discrimination Tasks Estimated Using External-Noise Methods.” Attention, perception & psychophysics 74(5):1042–55. Retrieved September 23, 2014 (http://www.ncbi.nlm.nih.gov/pubmed/22351481).

Zamboni, Elisa, Timothy Ledgeway, Paul V. McGraw, and Denis Schluppeck. 2016. “Do Perceptual Biases Emerge Early or Late in Visual Processing? Decision-Biases in Motion Perception.” Proceedings of the Royal Society of London B: Biological Sciences 283(1833). Retrieved June 6, 2017 (http://rspb.royalsocietypublishing.org/content/283/1833/20160263).

Zhang, Hang and Laurence T. Maloney. 2012. “Ubiquitous Log Odds: A Common Representation of Probability and Frequency Distortion in Perception, Action, and Cognition.” Frontiers in neuroscience 6:1. Retrieved September 5, 2015 (http://www.pubmedcentral.nih.gov/articlerender.fcgi?artid=3261445&tool=pmcentrez&rendertype=abstract).

Zhang, Hang, Camille Morvan, and Laurence T. Maloney. 2010. “Gambling in the Visual Periphery: A Conjoint-Measurement Analysis of Human Ability to Judge Visual Uncertainty.” PLoS computational biology 6(12):e1001023. Retrieved April 6, 2016 (http://dx.plos.org/10.1371/journal.pcbi.1001023).

Zylberberg, Ariel, Pablo Barttfeld, and Mariano Sigman. 2012. “The Construction of Confidence in a Perceptual Decision.” Frontiers in integrative neuroscience 6(September):79. Retrieved April 11, 2013 (http://www.pubmedcentral.nih.gov/articlerender.fcgi?artid=3448113&tool=pmcentrez&rendertype=abstract).

Zylberberg, Ariel, Pieter R. Roelfsema, and Mariano Sigman. 2014. “Variance Misperception Explains Illusions of Confidence in Simple Perceptual Decisions.” Consciousness and cognition 27:246–53. Retrieved June 23, 2014 (http://www.sciencedirect.com/science/article/pii/S1053810014000865).

